# Expanding the genetic toolbox for the obligate human pathogen *Streptococcus pyogenes*

**DOI:** 10.1101/2024.03.04.582890

**Authors:** Nina Lautenschläger, Katja Schmidt, Carolin Schiffer, Thomas F. Wulff, Karin Hahnke, Knut Finstermeier, Moïse Mansour, Alexander K. W. Elsholz, Emmanuelle Charpentier

## Abstract

Genetic tools form the basis for the study of molecular mechanisms. Despite many recent advances in the field of genetic engineering in bacteria, genetic toolsets remain scarce for non-model organisms, such as the obligatory human pathogen *Streptococcus pyogenes*.

In this study, we set out to develop a comprehensive set of plasmids, promoters and reporters for *S. pyogenes*. We present an expansion to the current genetic toolbox that comprises new replicative and site-specific integrative plasmids. Moreover, we established a collection of constitutive promoters with a wide variety of strengths as well as a set of novel inducible regulatory elements, including a zinc-inducible promoter, an erythromycin-inducible riboswitch and an IPTG-inducible promoter that outperform previously described inducible systems in terms of tightness and inducibility. In addition, we demonstrated the applicability of two codon-optimized fluorescent proteins, mNeongreen and mKate2, as reporters in *S. pyogenes*. For this, we adapted a novel chemically defined medium called RPMI4Spy. This medium showed a highly reduced autofluorescence compared to other growth media and allowed efficient signal detection in plate reader assays and fluorescence microscopy. Finally, we developed a plasmid-based system for genome engineering in *S. pyogenes* featuring the counterselection marker *pheS*\*, which improved the generation of scarless gene deletions.

This new toolbox simplifies previously laborious genetic manipulation procedures and lays the foundation for new methodologies to study gene functions in *S. pyogenes,* leading to a better understanding of its virulence mechanisms and physiology.

## 1 Introduction

Studying the function of a gene of interest requires appropriate tools for modifying, deleting or inserting genetic information. To do so, plasmid-based systems have been developed that aim to introduce point mutations, scarless deletions or allelic replacements (1). Various native or synthetic promoters can be used to modify the expression levels of a gene of interest to identify potential toxic or dose-dependent effects (2–4). Beyond that, different enzyme-based and fluorescent reporters have been established for investigating gene expression profiles, localizing specific proteins within a cell, determining protein-protein interactions, tracking bacteria during infection or studying microbial communities (5–7).

Genetic toolboxes have mainly been designed for highly-studied model organisms, such as *Bacillus subtilis*, or organisms useful for biotechnological purposes like Cyanobacteria (1). However, for many pathogenic bacteria, the genetic toolkit remains limited, despite their clinical relevance. One example is the obligatory human pathogen *Streptococcus pyogenes*, also known as Group A Streptococcus (GAS). Worldwide, GAS is responsible for over 700 million infections and around half a million deaths annually (8,9). Although there is extensive knowledge on many of the encoded virulence factors, a large percentage of the ∼1,800 genes remains to be studied (10). A better understanding of the physiological role of these genes could lead to the identification of potential new targets for drug and vaccine development.

Despite the availability of a number of plasmid systems for protein expression or genetic engineering in GAS, most current systems were developed decades ago and, since then, have rarely been optimized, updated or expanded (10). To edit the genome of *S. pyogenes*, researchers have previously used methodologies based on single- and double-crossover homologous recombination (HR), facilitated by thermosensitive plasmids or the Cre-*lox* system, respectively (10–12). However, plasmid curing makes these approaches rather laborious. A faster approach uses so-called “suicide” plasmids that are unable to replicate in the target strains and insert into specific sites in the genome via allelic replacement based on HR, also introducing an antibiotic marker for selection (13,14). To date, only one target-specific integrative plasmid has been described for *S. pyogenes*. This plasmid, pFW11e, inserts into the *SPy_0535* locus (*S. pyogenes* strain SF370) encoding a sugar-phosphate isomerase (15).

Integrases derived from bacteriophages have also been used for site-specific insertion of DNA fragments into the bacterial genome. One example is the plasmid p7INT, which encodes the integrase and respective *attP* site from the *S. pyogenes* temperate phage T12 (16). This integrase recognizes the *attB* site located at the 3’ end of the transfer-messenger RNA (tmRNA) gene and performs a precise insertion without impairing the functionality (16). However, the application of this plasmid is limited to only a few *S. pyogenes* strains (e.g. NZ131, SF370, and HSC5) possessing a matching *attB* site, although a single-base mutation in the *attP* site has been shown to expand the host range (10).

The establishment of more advanced genetic tools requires suitable genomic integration sites and inducible promoters. As a large part of the streptococcal genome is still uncharacterized, the identification of sites to insert genes of interest, without disrupting important genes and regulatory sequences, is challenging. Furthermore, only two inducible promoters are commonly used in GAS, both of which have certain drawbacks. One such system is the nisin-controlled gene expression (NICE) system, derived from a cluster of genes involved in nisin production and conferring immunity against it in *Lactococcus lactis* (17,18). Although the NICE system is functional in *S. pyogenes*, the sublethal concentrations of nisin required do not result in strong induction, and background expression is high (19). The most commonly used inducible expression system is based on the P*_tet_* promoter, driving transcription of a gene of interest in the presence of (anhydro-)tetracycline. A version of P*_tet_* containing three operator sites (P*_tet_*(O)_3_) has been constructed for inducible gene expression in GAS, but has shown limitations in its dynamic range (20). Hence, it is first necessary to develop new, optimized systems for tunable gene expression and to determine suitable integration sites, before more advanced genetic engineering approaches can be implemented.

Reporter genes are essential for studying the expression of a gene or the sub-cellular localization of a protein of interest. The panel of reporter genes available for GAS is small, with a primary focus on the firefly luciferase (*ffluc*) or the β-glucuronidase (*gusA*) for assessing promoter activities (5,15,21,22). Only few studies report the use of fluorescent proteins (FPs), such as GFP or mCherry in *S. pyogenes* (7,23–25). Previous studies have argued that streptococcal growth conditions, such as low oxygen supply and acidification of the growth medium by lactic acid production negatively impact GFP function (24,26). However, new generations of FPs with improved properties, e.g. higher stability at low pH, increased brightness and accelerated folding times, remain to be analyzed in GAS (27,28).

In this study, we compiled a comprehensive genetic toolbox for *S. pyogenes* to enable the implementation of more advanced genetic engineering technologies in GAS. We developed a set of replicative and integrative plasmids that enable sub-cloning procedures in *Escherichia coli* and the expression of proteins of interest in *S. pyogenes*. We compared the activity of various constitutive promoters to determine the variability in promoter strengths and characterized novel inducible regulatory elements allowing tunable gene expression. Furthermore, two codon-optimized fluorescent reporters, mNeongreen and mKate2, were tested for their functionality in *S. pyogenes*. For this, a chemically defined medium has been adapted and evaluated that enables fluorescence measurements in plate reader and microscopy setups. Finally, we designed and tested a new plasmid system featuring the counterselection marker *pheS\** for the implementation of scarless gene deletions in *S. pyogenes*.

## 2 Materials and Methods

### 2.1 Bacterial strains and growth conditions

*E. coli* DH5*α*, One Shot^TM^ TOP10 or BW25113 (for reporter plasmids containing P*_lac_*_(*Spn*)_ and derivatives) were grown in Luria-Bertani (LB medium) at 37°C with aeration (180 r.p.m.) or on LB agar plates at 37°C (Supplementary Table 1). Carbenicillin (100 μg/mL), chloramphenicol (20 μg/mL), kanamycin (50 μg/mL) or erythromycin (300 μg/mL) were used for selection (Supplementary Table 1). Transformation of chemically competent *E. coli* was performed using a heat shock-based protocol as described previously (29). *S. pyogenes* SF370 (serotype M1) and derivatives (Supplementary Table 1) were grown in Todd-Hewitt Broth with 2 g/L Yeast extract (THY medium), in GAS chemically defined medium (GAS CDM Powder from Alpha BioSciences supplemented with 1% w/v glucose, ferric nitrate nonahydrate (50 mg/L), ferrous sulfate heptahydrate (5 mg/L), manganese sulfate monohydrate (3.4 mg/L), NaHCO_3_ (2.5 g/L) and L-cysteine (708 mg/L)) (30), or on agar plates such as Trypticase Soy Agar (TSA) plates containing 5% sheep blood or THY agar plates at 37°C and 5% atmospheric CO_2_. For fluorescence measurements, *S. pyogenes* was cultured in RPMI4Spy medium [Gibco^TM^ RPMI1640 medium with glutamine and without phenol red, 1x Nucleobase mix (20 mg/L Guanine, 20 mg/L Adenine and 20 mg/L Uracil), 1x BME Vitamin solution, 1x RPMI Amino Acids solution, 7 g/L Glucose, 0.02 mM HEPES buffer (pH 7.4), 10 mg/L Niacinamide]. *S. pyogenes* electrocompetent cells were prepared and transformed as previously described using BioRad 1 mm cuvettes and electroporation at 400 Ω, 25 µF and 1800 V (31,32). Antibiotics were added at the following concentrations when necessary: chloramphenicol (20 μg/mL and 5 µg/mL for plasmids from pSpy2 series), kanamycin (300 μg/mL), erythromycin (3 μg/mL).

### 2.2 Extraction of genomic DNA

Genomic DNA of *S. pyogenes* and *B. subtilis* PY79 was prepared using the NucleoSpin Microbial DNA Kit (Macherey Nagel) following the manufacturer’s protocol. The lysis step was performed using the FastPrep-24 5G from MP Biomedicals (*S. pyogenes* program 20 seconds, 2x).

### 2.3 Plasmid construction

All PCR reactions were conducted using either Q5 polymerase or Phusion^TM^ High Fidelity Polymerase (Thermo Scientific) except for colony PCRs, which were performed using OneTaq^®^ Polymerase (NEB). For plasmid assembly, we followed either the Golden Gate Assembly or Gibson Assembly^®^ strategy (33,34). After transformation into *E. coli*, plasmid sequences were validated using Sanger sequencing (Microsynth Seqlab GmbH). All primers, plasmids and the DNA sequences of the most important genes and promoters from this study can be found in in Supplementary Tables 2, 3 and 4, respectively. All newly constructed backbones and plasmids containing relevant inserts used in this study will be made available through Addgene.

#### 2.3.1 Replicative plasmid backbones

All replicative plasmid backbones were constructed using the Golden Gate Assembly strategy with the type II restriction enzyme *Esp*3I (NEB) according to the manufacturer’s protocol (Table 1). The plasmid backbones were designed based on the shuttle plasmids pIB166, pIB184-Km and pIB185 as well as pBAV1K-T5-*gfp* (35–37). Plasmids pIB166 (Addgene plasmid # 90189), pIB184-Km (Addgene plasmid # 90195) and pIB185 (Addgene plasmid # 90196) were donated by Indranil Biswas. pBAV1K-T5-gfp was a gift from Ichiro Matsumura (Addgene plasmid # 26702).

**Table 1.** Replicative and integrative plasmid backbones developed in this study. MCS = multiple cloning site; Cm^R^ = chloramphenicol resistance; Kan^R^ = kanamycin resistance, Erm^R^ = erythromycin resistance

| Plasmid | Type | Size in bp | Features |
| --- | --- | --- | --- |
| pSpy1C | replicative | 4831 | pSH71 replicon, P<sub>lac(Eco)</sub>_mrfp in MCS, Cm<sup>R</sup> |
| pSpy1K | replicative | 4658 | pSH71 replicon, P<sub>lac(Eco)</sub>_mrfp in MCS, Kan<sup>R</sup> |
| pSpy1E | replicative | 4766 | pSH71 replicon, P<sub>lac(Eco)</sub>_mrfp in MCS, Erm<sup>R</sup> |
| pSpy2C | replicative | 7196 | pAMβ1 replicon, pUC origin, P<sub>lac(Eco)</sub>_mrfp in MCS, Cm<sup>R</sup> |
| pSpy2E | replicative | 7093 | pAMβ1 replicon, pUC origin, P<sub>lac(Eco)</sub>_mrfp in MCS, Erm<sup>R</sup> |
| pSpy3C | replicative | 4047 | pWV01* replicon, P<sub>lac(Eco)</sub>_mrfp in MCS, Cm<sup>R</sup> |
| pSpy3K | replicative | 3741 | pWV01* replicon, P<sub>lac(Eco)</sub>_mrfp in MCS, Kan<sup>R</sup> |
| pSpy3E | replicative | 3978 | pWV01* replicon, P<sub>lac(Eco)</sub>_mrfp in MCS, Erm<sup>R</sup> |
| pSpy0C2 | integrative | 5602 | pUC origin, P<sub>lac(Eco)</sub>_mrfp in MCS, homology arms for integration into *amyA*, Cm<sup>R</sup> |
| pSpy0C4 | integrative | 5237 | pUC origin, P<sub>lac(Eco)</sub>_mrfp in MCS, homology arms for integration into *sagB*, Cm<sup>R</sup> |
| pSpy0K6 | integrative | 5344 | pUC origin, P<sub>lac(Eco)</sub>_mrfp in MCS, homology arms for integration into *SPy_1078*, Kan<sup>R</sup> |
| p7INT.1 | integrative | 5035 | pUC origin, P<sub>lac(Eco)</sub>_mrfp in MCS, T12 integrase and *attP* site, Erm<sup>R</sup> |
| pERASE | integrative | 4457 | pUC origin, P<sub>lac(Eco)</sub> _mrfp in MCS, *pheS* * counterselection marker under control of P<sub>veg</sub>, Erm<sup>R</sup> |

First, all *Esp*3I recognition sites present in the backbones were removed by PCR mutagenesis using primer pairs OLEC9801/9802, OLEC9799/9800, OLEC9875/9876 and OLEC9831/9832, respectively. To construct pSpy1C (based on pIB166) and pSpy3K (based on pBAV1K-T5-*gfp*), the previously mutated plasmid backbones were amplified with the primer pairs OLEC9773/9774 and OLEC9829/9830, respectively. The multiple cloning site (MCS), which is separated by a reporter cassette encoding the *mrfp* gene controlled by the P*_lac_*_(*Eco*)_ promoter, was amplified from pBS3C*lux* (13) using OLEC9781/9782. pBS3Clux was a gift from Thorsten Mascher (Addgene plasmid # 55172). Next, restriction enzyme recognition sites for the MCS were attached to the amplified P*_lac_*_(*Eco*)__*mrfp* fragment by PCR using OLEC9775/9776. The final MCS was then inserted into the backbones using Golden Gate Assembly.

For the construction of pSpy1K and pSpy1E, the pSpy1C backbone was amplified using OLEC9826/9776 to remove the chloramphenicol resistance cassette. The kanamycin and erythromycin resistance cassettes were amplified using OLEC9824/9825 and OLEC9827/9828 from pIB184-Km and pIB185, respectively. Golden Gate Assembly was performed to introduce the marker genes into the backbones.

To construct pSpy2K, we used OLEC9777/9778 to amplify the replicon and resistance cassette from pIB184-Km (*Esp*3I removed). For pSpy2E, the replicon and resistance cassette were amplified by PCR using OLEC9779/9780 from pIB185. We then cloned the previously amplified MCS into the backbones using Golden Gate Assembly. To create pSpy2C, the plasmid backbone of pSpy2K was amplified by PCR using the primers OLEC9877/9878. The *cat* fragment to be introduced was amplified with OLEC9835/9836 from pIB166. Then, the fragments were joined using Golden Gate Assembly. As no colonies were obtained after transformation of the pSpy2 series, we hypothesized that *S. pyogenes* was not transformable with these plasmids, and exchanged the pIB185-derived replicon with the pAMβ1 replicon from pMSP3535. Thus, we used OLEC14365/14366 to amplify all plasmid parts except the pAMβ1 replicon of pSpy2C (not functional) and OLEC14367/14368 to amplify the replicon from a derivative of pMSP3535 (38). The fragments were joined by Gibson Assembly^®^ to create pSpy2C. For pSpy2E, we amplified part of the backbone using OLEC14365/14369 from pSpy2E (not functional) and used OLEC14370/14368 to obtain the new pAMβ1 replicon from the pMSP3535 derivative. The fragments were again joined by Gibson Assembly^®.^ We were unable to obtain a correct assembly of the repaired pSpy2K plasmid.

For pSpy3C and pSpy3E, the plasmid backbone of pSpy3K was amplified with primers OLEC9833/9834. Overhangs for cloning were attached to both resistance cassettes, *cat* and *ermB*, using the oligo pairs OLEC9835/9836 and OLEC9837/9838, respectively. The previously amplified cassettes used for pSpy1C and pSpy1E served as templates. The plasmids were assembled using Golden Gate Assembly.

#### 2.3.2 Luciferase reporter plasmids

For the construction of the reporter plasmids harboring the *ffluc* reporter gene (*Photinus pyralis*), we used either the Golden Gate Assembly strategy using *Eco*31I (Thermo Scientific™) or Gibson Assembly^®^.

To construct pSpy1C-*ffluc* (control plasmid), the *ffluc* gene was PCR-amplified using the oligos OLEC10182/10028 from pLZ12Km2-P*23*R:TA:*ffluc,* a gift from Thomas Proft (Addgene plasmid # 88900) (22), and assembled into the linearized pSpy1C plasmid. To assess the activity of different constitutive promoters, the promoters P*_23_* (*L. lactis*), P*_veg_* (*B. subtilis*), P*_gyrA_*_(*Spy*)_ (*S. pyogenes*) and P*_xylS2_* (synthetic promoter) were cloned upstream of *ffluc* into pSpy1C as detailed in Supplementary Table 5. Each inducible promoter or regulatory element was placed upstream of *ffluc* in pSpy1C as described in Supplementary Table 6. For some of the PCRs, we used the following plasmids as template: pEU8517 (39), pMSP3535 donated by Gary Dunny (Addgene plasmid # 46886) (38), pPEPY-PF6-lacI (Addgene plasmid # 85589) and pJWV102-PL-dCas9 (Addgene plasmid # 85588) donated by Jan-Willem Veening (40).

To improve the signal-to-noise ratio during luciferase assays, an additional terminator was introduced upstream of all promoter-*ffluc* reporter fusions by PCR mutagenesis using OLEC11104/11105.

#### 2.3.3 Integrative plasmid backbones

We designed a set of integrative plasmids for use in *S. pyogenes* SF370 that integrate into different genomic loci as described in Table 1.

pSpy0C2 was cloned using Golden Gate Assembly using the type II restriction enzyme *Esp*3I (NEB). The MCS-*cat* fragment and the pUC origin were amplified using OLEC10311/10312 from pSpy1C and OLEC10315/10316 from pUC19-*mKate2∼ssrA*, respectively. Up- and downstream homologous regions of *amyA* were amplified from *S. pyogenes* genomic DNA using primers OLEC10930/10931 and OLEC10932/10933, respectively. All fragments were joined using Golden Gate Assembly. To construct pSpy0C4, the MCS-*cat* fragment and the pUC origin were amplified using OLEC11471/11472 from pSpy1C and OLEC11480/11476 from pUC19-*mKate2∼ssrA*, respectively. Up- and downstream homologous regions of the *sagB* gene were amplified from *S. pyogenes* genomic DNA using primers OLEC11473/11479 and OLEC11477/11535, respectively. Gibson Assembly^®^ was used to ligate all fragments. To clone pSpy0K6, the MCS was amplified from pSpy0C4 using OLEC14327/14328. The kanamycin resistance cassette was amplified from pLZ12Km2-P*23*R:TA:*ffluc* using OLEC14329/14330. Up- and downstream homology arms of *SPy_1078* were amplified from *S. pyogenes* genomic DNA with OLEC14331/14332 and OLEC14335/14336, respectively. The pUC origin was amplified from pUC19*-mKate2∼ssrA* using OLEC14333/14334. Up- and downstream homology arms and the origin were joined by LFH-PCR using OLEC14337/14335. The PCR product and the MCS-*cat* fragment were then ligated using Gibson Assembly^®^.

p7INT was a gift from Prof. Michael Federle (University of Illinois Chicago, USA) (16). To obtain p7INT.1 featuring the same MCS as the remaining plasmids of the collection, we amplified the p7INT backbone using OLEC14308/14309. The MCS was amplified using OLEC14306/14307 from pSpy1C and both fragments were joined using Gibson Assembly^®^. Next, we removed a restriction enzyme recognition site in the integrase open reading frame by PCR mutagenesis using OLEC14310/14311 to ensure that restriction sites of the MCS are only present once.

All plasmids, except p7INT and p7INT.1, were linearized using *Alw*44I prior to transformation into *S. pyogenes* SF370. After transformation, the integration of the plasmids was verified by PCR and the integration sites and distinct virulence genes (*ropB*, *csrRS*, *mga*) were sequenced using Sanger sequencing. In addition, whole genome sequencing of all strains harboring the integrative plasmids was performed using Nanopore sequencing.

#### 2.3.4 Reporter plasmids featuring mNeongreen and mKate2

The codon-optimized (41) reporters *mKate2* and *mNeongreen* (gene synthesis by GenScript) were assembled into p7INT and other backbones as described in Supplementary Table 7. For the cloning of *mNeongreen* and *mKate2* into pSpy0C4, inserts and backbone were digested with *Eco*RI and *Pst*I (Thermo Scientific^TM^). The fragments were ligated using T4 DNA Ligase (Thermo Scientific^TM^), creating pSpy0C4- *mNeongreen∼ssrA* and pSpy0C4-*mKate2∼ssrA*.

The DNA sequence coding for the last three amino acids of the native SsrA tag sequence downstream of the *mNeongreen* reporter gene was modified from LAA (WT) to LDD or ASV by PCR using the primer pairs OLEC13042/13043 and OLEC13040/13041, respectively. As control, a stop codon (*) was introduced upstream of the *ssrA* tag to prevent any SsrA-driven proteolytic degradation. The stop codon was introduced by PCR using OLEC13044/13045 for *mNeongreen* reporter constructs and OLEC13046/13047 for all *mKate2* reporter constructs.

#### 2.3.5 Construction of pERASE and deletion of *sagA* using pERASE-*ΔsagA*

Fragments for the assembly of pERASE, such as the pUC origin (from pUC19-*mKate2∼ssrA*), the erythromycin resistance cassette (from p7INT), the MCS (from pSpy0C4) and the *pheS*\* counterselection marker under control of P*_veg_* were amplified using primers OLEC14398/14523, OLEC14522/14403, OLEC14404/14405 and OLEC14524/14525, respectively. Fragments were assembled using Gibson Assembly^®^ and checked by Sanger sequencing. An optimal RBS was added upstream of *pheS\** using PCR mutagenesis with primers OLEC14726/14727. The nucleotide sequence for the counterselection marker *pheS\** encoding a variant of the phenylalanyl-tRNA-synthetase (*α*-subunit) was adapted to be less homologous to the native *pheS* locus in *S. pyogenes* to prevent recombination events at this site. The *pheS\** DNA sequence was obtained by gene synthesis (service provided by GenScript).

To construct pERASE-*ΔsagA* for the exemplary deletion of *sagA*, we amplified the pERASE backbone using primers OLEC14883/14884. Fragments up and- downstream of *sagA* were amplified from *S. pyogenes* genomic DNA with oligos OLEC14875/14876 and OLEC14877/14878, respectively. The three amplicons were assembled using Gibson Assembly^®^ and correct assembly was analyzed by restriction digestion and Sanger sequencing.

To delete *sagA*, pERASE-*ΔsagA* was transformed as described above into sucrose-competent *S. pyogenes* SF370 cells and integration events were selected on TSA blood agar plates with antibiotic selection at 37°C and 5% CO_2_. After 48 hours of growth, several clones were re-streaked onto selective TSA blood agar plates and incubated overnight. Then, four single colonies were inoculated into THY medium without antibiotic selection. Once the cultures reached the exponential growth phase (OD_620nm_ = 0.25), they were diluted 1:1000 in THY medium without antibiotics and incubated overnight at 37°C and 5% CO_2_. The following day, cultures were again diluted 1:1000 in THY medium and grown until the exponential growth phase. After three passages in liquid THY to enable curing of the plasmid, dilutions (10^-3^) were plated on THY agar plates with 5 mM PCPA (4-chlor-DL-phenylalanine). Selection on PCPA plates will result in death of all clones still harboring the integrated pERASE plasmid. The plates were incubated overnight at 37°C and 5% CO_2_. The next day, around 70 clones were picked and streaked onto TSA blood agar plates with or without antibiotic selection and on THY agar plates with PCPA and incubated for 48 hours. Loss of *sagA* will result in decreased hemolysis on TSA blood agar plates. Eight clones that demonstrated a decrease in hemolysis and three clones that did not were analyzed by PCR analysis using OLEC14875/14878. For three of the eight clones indicating loss of *sagA* in the PCR, we confirmed the deletion by Sanger sequencing.

### 2.4 Identification of transcriptionally silent sites

RNA reads were mapped to the respective reference sequences of *S. pyogenes* strains SF370 (NCBI assembly GCF_000006785.2) and 5448 (NCBI assembly GCF_900619585.1) using BWA mem2 V2.2 (42) and Samtools V1.15.1 (43), each with default settings. The coverage for each position and each RNA read file was extracted using the Python library Pysam V0.22.0 and its function pileup (43). Every position of each coverage dataset was normalized by dividing its coverage by the median of the upper quartile of the coverage values of the whole dataset. Values above 1 were set to 1. Next, for each position, the average across all strain-specific normalized coverages was calculated. Based on these strain-specific datasets, candidate regions were identified by masking prophage regions and annotated genes showing an average normalized expression of >0.02, including 250 bp upstream of every expressed gene. The remaining unmasked consecutive positions were categorized into unexpressed genes based on available strain annotation, unexpressed intergenic regions and true zero coverage regions. Unexpressed refers to a normalized average coverage of ≤0.02. All identified regions were further filtered for a minimum size of 100 bp (Supplementary Tables 8 and 9). Finally, candidate regions were identified in companion strains by mapping region sequences using BWA mem2 and Samtools, each with default settings and extracting companion sequences utilizing Python’s Pysam library as described above. Then, every pair of sequences was aligned to each other using Biopython’s pairwise2 global alignment module (V1.81) followed by reporting the Hamming distance normalized by sequence length (44). Pairs with a score <90%, arbitrarily chosen, were considered paralogs (Supplementary Table 10). The code is provided on GitHub (see data availability statement).

### 2.5 Nanopore sequencing for strain validation

Genomic DNA of strains harboring the integrative plasmids pSpy0C2, pSpy0C4, pSpy0K6 and p7INT.1 was extracted using the Monarch^®^ HMW DNA Extraction Kit for Tissue (New England Biolabs^®^) according to the manufacturers protocol for Gram-positive bacteria (low input) and sheared 15x with a G27 needle (B.Braun). Barcoded libraries were prepared using the Native Barcoding Kit (SQK-NBD114.24, Oxford Nanopore Technologies) according to the manufacturer’s protocol with 400 ng genomic DNA per sample, loaded on a MinION Flow Cell (R10.4.1, FLO-MIN114) and sequenced for 72 h on a MinION Mk1C (MinKNOW v23.04.8). POD5 files were basecalled and demultiplexed with the super accuracy model (v4.3.0) using Dorado (v0.5.0) with the “--barcode-both-ends” option to reduce false-positives during barcode classification.

We developed a Snakemake workflow for the identification of variants in bacterial genomes (see data availability statement) (45). In brief, reads were filtered using Filtlong (v0.2.1, available on GitHub: https://github.com/rrwick/Filtlong) with options “--keep_percent 90 --min_length 500 --min_mean_q 90” and aligned to the reference genome using NGMLR (v0.2.7) with default settings (46). Alignments files were sorted and indexed using Samtools (v1.9) and used for structural variants calling using Sniffles2 (v2.2) with standard settings and cuteSV (v2.1.0) with parameters suggested for ONT data (43,46–48). Single nucleotide variants were identified using Clair3 (v1.0.5) with the alignment generated by NGMLR and using Medaka (v1.11.3, available on GitHub: https://github.com/nanoporetech/medaka), which internally uses minimap2 for read alignment (46,49). Since we noticed putative assembly artefacts in the rRNA operons of the reference genome (data not shown), we additionally filtered the resulting variant files using VCFtools (v0.1.16) (50) to exclude rRNA-tRNA operon regions (± 10 bp): 17,055..29,081; 79,274..85,457; 264,339..269,670; 1,330,440..1,336,193; and 1,557,792..1,583,127 of NC_002737.2. NanoPlot (v1.42.0) and MultiQC (v1.19) were used for read quality control (51,52). Finally, identified variants were reported in a custom HTML report (RMarkdown v2.25) with alignments visualized for relevant regions using igv-reports (v1.10.0, available at GitHub: https://github.com/igvteam/igv-reports) (53).

As both cuteSV and Sniffles2 had difficulties to accurately identify allelic replacements, we provided the reference genome of *S. pyogenes* SF370 (NC_002737.2) as well as the sequence of the transformed integrative plasmid to facilitate the detection of allelic replacements as translocations. In addition, we re-performed the analysis using the expected genotype as reference.

### 2.6 Growth curves

*S. pyogenes* SF370 wildtype and derivatives were streaked on TSA blood agar plates with selective antibiotics if required. Alternatively, replicative plasmids were freshly transformed into *S. pyogenes* SF370 sucrose-competent cells. Strains or transformants were grown at 37°C and 5% CO_2_ overnight. The next day, overnight cultures were inoculated from single colonies in THY medium and incubated at 37°C with 5% CO_2_ overnight. Overnight cultures were harvested by centrifugation at 4,500 x g for 10 min and adjusted to an OD_620nm_ of 0.02 in THY medium. For maintenance of the replicative plasmids, corresponding selective antibiotics were supplemented at all times, while no antibiotics were applied for strains harboring integrative plasmids. To assess the effect of different chloramphenicol concentrations on the growth of the wildtype or *S. pyogenes* harboring pSpy1C, pSpy2C or pSpy3C, we added different chloramphenicol concentrations to the cultures (1 µg/mL, 2.5 µg/mL, 5 µg/mL, 10 mg/mL and 20 µg/mL) as well as 0.1% ethanol as control. Then, 200 µL of each sample were applied to a clear 96-well plate in technical triplicates together with the blank (medium only). Growth was measured every 20 min for 15 hours in a plate reader (Synergy H1, BioTek) at 37°C and 5% CO_2_. Experiments were performed in biological triplicates.

### 2.7 Luciferase assay to determine promoter activities

Plasmids were transformed into electrocompetent cells of the *S. pyogenes* SF370 wildtype or the LacI-expressing strain (SF370 *attB*(T12)::p7INT-P*_veg_*_*lacI*). Cultures were started as described for the growth curve experiments and incubated at 37°C with 5% CO_2_; OD_620nm_ and luminescence were measured every 30 min in a plate reader (Cytation 3, BioTek). To detect luminescence, 10 μL of a 1 mg/mL luciferin stock solution were added to the samples immediately before the measurement (integration time of 1s and read height of 1 mm, white 96-well plate).

To test the activity of the inducible promoters over the course of growth, the corresponding inducer compounds were added right at the start of the experiment (t_0_). The promoters P*_nisA_*, P*_tet_*, P*_Zn_*, P*_lac_*_(*Spn*),_ and P*_tre_* were induced with 0.75 μg/mL nisin, 10 ng/mL anhydrotetracycline (AHT), 80 μM ZnSO_4,_ 1 mM IPTG, and 0.4% trehalose, respectively. MilliQ water served as a control. For induction of the erythromycin-inducible riboswitch, we added 3 ng/mL erythromycin or 0.1% ethanol as control.

To assess the dose-response of the inducible promoters, day cultures of strains harboring the control plasmid (without promoter) or the reporter plasmid with the inducible promoter were set up and grown until the exponential growth phase (OD_620nm_ = 0.25). Then, the cultures were split and induced with a range of concentrations of the inducer compound as well as 0.1% ethanol or distilled water as control. Growth and luminescence were measured as described above every 30 minutes for three hours. All experiments were performed in biological triplicates.

### 2.8 Development of a new RPMI-based chemically defined medium for plate reader-based fluorescence measurements

To develop a new chemically defined medium (CDM) with less autofluorescence, we utilized the cell culture medium RPMI1640 with glutamine and without phenol red. We assessed the growth of *S. pyogenes* SF370 in different RPMI1640-based media compositions as described under ‘growth curves’ with minor adaptations. An additional washing step with 1x PBS solution was added before cell pellets were resuspended in the test media to remove any remaining THY medium. The initial recipes tested can be found in Supplementary Table 11.

We then compared the fluorescence signal of *S. pyogenes* cells expressing mNeongreen in the media RPMI4Spy, THY and CDM as follows. Reporter and control strains were streaked onto TSA blood agar plates and incubated at 37°C and 5% CO_2_ overnight. The following day, overnight cultures were prepared in THY medium and placed for incubation at 37°C and 5% CO_2_. Overnight cultures were harvested by centrifugation at 4,500 x g for 10 min, washed with sterile 1x PBS solution and adjusted to an OD_620nm_ of 0.02 in the different media. Cultures were distributed into a black 96-well plate (in duplicates) with clear bottom and incubated in a plate reader (Synergy H1, BioTek). Growth (OD_620nm_) and mNeongreen fluorescence (500 nm ex, 530 nm em) were measured every 20 minutes for 15 hours. For the comparison of mKate2 signals from different genomic loci, we proceeded as described above, but measured fluorescence using 588 nm ex, 633 nm em. Experiments were performed in biological triplicates.

### 2.9 Analyzing the influence of SsrA tag variants on mNeongreen stability

To analyze the effect of the SsrA tag variants on mNeongreen protein stability, we measured the decrease in fluorescence signal after inhibiting translation using spectinomycin. A growth curve experiment was set up as described previously with strains harboring the mNeongreen reporter fused to different SsrA tag variants as well as the control strain (no promoter) in RPMI4Spy medium (Supplementary Table 1). The strains were incubated in a plate reader, and growth (OD_620nm_) and fluorescence (500 nm ex, 530 nm em) were measured every 20 minutes until the exponential growth phase (OD_620nm_ = 0.2). Then, 100 μg/mL spectinomycin was applied to the samples to inhibit translation, and growth and fluorescence were measured for 10 more hours. Experiments were performed in biological triplicates.

### 2.10 Fluorescence microscopy

Overnight cultures of the mNeongreen and mKate2 reporter strains as well as the corresponding control strains (no promoter) (Supplementary Table 1) were grown as described before. Overnight cultures were used to prepare day cultures in RPMI4Spy as described above (final OD_620nm_ of 0.02). Cultures were grown at 37°C and 5% CO_2_ until the exponential growth phase (OD_620nm_ = 0.25). Then, 5 μL of each culture were mounted onto an agarose pad (0.75% agarose in 1x PBS) for fluorescence microscopy. Microscopy images were obtained using an inverted fluorescence microscope (Leica Dmi8, DFC9000 GT VSC-D6212 camera) using a 100x phase-contrast lens, and exposure times of 300 ms (mNeongreen, control) and 2 s (mKate2, control), respectively. Images were edited for publication using the Leica Application Suite X (LAS X) Software from Leica Microsystems. Experiments were performed in biological triplicates.

## 3 Results

### 3.1 Standardized replicative plasmids for *S. pyogenes*

We have designed and constructed a collection of plasmids for *S. pyogenes* that allows an easy transfer of inserts between backbones featuring different antibiotic markers and origins of replication with varying copy numbers. All plasmids share the same MCS containing an *mrfp* reporter gene controlled by the P*_lac_*_(*Eco*)_ promoter that is flanked by restriction enzyme recognition sites and M13 primer binding sites. In *E. coli*, the mRFP reporter enables screening for clones harboring the plasmid with the desired insert, which appear as white colonies, while clones harboring the empty plasmid appear red. The M13 primer binding sites can be used for colony PCR and subsequent sequencing of the assembled plasmids. We selected the replication origins pSH71, pAMβ1 and a previously engineered version of pWV01, pVW01*. While pSH71 and pWV01* are functional in both Gram-negative and Gram-positive bacteria, pAMβ1 is restricted to Gram-positive bacteria and requires the addition of a Gram-negative origin to allow for replication in *E. coli*. The replicons pSH71 and pWV01 have been reported as high-copy number replicons in several lactic acid bacteria, whereas pAMβ1 is commonly a low-copy replicon (54). For plasmid maintenance, we selected three antibiotic resistance cassettes (*cat*, *ermB* and *aph(3′)-IIIa*) that are functional in both *E. coli* and *S. pyogenes*.

The constructed pSpy1 plasmid series is based on the rolling-circle replicon (RCR) pSH71 and was combined with antibiotic resistance cassettes encoding for chloramphenicol (pSpy1C) (Figure 1A and Table 1), kanamycin (pSpy1K) and erythromycin (pSpy1E) (35). Based on the theta replicon pAMβ1 (10,55), we constructed the pSpy2 series comprising plasmids pSpy2C (Figure 1A and Table 1) and pSpy2E, featuring a chloramphenicol and erythromycin resistance cassette, respectively. Lastly, we designed the pSpy3 series using the RCR pWV01* (36), resulting in pSpy3C (Figure 1A and Table 1) (chloramphenicol resistance), pSpy3K (kanamycin resistance) and pSpy3E (erythromycin resistance). All replicative plasmids were transformed into *S. pyogenes* SF370 and analyzed for their effect on growth in THY medium with selection. Our results showed that bacteria harboring plasmids of the pSpy1 series grew very similar (Figure 1B). Interestingly, bacteria harboring pSpy2C from the pSpy2 series had a longer lag phase than cells harboring pSpy2E (Figure 1B). *S. pyogenes* cells harboring pSpy3C and pSpy3E from the pSpy3 series had very similar growth profiles, whereas bacteria harboring pSpy3K showed a longer lag phase (Figure 1B). As the variations in the growth profiles seemed to be based on the different antibiotic resistances, we hypothesized that optimizing the selective antibiotic concentration could improve the growth of the strains. To demonstrate this, we grew *S. pyogenes* harboring pSpy1C, pSpy2C and pSpy3C as well as the wildtype without any plasmid in the presence of different concentrations of chloramphenicol. Due to their higher copy number, the growth of *S. pyogenes* harboring pSpy1C and pSpy3C was less affected at higher concentrations of chloramphenicol, such as 10 µg/mL or 20 µg/mL (Supplementary Figure 1). In line with this, we observed faster growth of *S. pyogenes* harboring the low copy number plasmid pSpy2C only at concentrations of 5 µg/mL chloramphenicol, which is the minimum concentration required for growth inhibition of the wildtype (Supplementary Figure 1). Thus, optimization of the antibiotic concentration further improved the growth of strains containing the replicative plasmids.

**Figure 1:**
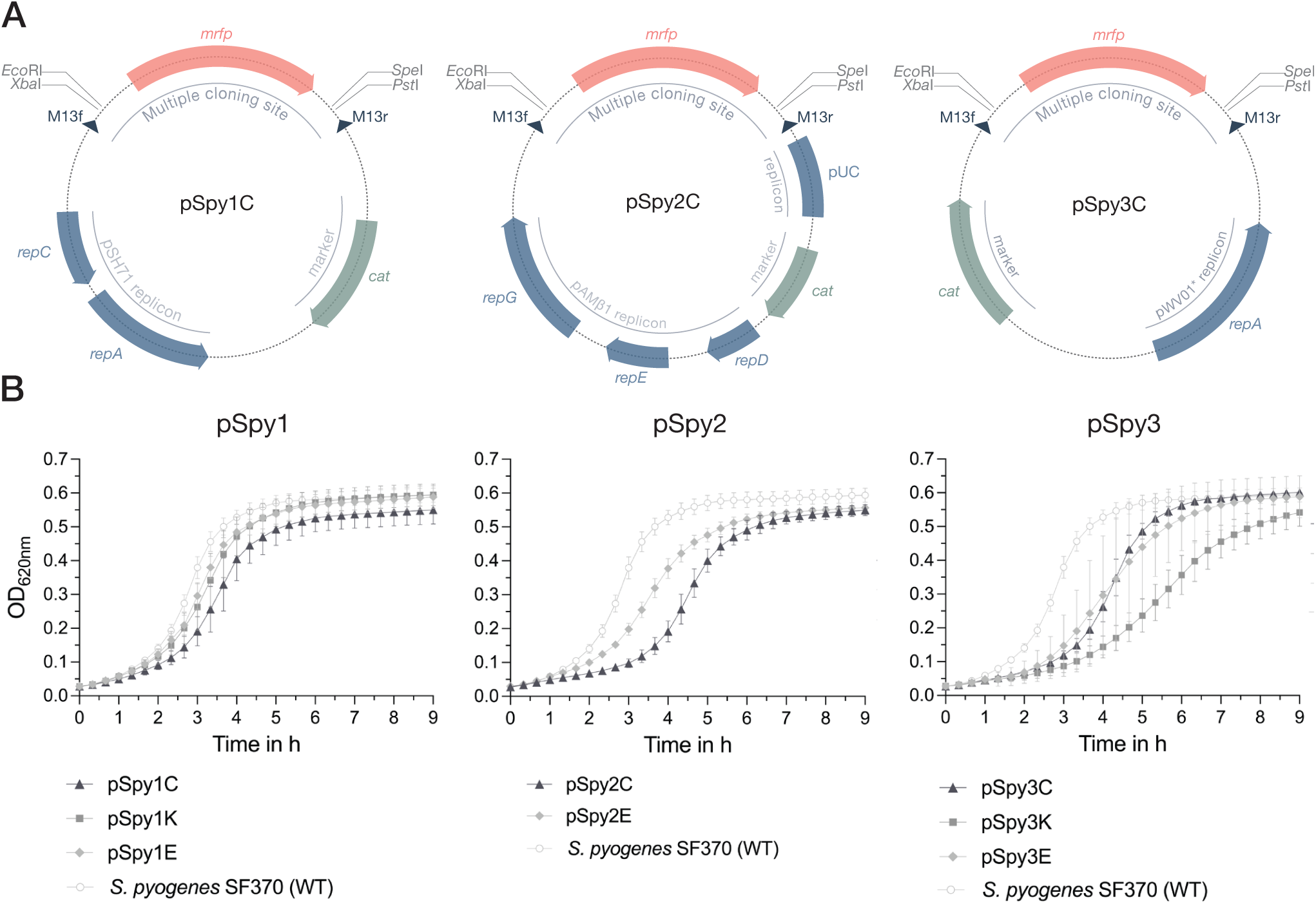
Novel replicative plasmids for use in *S. pyogenes*. **(A)** Representative schematic maps of the replicative plasmids pSpy1C (pSH71 replicon), pSpy2C (pAMβ1 replicon) and pSpy3C (engineered pWV01* replicon) for use in *S. pyogenes*. The origin of replication is indicated in blue, while the resistance cassette (*cat*) is highlighted in sage green. The specific restriction sites of the MCS flank the *mrfp* reporter gene, the expression of which is driven by P*_lac_*_(*Eco*)_, enabling red-white screening of single colonies during subcloning in *E. coli*. The dark blue triangles indicate the binding sites for the M13f and M13r standard primers. **(B)** Growth, shown as OD_620nm_, of *S. pyogenes* SF370 transformed with the different replicative plasmids in THY medium with antibiotic selection. Growth curves of cultures harboring pSpy1C (triangle), pSpy1K (square) and pSpy1E (diamond) are depicted on the left, while the graph in the center shows the growth of *S. pyogenes* transformed with pSpy2C (triangle) and pSpy2E (diamond). The graph on the right displays the growth of *S. pyogenes* harboring pSpy3C (triangle), pSpy3K (square) and pSpy3E (diamond). Growth of the *S. pyogenes* SF370 wildtype in THY medium without antibiotic selection is indicated using round data points with a white fill. Experiments were performed in biological triplicates and each measurement in technical duplicates.

### 3.2 Novel integrative plasmids targeting specific loci in the streptococcal genome

Previously, only two integrative plasmids have been described for use in GAS, pFW11e and p7INT (15,16). To enable the stable integration of multiple genes of interest into the streptococcal genome at different loci, we explored alternative integration sites. We first designed two integrative plasmids, pSpy0C2 and pSpy0C4, targeting the *amyA* and the *sagB* genes, respectively, for insertion by homologous recombination (Figure 2A and B, Table 1). Disruption of the *amyA* or *sagB* locus will enable positive integration events to be screened by selection on specific TSA plates. Whereas allelic replacement of *amyA* leads to inhibited starch degradation on starch plates, the deletion of *sagB* results in the loss of hemolysis on TSA blood plates.

**Figure 2.**
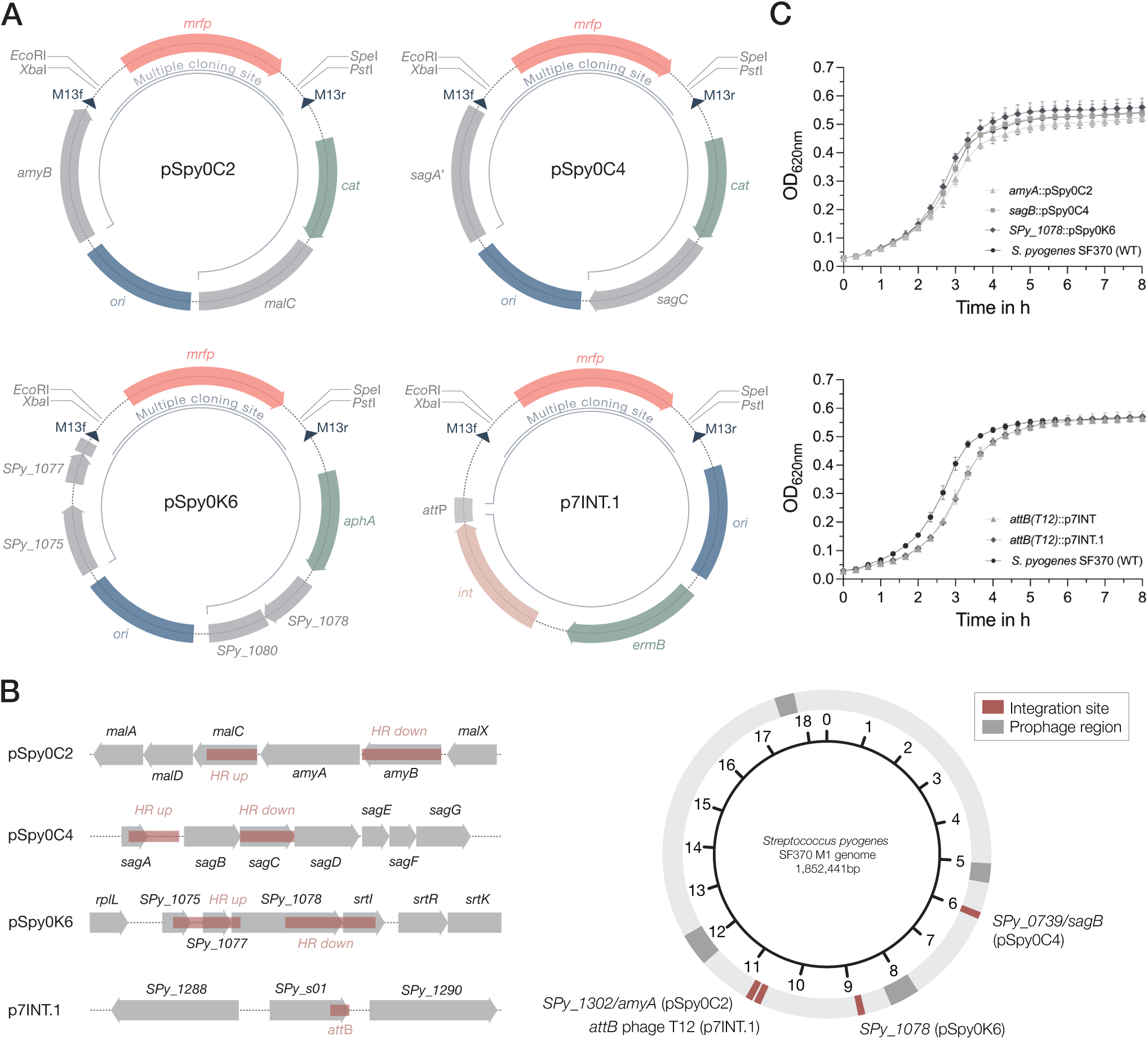
Plasmids for site-specific integration into the streptococcal genome. **(A)** Schematic plasmid maps of the three integrative plasmids designed for *S. pyogenes* SF370 and the integrase-based plasmid p7INT.1. The pUC replicon for propagation in *E. coli* is marked in blue. Resistance cassettes for chloramphenicol (*cat*), erythromycin (*ermB*) and kanamycin (*aph(3′)-IIIa*) are highlighted in sage green. DNA sequences with homology to the genome of SF370 are shown in grey. Restriction sites of the MCS flank the *mrfp* reporter gene under the control of P*_lac_*_(*Eco*)_, enabling red-white screening of clones containing the desired insert in *E. coli*. Dark blue triangles indicate the binding sites for the M13f and M13r standard primers. The inner black circular line marks the part of the plasmid that will integrate into the streptococcal genome. For the p7INT derivative p7INT.1, the pale pink color indicates the integrase gene (*int*) with the corresponding attachment site of prophage T12 (*att*P) marked in grey. **(B)** Left: Linear view of the genomic regions targeted by the integrative plasmids. Sites used for homologous recombination or insertion mediated by the integrase (*att*B) are marked in red. Genes are indicated in grey. Right: Circular map of the *S. pyogenes* SF370 genome indicating the localization of the different plasmid integration sites in red. Prophage regions are marked in dark grey. **(C)** Growth of *S. pyogenes* SF370 strains, shown as OD_620nm_, harboring different integrated plasmids in THY medium without antibiotics. The effect on growth of the integrative plasmids pSpy0C2 (triangle), pSpy0C4 (square) and pSpy0K6 (diamond) is shown in the top graph, while the bottom graph displays the growth of the strains harboring p7INT (triangle) and p7INT.1 (diamond) in the genome. An exemplary growth curve of the *S. pyogenes* SF370 wildtype is plotted (WT; black circles). Experiments were performed in biological triplicates and each measurement in technical duplicates.

We performed growth curve experiments to assess the effect of plasmid integration on growth. Strains harboring the novel integrative plasmids grew very similar compared to the *S. pyogenes* wildtype in THY medium without antibiotics, suggesting that these allelic replacements have no effect on streptococcal growth under this condition (Figure 2C). To propose alternative integration sites that do not disrupt transcribed genes, we screened the genome for transcriptionally silent sites suitable for integration, similar to the reported integrative vectors for *Streptococcus pneumoniae* (14). To do this, we used published RNA-seq datasets from *S. pyogenes* SF370 and the more clinically relevant M1T1 5448 strain grown under various conditions to identify transcriptionally silent regions (Supplementary Figures 2 and 3 and Supplementary Tables 8 and 9). We then analyzed, which of the obtained sites are present in both strains, and could therefore be targeted using the same plasmid (Supplementary Table 10). Among those sites, we identified the gene *SPy_1078* as a transcriptionally silent site in *S. pyogenes* SF370 (Supplementary Figures 2 and 3, Supplementary Table 8). This gene contains frameshift mutations resulting in several early stop codons. Based on this locus, we designed the plasmid pSpy0K6 featuring homologous regions to target *SPy_1078* for allelic replacement (Figure 2A and B, Table 1). Following successful integration of pSpy0K6, we conducted growth curve experiments. The strain harboring pSpy0K6 grew very similar to the wildtype in THY medium without antibiotics, indicating that disruption of *SPy_1078* had no effect on growth under the conditions tested (Figure 2C). In addition, we designed the integrative plasmid pSpy0C5 targeting the transcriptionally silent gene *SPy_1930* in *S. pyogenes* SF370, but were unable to obtain clones even with two different designs of the plasmid, potentially due to a stabilizing toxin-antitoxin locus just upstream (Supplementary Figure 4A and D).

To make the already available p7INT plasmid (Supplementary Figure 4B) compatible with our plasmid collection, we exchanged the MCS featuring *lacZ* to fit our MCS design, resulting in p7INT.1 (Figure 2A and Table 1). When growing the strains in THY medium without antibiotics, we noticed a growth delay of approximately 20 minutes for strains harboring p7INT or its derivative p7INT.1 compared to the wildtype (Figure 2C). This observation could depend on the growth conditions tested and is not caused by plasmid integration at the tmRNA locus as we have shown that SsrA-mediated proteolysis is functional in strains harboring p7INT-based constructs (Figure 6). The correct genome integration of the pSpy0 series and p7INT.1 was validated by nanopore sequencing for two independent clones obtained.

Next, we sought to determine the host range of the newly designed integrative plasmids. We conducted multiple sequence alignments of the *amyA*, *sagB* and *Spy_1078* integration sites found in various streptococcal serotypes. To this end, we compared the SF370 and M1T1 5448 genomic regions to genome sequences representative of M3, M4, M5, M6, M12, M18, M28, M44, M49 and M53 serotypes. We found that the *amyA* locus of SF370 is 100% conserved in M1T1 5448, 99.88% conserved in M28 GAS6180 and 99.95% conserved in the M4 GAS10750 strain (Supplementary Table 12), whereas *amyA* is not encoded in other serotypes. The *sagB* locus is well conserved between SF370, M1T1 5448 and M28 GAS6180 (100% identity) (Supplementary Table 13). While the SF370 *sagB* locus is highly conserved in M49 NZ131 (99.51% identity), M12 MGAS9429 (99.57% identity), M4 MGAS10750 (99.76%) and M44 STAB901 (99.57%), we observed less conservation (< 85% identity) in M3 MGAS315, M5 Manfredo, M6 MGAS10394, M18 MGAS8232 and M53 Alab49 (Supplementary Table 13). *Spy_1078*, similar to *amyA*, does not seem to be widely distributed among different serotypes. We found high conservation of this locus between SF370 and M1T1 5448 (99.95% identity), M44 STAB901 (99.27% identity) and M12 MGAS9429 (99.37% identity) (Supplementary Table 14). Our analysis indicates that the integrative plasmids are most likely functional in other serotypes with 100% sequence conservation, but may also be applicable in serotypes with a slightly deviating DNA sequence homology from the integration site (e.g. >99% sequence conservation).

Taken together, we present here three novel integrative plasmids that can be used in combination with the p7INT.1 plasmid to create strains harboring multiple genetic constructs of interest, e.g. for complex regulatory switches.

### 3.3 Characterization and adaptation of constitutive promoters

Constitutive promoters drive the stable expression of a protein of interest over the course of growth or under changing environmental conditions. To establish a set of promoters with different strengths, we assessed the strengths of the lactococcal P*_23_* promoter, the native P*_gyrA_*_(*Spy*)_ promoter from *S. pyogenes*, the synthetic promoter P*_xylS2_*, and the P*_veg_* promoter from *B. subtilis* by fusing each regulatory element (Figure 3B) to the luciferase reporter gene *ffluc* (*Photinus pyralis*) (13,35,56,57). The functionality of P*_23_* and P*_veg_* has previously been demonstrated in *S. pyogenes* (22,35). However, activity has only been measured for a few time points. The P*_xylS2_* promoter was originally tested in combination with a XylR repressor as an inducible system for *Streptococcus mutans* (57). We did not observe a response of this promoter to xylose (data not shown), which had already been observed in a recent study exploring the activity of the P*_xylA_* promoter in GAS (19). As the P*_xylS2_* promoter showed constant activity in the absence of xylose, we decided to test whether this promoter could serve as a heterologous constitutive promoter for *S. pyogenes*.

**Figure 3.**
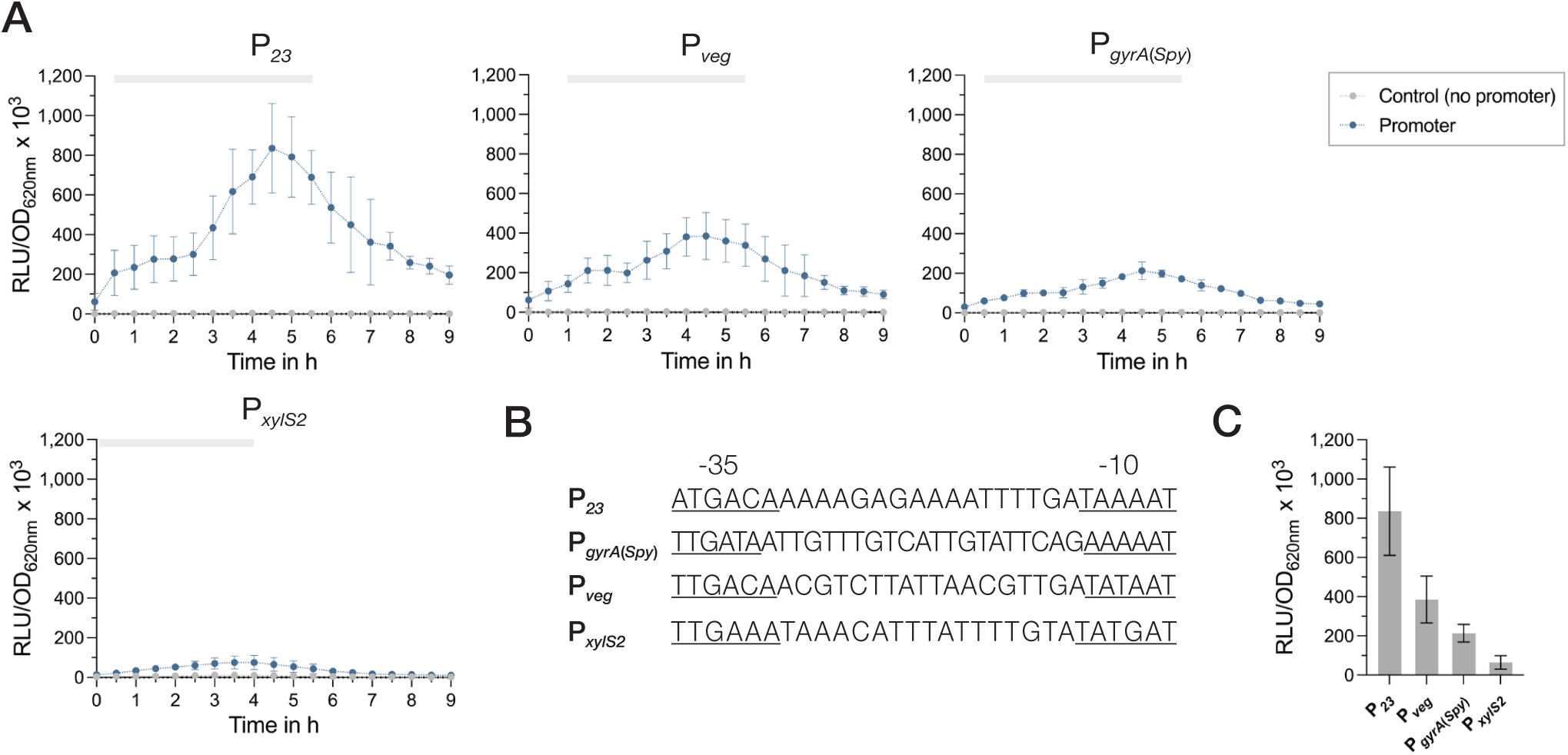
Activities of constitutive promoters for use in *S. pyogenes*. **(A)** Promoter activities of the four tested constitutive promoters P*_23_*, P*_veg_*, P*_gyrA_* and P*_xylS2_* in *S. pyogenes* SF370 grown in THY medium are shown as relative luminescence units normalized by optical density (RLU/OD_620nm_). Promoter activities of the reporter strains (blue) and the control without a promoter (grey) were measured every 30 min over the course of growth using the luciferase assay. The grey bar at the top of each graph marks the timespan of the exponential growth phase. **(B)** DNA sequences of the constitutive promoters P*_23_*, P*_veg_*, P*_gyrA_* and P*_xylS2_* comprising the -35 region, the spacer and the -10 region. The -35 and -10 regions are underlined. **(C)** Comparison of the mean promoter activities (RLU/OD_620nm_) of P*_23_*, P*_veg_*, P*_gyrA_* and P*_xylS2_* after 4.5 hours of growth in THY. Experiments were performed in biological triplicates and each measurement in technical duplicates.

All promoters exhibited their highest activity towards the end of the exponential growth phase at around 4.5 hours of growth, followed by a slow decline in the promoter signal (Figure 3A). The comparison of the promoter activities from exponential cultures (4.5 hours of growth) indicated that P*_23_* had by far the highest activity, followed by the promoters P*_veg_* and P*_gyrA_*_(*Spy*)_ (Figure 3C). The synthetic promoter P*_xylS2_* showed the lowest activity of all promoters tested (Figure 3C). For all promoter constructs, except P*_xylS2_*, we noticed a slight effect on the growth, as the cultures grew marginally slower than the control without promoter (Supplementary Figure 5A).

Next, we investigated whether we could modulate the activity of P*_veg_* by varying the spacer sequence between the -35 and -10 region of the promoter to create a set of P*_veg_* derivatives with different levels of activity. It has been shown for *E. coli* that modifying the spacer length or changing nucleotides within the spacer sequence can reduce the promoter strength (58). In the first version of P*_veg_* (P*_veg_*_(1)_), we introduced two nucleotide changes to the 17 bp between -35 and -10 (G to A and T to G) that also affect the extended -10 box sequence (Supplementary Figure 5C). For the second derivative (P*_veg_*_(2)_), we extended the spacer length from 17 bp to 19 bp through the insertion of two adenines, while for the third derivative (P*_veg_*_(3)_), we deleted two nucleotides resulting in a spacer of 15 bp (Supplementary Figure 5C). The introduced modifications resulted in a decrease in promoter activity that was not significantly different from that of the native promoter. Changing two nucleotides (P*_veg_*_(1)_) and shortening the spacer sequence (P*_veg_*_(3)_) decreased the activity by 17.2% and 10.3%, respectively (Supplementary Figure 5B and D). The extension of the spacer sequence (P*_veg_*_(2)_) showed the strongest effect, lowering promoter activity by 19.4% (Supplementary Figure 5B and D).

### 3.4 Inducible regulatory elements with high inducibility and low background expression

Next, we aimed to assess the activity of different inducible promoters in *S. pyogenes*. Both of the previously used inducible elements, P*_tet_* and P*_nisA_*, have demonstrated sub-optimal performance due to high background noise or low inducibility (Supplementary Figures 6 and 7) (19,20). Inspired by the work of Sorg and colleagues, who used the native trehalose- (P*_tre_*) and zinc-inducible promoters (P*_czcD_*) found in *S. pneumoniae* (3), we took advantage of the homologous systems from *S. pyogenes* SF370. Both promoters were placed upstream of the *ffluc* reporter gene in the replicative plasmid pSpy1C and their activity was analyzed.

We did not observe an effect on growth by the inducer compounds (Supplementary Figures 6 and 7). However, some of the constructs harboring the inducible promoters grew slightly slower than the controls without the promoter (Supplementary Figures 6 and 7). For P*_tre_*, we did not detect an increase in promoter activity after addition of 0.4% trehalose (Supplementary Figure 6A). Instead, the signal decreased during the exponential growth phase, potentially due to a catabolite repression mechanism in the presence of glucose. In contrast, the activity of the zinc-inducible promoter increased 12.8-fold when 80 µM ZnSO_4_ were added to the culture (Figure 4A). P*_Zn_* therefore showed a higher maximum fold change than the previously used inducible promoters P*_tet_* and P*_nisA_* (Figure 4B). We also tested other metal-inducible promoters, such as the promoter of the copper-specific exporter CopA (59,60) and the cation-diffusion facilitator (CDF) family transporter MntE, a paralog of the zinc exporter CzcD reported to transport manganese (61,62). We noticed only a modest induction of P*_copA_* after the addition of 80 µM CuSO_4_ and no induction of P*_mntE_*, when 80 µM MnSO_4_ were added (Supplementary Figure 8).

**Figure 4.**
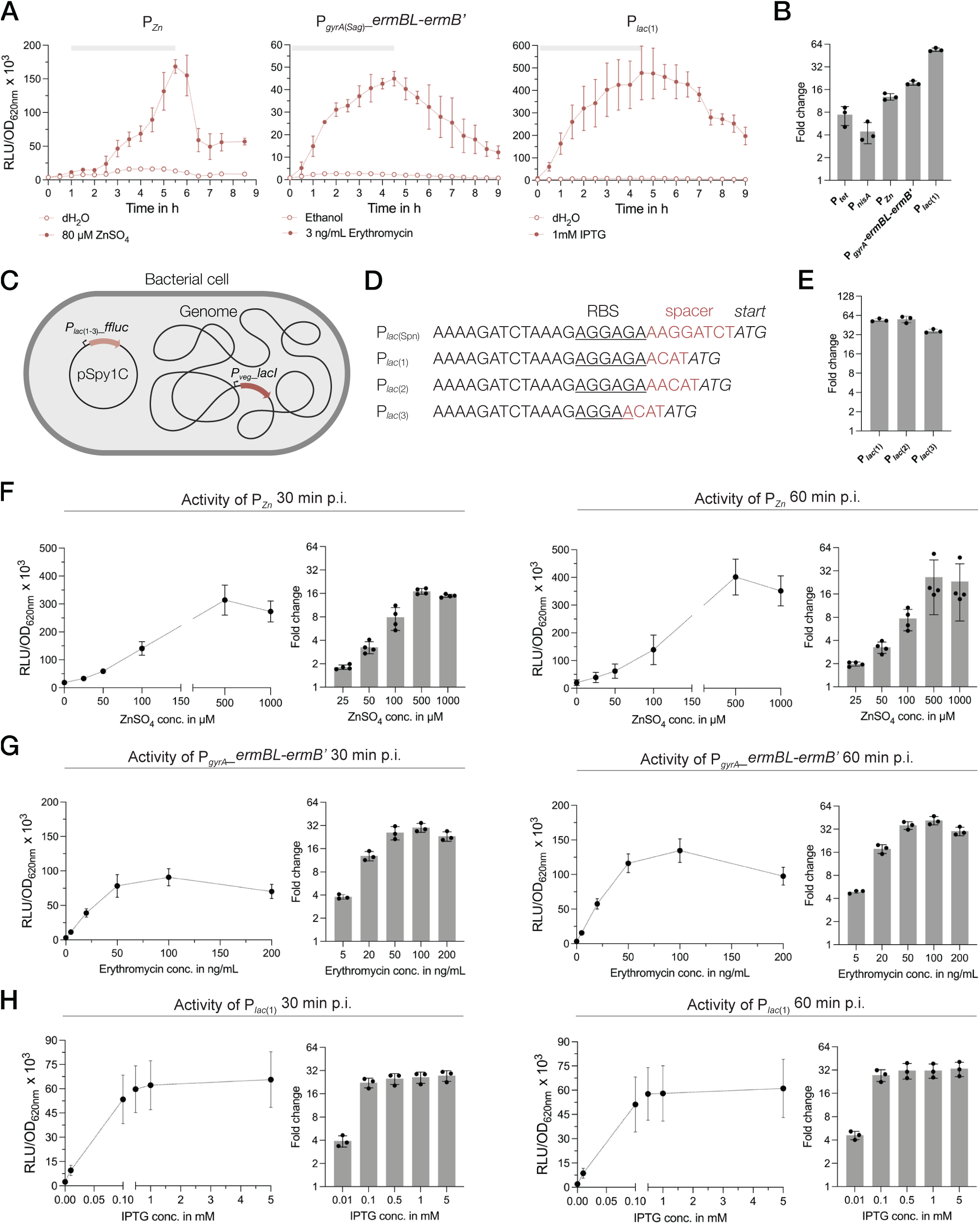
Characterization of novel inducible regulatory elements in *S. pyogenes*. **(A)** Activities of the zinc-inducible promoter, the erythromycin-inducible riboswitch and the IPTG-inducible promoter in *S. pyogenes* SF370 grown in THY medium. Promoter activities are shown in relative luminescence units normalized by optical density (RLU/OD_620nm_). Filled data points represent the signal in the presence of the respective inducer (80 µM ZnSO_4_, 3 ng/mL erythromycin and 1 mM IPTG), while empty data points show the signal in the presence of the Figure 5. The chemically defined medium RPMI4Spy enables fluorescence measurements in plate readers. **(A)** Composition of the RPMI4Spy growth medium. RPMI1640 (with glutamine and without phenol red) forms the base of this medium. **(B)** Growth of the *S. pyogenes* SF370 wildtype shown as OD_620nm_ in THY medium compared to RPMI4Spy. **(C)** Top row: Growth (OD_620nm_) of the mNeongreen reporter strain (green) and the control strain harboring mNeongreen without any promoter in THY medium (left), chemically defined medium (CDM) (center) and the RPMI4Spy medium (right). Middle row: Relative fluorescence units (RFUs) measured for medium only (black), the mNeongreen reporter strain (green) and the control strain harboring mNeongreen without any promoter (grey) in THY medium (left), CDM (center) and the RPMI4Spy medium (right). Bottom row: Relative fluorescence units calculated and normalized by optical density (RFU/OD_620nm_) measured for the mNeongreen reporter strain (green) and the control strain harboring mNeongreen without any promoter in THY medium (left), CDM (center) and the RPMI4Spy medium (right). Experiments were performed in biological triplicates and each measurement in technical duplicates.

Recently, a study reported the use of an IPTG-inducible expression system in *S. pneumoniae* (40). Based on this system, we created a reporter plasmid harboring the *lacI* repressor and the P*_lac_*_(*Spn*)_ promoter driving expression of the *ffluc* reporter (pSpy1C-PF6_*lacI*-P*_lac_*_(*Spn*)__*ffluc*). When we tested this construct, we observed low background expression and high inducibility upon induction with 1 mM IPTG (Supplementary Figure 6). We also noticed a growth delay caused by the construct, indicating that the expression of LacI from the high-copy plasmid pSpy1C had negative effects on streptococcal growth (Supplementary Figure 6). Thus, we optimized the construct and integrated the *lacI* repressor gene under control of P*_veg_* into the genome using p7INT (Figure 4C and Supplementary Table 1). The LacI-expressing strain was then transformed with three different *ffluc* reporter plasmids containing modifications downstream of the P*_lac_*_(*Spn*)_ promoter (1, 2 and 3) such as changes in the length of the spacer and RBS sequence (Figure 4D). These modifications were introduced as no viable *E. coli* clones were obtained with the original P*_lac_*_(*Spn*)_ sequence. Genomic integration of the *lacI* gene improved the growth of the strains, although not back to wildtype level (Supplementary Figure 9). The P*_lac_*_(*Spn*)_ promoter versions P*_lac_*_(1)_ and P*_lac_*_(2)_ showed very similar activity profiles over the course of growth after addition of 1 mM IPTG, with a maximum induction of 54-fold and 55.8-fold in the exponential growth phase, respectively (Figure 4A and E and Supplementary Figure 9). The P*_lac_*_(3)_ promoter showed a lower signal and reduced background noise, resulting in a 36.5-fold increase in signal during the exponential growth phase (Figure 4E and Supplementary Figure 9). When comparing the maximum induction levels reached by each inducible regulatory element, the promoters P*_lac_*_(1)_ and P*_lac_*_(2)_ displayed the highest fold change overall (Figure 4B and E).

Finally, we examined the activity of an erythromycin-sensitive riboswitch in *S. pyogenes*, which was successfully applied in a recent study (63). The P*_gyrA_*_(*Sag*)_ promoter from *Streptococcus agalactiae*, together with the erythromycin-responsive 5’UTR of the *ermB* gene, was placed upstream of the *ffluc* reporter in pSpy1C. Upon induction with 3 ng/mL erythromycin, the luminescence signal increased up to 19.5-fold during the exponential growth phase and showed low background expression in the absence of the inducer (Figure 4A). It is worth mentioning that erythromycin concentrations for induction should not exceed 3 ng/mL at low cell densities (e.g. OD_620nm_ = 0.02), whereas up to 50 ng/mL can be used without affecting growth when samples are induced during the exponential growth (e.g. OD_620nm_ = 0.25) (Figure 4A and G and Supplementary Figures 6, 7 and 10).

Next, we selected the three regulatory elements P*_lac_*_(1)_, P*_Zn_* and P*_gyrA_*_(*Sag*)_*_ermBL-ermB*’ for the analysis of their dose-response characteristics. The reporter strains were grown to the exponential growth phase and then induced with increasing concentrations of the respective inducer. We observed a dose-dependent response of P*_Zn_* for concentrations between 25 µM and 500 µM ZnSO_4_ (Figure 4F) already 30 minutes post induction. Higher concentrations, such as 1 mM ZnSO_4_, resulted in slightly inhibited growth, and thus, in a reduction in signal (Figure 4F and Supplementary Figure 10). We obtained the maximum fold induction for P*_Zn_* using 500 µM ZnSO_4_, resulting in a 17- and 26.5-fold change of the signal at 30 minutes and 60 minutes post induction, respectively (Figure 4F). Maximum activity of P*_Zn_* was reached at around 60 minutes post induction (Supplementary Figure 10).

The erythromycin-inducible riboswitch responded in a dose-dependent manner to concentrations between 5 ng/mL and 100 ng/mL of the antibiotic (Figure 4G). We observed a weak inhibitory effect on streptococcal growth for erythromycin concentrations above 100 ng/mL (Supplementary Figure 10). Consequently, the fold change in signal did not increase much further for concentrations above 50 ng/mL and even decreased slightly when 200 ng/mL erythromycin were supplemented (Figure 4G). As mentioned above, we thus recommend a maximum inducer concentration of 50 ng/mL for cultures in exponential growth phase, resulting in a 25.8- and 35.9-fold change in signal at 30 minutes and 60 minutes post induction, respectively (Figure 4G).

The P*_lac_*_(1)_ promoter showed the fastest response, with maximum induction reached already 30 minutes post addition of IPTG, irrespective of the inducer concentration (Supplementary Figure 10). While adding 0.01 mM IPTG triggered only a 3.9-fold increase in signal at 30 min post induction, the fold-change increased and varied only slightly when concentrations between 0.1 mM and 5 mM IPTG were applied (22.3 to 27.5-fold) (Figure 4H). These values increased only marginally after 60 minutes with 27.5 to 33.3-fold for concentrations ranging from 0.1 mM to 5 mM, respectively (Figure 4H). As expected, the addition of IPTG, even at high concentrations, had no inhibitory effect on growth (Supplementary Figure 10).

In summary, we found that P*_lac_*_(1)_, P*_Zn_* and P*_gyrA_*_(*Sag*)_*_ermBL-ermB*’ enable inducible gene expression in *S. pyogenes,* showing high inducibility, dose-dependency and low-to-no background expression.

### 3.5 Optimizing the application of fluorescence reporters in *S. pyogenes*

Early studies on the use of GFP in *S. pyogenes* suggested that growth conditions, such as low oxygen and acidification of the growth medium, contradict the function of FPs (24). Only a few studies subsequently demonstrated the use of FPs, such as mCherry or GFP, to label GAS to monitor the progression of infection or determine protein localization (7,23,25). However, the performance of newer generations of fluorescent reporters with improved features has yet to be evaluated.

When we initially attempted to analyze the performance of the fluorescent reporter mNeongreen in the plate reader, we were unable to detect a signal in THY medium. Indeed, it has been described previously for Group B Streptococci (GBS) that fluorescence measurements are difficult due to the strong autofluorescence of the growth medium (64). Thus, to enable continuous fluorescence measurements in the plate reader, we developed a growth medium with reduced autofluorescence.

Earlier, Schulz and colleagues adapted the cell culture medium RPMI1640 to facilitate growth of *S. pneumoniae* (65,66). RPMI1640 is available without phenol red and its colorless appearance presented a good starting point for the adaptation of a novel chemically defined medium for *S. pyogenes*. We established several formulations until *S. pyogenes* growth in this medium was comparable to that in other commonly used chemically defined medium (CDM) (Supplementary Figure 11 and Supplementary Table 11). The final recipe, adapted from a formulation published while we worked on the composition (67), contains a Nucleobase mix, the commercially available BME vitamin and RPMI amino acid solutions, glucose and niacinamide, and is further buffered with HEPES buffer (Figure 5A). When we compared the growth of *S. pyogenes* SF370 in THY and RPMI4Spy, bacteria grew to a higher final optical density in the nutrient-rich THY medium, reaching a maximum OD_620nm_ of ∼0.5 in THY and ∼0.35 in RPMI4Spy (Figure 5B). However, the growth curve profiles appeared similar, showing neither a prolonged lag phase nor an early onset of cell death in the stationary growth phase (Figure 5B).

**Figure 5.**
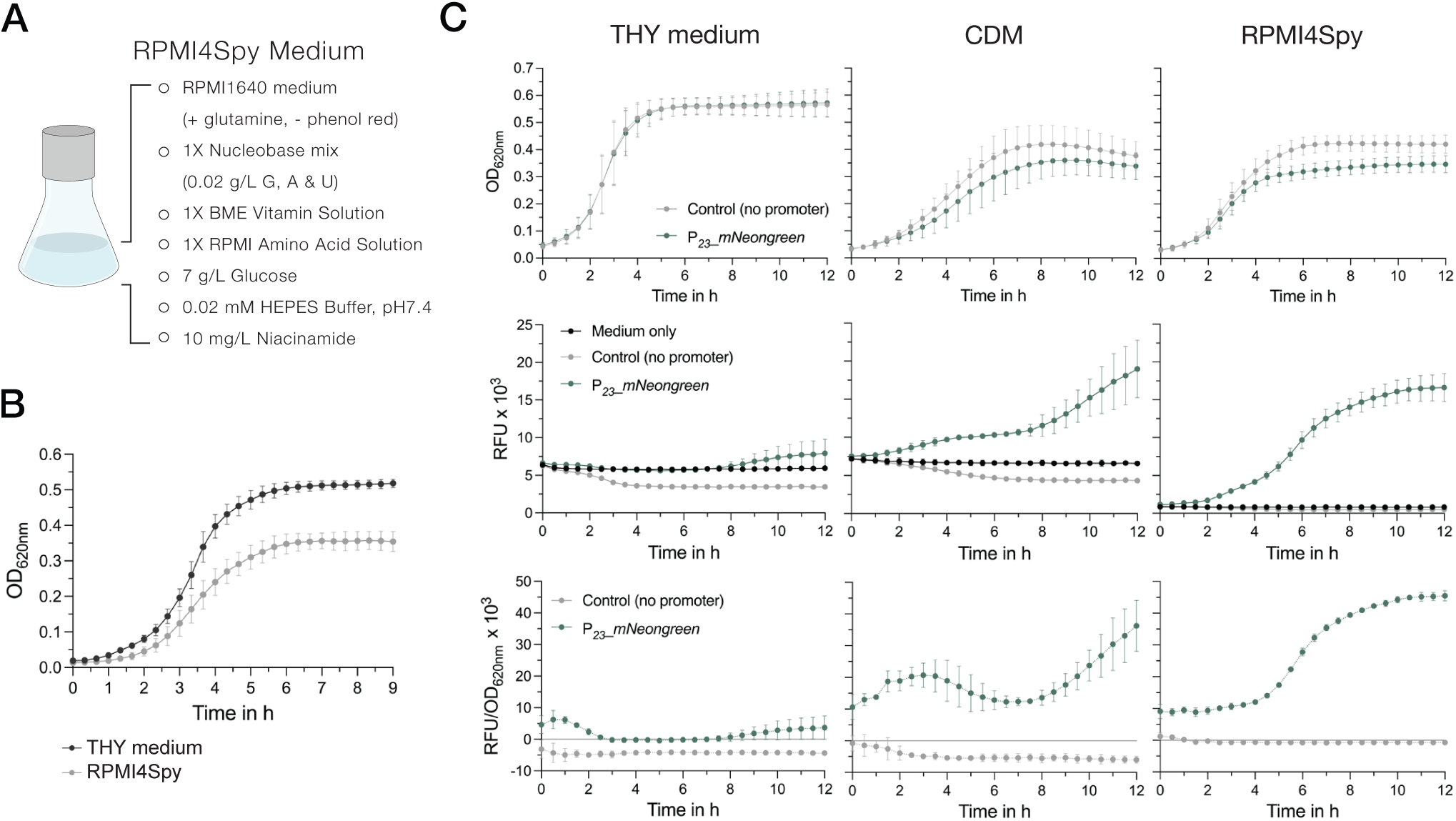
The chemically defined medium RPMI4Spy enables fluorescence measurements in plate readers. **(A)** Composition of the RPMI4Spy growth medium. RPMI1640 (with glutamine and without phenol red) forms the base of this medium. **(B)** Growth of the *S. pyogenes* SF370 wildtype shown as OD_620nm_ in THY medium compared to RPMI4Spy. **(C)** Top row: Growth (OD_620nm_) of the mNeongreen reporter strain (green) and the control strain harboring mNeongreen without any promoter in THY medium (left), chemically defined medium (CDM) (center) and the RPMI4Spy medium (right). Middle row: Relative fluorescence units (RFUs) measured for medium only (black), the mNeongreen reporter strain (green) and the control strain harboring mNeongreen without any promoter (grey) in THY medium (left), CDM (center) and the RPMI4Spy medium (right). Bottom row: Relative fluorescence units calculated and normalized by optical density (RFU/OD_620nm_) measured for the mNeongreen reporter strain (green) and the control strain harboring mNeongreen without any promoter in THY medium (left), CDM (center) and the RPMI4Spy medium (right). Experiments were performed in biological triplicates and each measurement in technical duplicates.

Next, we assessed the differences in fluorescence signals of the mNeongreen reporter strain grown in THY, CDM or RPMI4Spy. In both chemically defined media, CDM and RPMI4Spy, the mNeongreen reporter strain showed reduced growth, indicating a potential fitness burden in the defined media due to the high expression of mNeongreen compared to the control (Figure 5C). This was not observed in nutrient-rich THY medium (Figure 5C). When comparing the relative fluorescence units (RFU) of the growth medium itself with the control and reporter strain, we found that the signal of medium and reporter in THY medium overlap, creating the impression that the reporter strain is not producing mNeongreen (Figure 5C). Despite the autofluorescence of the CDM, we detected an increase in mNeongreen signal after several hours of growth (Figure 5C). We hypothesize that, in comparison to THY, CDM contains lower concentrations of autofluorescence-causing vitamins, such as riboflavin (68). Once riboflavin is taken up and used by the bacteria, the autofluorescence decreases, which is not observed in the medium control without bacteria (Figure 5C). In line with this, the signal of the control strain was lower than the autofluorescence of the medium, resulting in negative values when subtracting the blank and interfering with the subsequent calculation of a normalized fluorescence (Figure 5C). However, the RFU values of control strain and medium for RPMI4Spy were highly similar, and mNeongreen fluorescence was well distinguishable during growth (Figure 5C). Taken together, our results demonstrate that RPMI4Spy enables *S. pyogenes* to grow at optical densities similar to those of the chemically defined medium (CDM) previously used, and exhibits reduced autofluorescence that allows continuous fluorescence measurements in a plate reader.

Next, we explored the use of the two codon-optimized FPs mKate2 and mNeongreen, which are characterized by enhanced stability at lower pH and increased brightness. To test their functionality in *S. pyogenes*, both reporter genes under the control of P*_23_* and the respective control harboring each reporter gene without the promoter were integrated using p7INT (Supplementary Table 1). We then grew the strains in RPMI4Spy medium and took samples at the exponential growth phase for fluorescence microscopy. Cells expressing the mNeongreen or mKate2 reporter displayed a detectable green and red fluorescence, respectively, while the control strains without the promoter exhibited no fluorescence (Figure 6A and B). The fluorescence intensity varied slightly between individual cells (Figure 6A and B).

**Figure 6.**
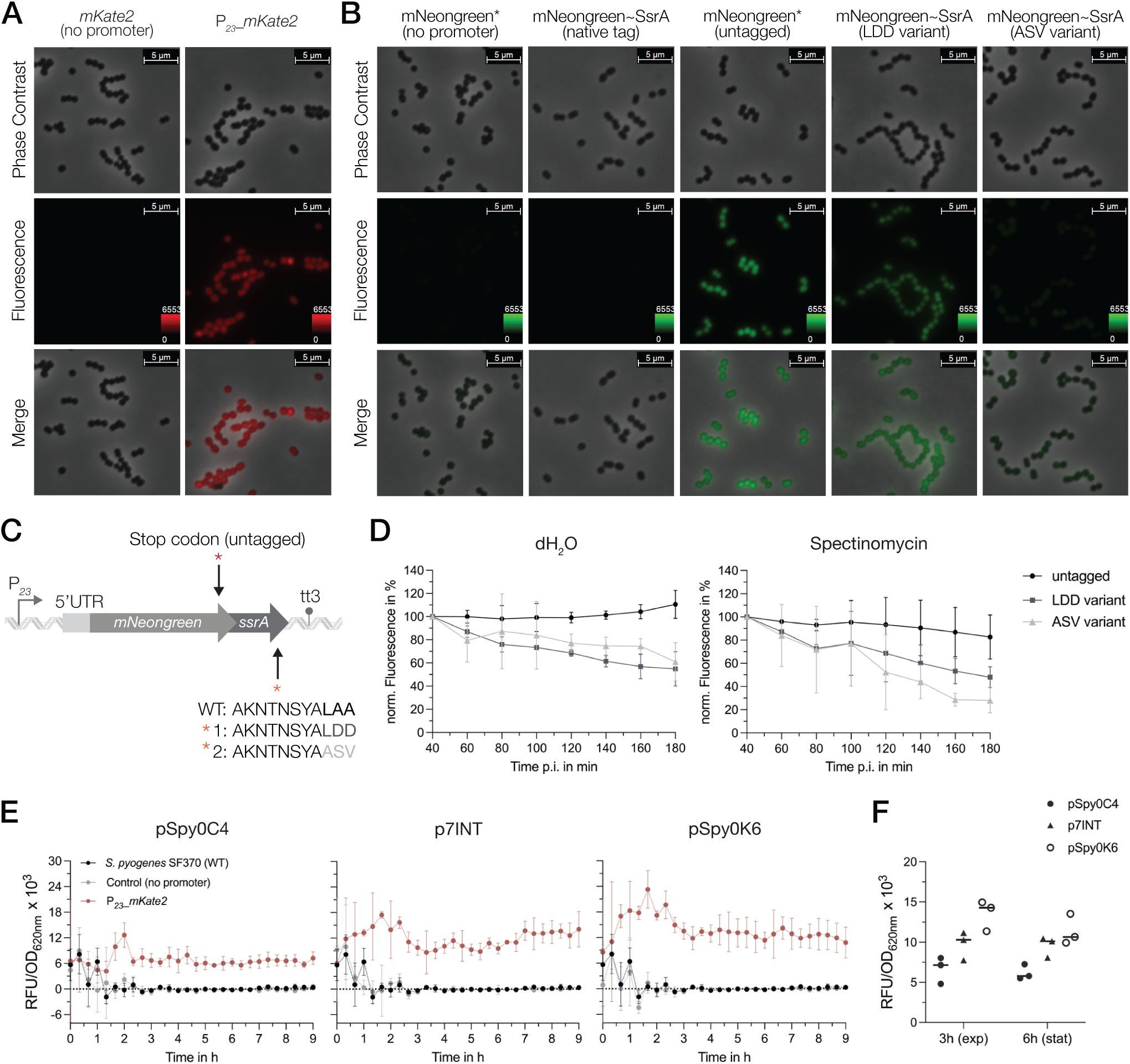
Characterization of the mNeongreen and mKate2 fluorescence reporters in *S. pyogenes*. **(A)** Fluorescence microscopy with the mKate2 reporter strain (right panel) and the respective control harboring the *mKate2* gene without the promoter (left panel). The top and middle rows show images from phase contrast and fluorescence channel, respectively. The bottom row displays a merged image of both channels. Representative images of three replicates are shown. **(B)** Fluorescence microscopy with the mNeongreen reporter strain (untagged reporter, third panel), mNeongreen fused to different SsrA degradation tag variants (native tag in the second, LDD variant tag in the fourth and ASV variant tag in the fifth panel) and the respective control strain harboring the *mNeongreen* gene without the promoter and without the SsrA tag (first panel). Top and middle rows show images from phase contrast and fluorescence channel, respectively. The bottom row displays a merged image of both channels. Representative images of three replicates are shown. **(C)** Schematic of the genetic construct design of the SsrA tag fusions to mNeongreen (native tag ending in LAA, variant 1 with an LAA to LDD mutation and variant 2 with LAA to ASV mutation) and the control harboring a stop codon (TAA) preceding the coding sequence of the *ssrA* tag. **(D)** Translation inhibition assay using spectinomycin as inhibitor to monitor the degradation of mNeongreen fused to the different SsrA tag variants over time. Left: Development of the mNeongreen signal after addition of distilled water (dH_2_O) as control. Right: Changes in mNeongreen signal after translation inhibition using 100 µg/mL Spectinomycin. **(E)** The mKate2 reporter signal (RFU/OD_620nm_) of *S. pyogenes* strains expressing the reporter from different genomic sites: pSpy0C4 (*sagB*), p7INT (*attB*(T12)) and pSpy0K6 (*SPy_1078*). The fluorescence signals are depicted as RFU/OD_620nm_ for the strain expressing mKate2 (red), the control harboring mKate2 without a promoter (grey) and the SF70 wildtype (black) for each individual integration site. **(F)** Comparison of the fluorescence signal (RFU/OD_620nm_) produced by the three different genomic integration sites in the exponential growth phase (3h) and early stationary growth phase (6h). Experiments were performed in biological triplicates and each measurement in technical duplicates.

When using FPs as reporters for gene expression, their high stability can be disadvantageous for the detection of transient changes in transcriptional activity. To overcome this problem, unstable variants have been developed in which the FP is fused to a degradation tag, thereby taking advantage of SsrA tag-mediated proteolysis (69,70). It has been shown that the last three amino acids of the SsrA tag influence the degradation rate of a protein (69,71). Thus, based on the native SsrA tag sequence from *S. pyogenes* (AKNTNSYALAA), we designed different SsrA tag variants that were fused to mNeongreen including (1) an untagged control with a stop codon introduced upstream of the native tag (mNeongreen*), (2) a mutant in which the last three amino acids were changed from LAA to LDD (LDD variant) and (3) a mutant in which the last three amino acids were modified to ASV (ASV variant) to modulate stability of the reporter (Figure 6C and Supplementary Table 1). We then compared the effects of the SsrA tag variants on the fluorescence signal using microscopy. No fluorescence was observed when mNeongreen was fused to the native SsrA tag, indicating a high degradation rate of the reporter (Figure 6B). In comparison to the untagged mNeongreen (mNeongreen*), fluorescence of the FP tagged with the LDD variant appeared less bright, indicating a slight destabilization (Figure 6B). Interestingly, tagging mNeongreen with the ASV variant led to even higher destabilization, with cells emitting almost no fluorescence under the microscope (Figure 6B). To investigate this further, we performed a plate reader assay to determine the decrease in signal upon global inhibition of translation using spectinomycin (Figure 6D). Both, the LDD variant and the ASV variant tag resulted in a decline in fluorescence of about 52% and 72%, respectively, 180 minutes after the treatment (Figure 6D). For the untagged control, the signal decreased only marginally upon spectinomycin treatment (∼17% at 180 min pi.), while the fluorescence continued to increase when distilled water was added, indicating high stability of the untagged mNeongreen protein under normal growth conditions (Figure 6D).

Finally, we thought to assess the effect of the genomic context on the expression of the reporters. Therefore, we compared the fluorescence signals of the mKate2 reporter from three different integration sites: *sagB*, *attB*(T12) and *SPy_1078* (Figure 2B). To do so, *mKate2* was cloned into pSpy0C4, p7INT and pSpy0K6 with or without the P*_23_* promoter, respectively (Supplementary Table 1). We then measured the fluorescence signal over the course of growth in a plate reader. Our data indicate a weak but steady expression of mKate2 from the *sagB* locus (Figure 6E). When integrated into the *attB* site of the T12 phage, the fluorescence signal appeared slightly stronger and increased gradually over time (Figure 6E). Interestingly, integration of the reporter into *Spy_1078* resulted in the highest fluorescence signal (Figure 6E). However, this signal began to decrease towards stationary growth phase (Figure 6E). The differences in signal between the integration sites became more apparent when we directly compared the mKate2 signals from the exponential (3h) and the early stationary growth phase (6h). Indeed, integration using pSpy0K6 resulted in the highest fluorescence signal (Figure 6F). However, this signal decreased at 6 hours, whereas it remained stable when pSpy0C4 or p7INT were used for integration (Figure 6F). The growth of *S. pyogenes* did not appear to be affected by the different integrated constructs (Supplementary Figure 12). We conclude that the integration site, and thus, the surrounding genomic context may influence the heterologous expression of a gene of interest. Depending on the aim of an experiment, the genomic location of the construct must be chosen accordingly.

### 3.6 pERASE for scarless gene deletions in *S. pyogenes*

For the implementation of gene deletions in *S. pyogenes*, researchers have recently mainly relied on allelic replacement of the gene with an antibiotic resistance cassette, plasmids harboring thermosensitive replicons or the Cre-*lox* system. Particularly the last two strategies involve a time-consuming and laborious process for plasmid curing or Cre-*lox* cassette removal. Additionally, the latter procedure leaves a *lox* scar site in the genome, which in some scenarios might disrupt important regulatory sequences.

To accelerate this process and enable straightforward selection of putative scarless deletion mutants, we designed the gene editing plasmid pERASE. This plasmid features the same MCS as the other plasmids in this collection, an erythromycin resistance cassette and the high-copy origin pUC replicating in Gram-negative bacteria (Figure 7A and Supplementary Figure 4C). Moreover, the plasmid harbors the *pheS\** counterselection marker encoding a modified phenylalanyl-tRNA-synthetase (*α*-subunit) of which expression is controlled by P*_veg_* (Figure 7A and Supplementary Figure 4C). The *pheS* gene sequence was obtained from the *S. pyogenes* genome and adapted by introducing silent mutations to the *pheS* coding sequence to prevent recombination events at the native streptococcal *pheS* locus. Moreover, two mutations were inserted to create a *pheS*\* variant which, when expressed, incorporates the toxic halogenated analogue of phenylalanine (4-chloro-phenylalanine, short PCPA or 4CP) into proteins during translation, leading to cell death. Selection on agar plates containing PCPA therefore only allows the growth of mutants that have lost the plasmid (Figure 7A). As PCPA is itself toxic at a certain concentration, we performed an initial test experiment using TSA agar plates containing various concentrations of PCPA and found that 0.6% PCPA (30 mM) resulted in growth inhibition of the *S. pyogenes* SF370 wildtype (data not shown). Thus, a lower concentration (5 mM) was selected to ensure that only bacteria still harboring the plasmid in the genome had their growth suppressed, while deletion mutants and wildtype revertants could grow.

**Figure 7.**
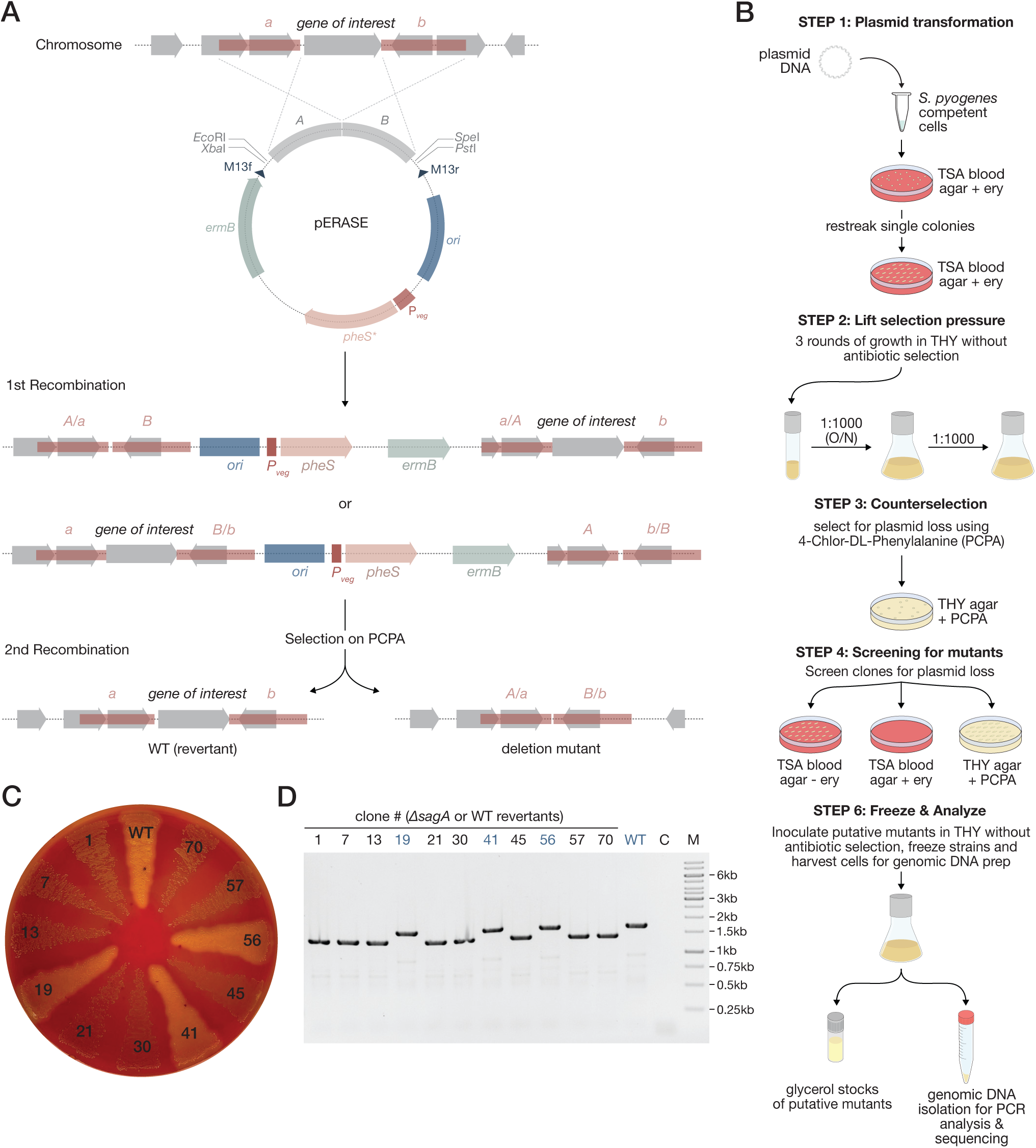
Creating scarless gene deletions using pERASE. **(A)** Schematic representation of the pERASE plasmid, subsequent recombination events after transformation into *S. pyogenes* and plasmid curing using selection on 4-chloro-phenylalanine (PCPA) agar plates. **(B)** Workflow showing the individual steps for mutant generation with pERASE in *S. pyogenes*. After transformation into *S. pyogenes*, clones with the integrated plasmid are selected on TSA blood agar with erythromycin. The plasmid is cured from the genome using three passages in liquid THY medium without antibiotic selection. Clones that have lost the plasmid are selected on THY agar plates containing 5 mM PCPA. Putative mutants growing on PCPA plates are then streaked onto TSA agar plates with and without erythromycin, as well as on THY agar plates with PCPA. For mutants with no visible phenotype, such as the loss of hemolysis, a colony PCR step can be introduced here. Several putative mutants are inoculated into THY medium to prepare glycerol stocks and isolate genomic DNA for PCR analysis. **(C)** TSA blood agar plate showing the loss of hemolysis of the putative *sagA* mutants compared to the *S. pyogenes* wildtype (WT) and wildtype revertants such as clones 19, 41 and 56. **(D)** Agarose gel showing PCR amplicons of the *sagA* region. A DNA band of ∼1.4 kb indicates the loss of *sagA*, while a band of 1.7 kb in size indicates that *sagA* is still present in the genome. WT = wildtype; C = Control without template; M = Marker (1kb GeneRuler^TM^ from Thermo Scientific^TM^).

To test the applicability of pERASE for generating scarless gene deletions in *S. pyogenes*, we aimed to delete the *sagA* gene encoding the toxin streptolysin S. Deletion of this gene results in a non-hemolytic phenotype on TSA blood plates compared to the hemolytic wildtype strain. To this end, we cloned pERASE-*ΔsagA*, containing homologous fragments upstream and downstream of *sagA*. After transformation of the plasmid into *S. pyogenes* SF370, we selected clones that had undergone homologous recombination and integrated the plasmid on TSA blood plates containing erythromycin (Figure 7B). After one passage on TSA blood plates, we inoculated single clones into THY medium without antibiotics to allow for a second recombination event and to promote subsequent loss of the plasmid (Figure 7B). We performed three passages in liquid culture before selecting for plasmid loss on THY agar plates containing 5 mM PCPA (Figure 7B). Several clones (n=70) growing in the presence of PCPA were again streaked onto TSA blood plates with and without erythromycin and onto THY agar plates with 5 mM PCPA (Figure 7B). Putative deletion mutants and wildtype revertants were expected to grow on TSA blood plates without antibiotics and on the PCPA plates, but not on TSA blood plates with erythromycin. We identified 17 out of the 70 clones (∼24%) that exhibited a loss of hemolysis on TSA blood plates. From these 17 clones, we selected eight clones showing no hemolysis, while from the remaining 53 clones displaying a hemolytic phenotype, we selected three clones as controls (representing potential wildtype revertants) for PCR analysis (Figure 7C and D). The absence of *sagA* in these eight clones was proven by PCR and Sanger sequencing of the three final clones and the results were consistent with the observed non-hemolytic phenotype (Figure 7C and D). These results prove that pERASE can indeed be applied to generate clean deletion mutants in just eight days.

## 4 Discussion

### 4.1 *E. coli*-*S. pyogenes* shuttle plasmids for rapid cloning and heterologous gene expression

In this study, we generated a novel set of low- and high-copy replicative plasmids and demonstrated the applicability of pSpy1C for the creation of reporter plasmids and the subsequent assessment of promoter activities in GAS. The modular structure of the plasmids and uniform design of the MCS enable straightforward cloning procedures in *E. coli*. It is noteworthy that RCR derivatives, such as pSH71 and pWV01* are considered less stable when harboring inserts larger than 8 kb, whereas theta replicons like pAMβ1 show higher stability (72). We therefore recommend using plasmids of the low-copy pSpy2 series when working with very large inserts, while high-copy plasmids of the pSpy1 and pSpy3 series can be applied for smaller inserts and high expression levels.

When *S. pyogenes* was transformed with the empty plasmids, we found that the cultures displayed varying growth profiles. One reason for a plasmid-caused growth delay could be a divergent directionality of genes encoded on the plasmid, leading to interference of replication and transcription processes (73,74). However, this did not seem to be the case as all open reading frames encoded on e.g. pSpy2C show the same directionality. We suspect this observation to be an effect of the plasmid copy number (PCN) and, consequently, a gene dosage effect of the antibiotic resistance-conferring gene encoded on each plasmid, given that growth profiles varied between the antibiotic markers. It has been shown that higher PCNs are associated with increased resistance levels against antibiotics (75). This is consistent with the results from the adaptation of the antibiotic concentration, where a decreased chloramphenicol concentration resulted in accelerated growth of *S. pyogenes* harboring the low-copy plasmid pSpy2C, while still ensuring plasmid maintenance. We propose that further adaptation of the selective concentrations of kanamycin and erythromycin will result in a higher overlap of the growth curves for each plasmid series.

### 4.2 Integrative plasmids allow for the stable and site-specific integration of genes of interest into the streptococcal genome

To enable the stable integration of genetic elements into the genome and thereby avoid PCN effects on the expression of a gene of interest, integrative plasmids have been widely applied in various bacterial model organisms (13,14). To our knowledge, only two site-specific integrative plasmids for use in GAS have been described: pFW11e integrating into a gene coding for a sugar phosphate isomerase (*SPy_0535*) and p7INT inserting at the 3’end of the tmRNA locus (16,22,76). Compared to integrative plasmids previously used, pSpy0C2 and pSpy0C4 allow direct selection of integration events on starch and blood plates, respectively. It is worth noting that both genes, *amyA* and *sagB,* are involved in streptococcal virulence, which needs to be taken into account when performing infection experiments (77,78). As an alternative, we designed pSpy0K6 that integrates into a transcriptionally silent locus, offering two main advantages. First, the integrated DNA fragments are not affected by any read-through from strongly transcribed surrounding genomic regions (79), and secondly, the integration does not disrupt a functional gene required for growth or virulence.

The integration sites for pSpy0C2, pSpy0C4 and pSpy0K6 are highly conserved between SF370 and the more clinically relevant M1T1 5448 strain. We also found GAS strains of other serotypes, such as M28 MGAS6180 or M12 MGAS9429, to have high DNA sequence identity with the integration sites, potentially expanding the host range of these plasmids to other streptococcal strains. Shortening the plasmid-encoded homologous regions required for recombination could result in higher sequence homology with other serotypes, and thus, a broader host range of the plasmids. Previous research suggests the use of homologous regions between 250 and 500 bp in length, although longer sequences typically achieve higher recombination efficiencies (10). Because of the different resistance cassettes, these plasmids can also be used in combination, enabling more advanced genetic modifications. Finally, the provided list of transcriptionally silent sites found in SF370 and M1T1 5448 will enable other research groups focusing on GAS to design further site-specific integrative plasmids.

### 4.3 Novel promoters for controlled gene expression in *S. pyogenes*

In this study, we assessed the activity of different constitutive promoters and tested potential novel inducible promoters for tunable gene expression in GAS. Our side-by-side comparison of the different constitutive promoters highlighted P*_23_* and P*_veg_* as the strongest promoters available, while P*_gyrA_* and P*_xylS2_* showed weaker activities. These promoters cover a wide range of expression levels that researchers can select from depending on the experimental setup. Strong promoters are advantageous when studying the effect of gene overexpression on the cell or to obtain higher yields in protein purification procedures, while low expression may be preferred when studying toxic proteins or poorly soluble proteins (80). To obtain an increased variety in promoter strengths to select from, we modified the P*_veg_* promoter by mutating the spacer sequence between the -35 and -10 region of the promoter. Although we observed a decrease in promoter activity, these modifications did not result in a significant reduction of the promoter strength. Consistent with these results, a recent study demonstrated for a set of *E. coli σ*^70^-promoters that changes in spacer length from 15 bp to 19 bp have no significant impact on promoter strength, whereas longer spacers (>19 bp) result in significantly lower expression levels (58).

As the number of inducible regulatory elements for *S. pyogenes* is very limited, we sought to evaluate different inducible promoters for use in GAS. The two commonly used inducible promoters, P*_tet_* and P*_nisA_*, showed high leakage and low inducibility in our experiments, respectively. In contrast, the three novel inducible regulatory systems described here outperformed the previous systems, demonstrating high inducibility, no-to-low background expression and a dose-dependent response to different inducer concentrations. The P*_lac_* promoter performed best when the LacI repressor was constitutively expressed from the streptococcal genome, consistent with previous observations made in *S. pneumoniae* (14). We hypothesize that high LacI expression levels from replicative plasmids could result in unspecific binding of the repressor to DNA sequences with high similarity to LacI operator sites, potentially affecting the expression of genes involved in the streptococcal metabolism, and thus, growth. Nevertheless, we observed a slight influence on streptococcal growth for the strain expressing LacI from the bacterial chromosome, independently of the presence of the reporter plasmid. Potentially, replacing P*_veg_* driving *lacI* transcription with a weaker promoter, e.g. P*_xylS2_*, could resolve this minor growth defect. However, the reduced expression of the LacI repressor may result in a higher background noise of the promoter when used in combination with a high-copy reporter plasmid.

Compared to e.g. P*_lac_*_(1)_, our data demonstrated a high tunability of P*_Zn_* induction levels, making this promoter interesting for the study of dose-dependent effects. It is noteworthy that a tight control of metal ion homeostasis is critical for growth and survival of GAS, and thus, inducer concentrations need to be selected appropriately. During infection, *S. pyogenes* not only encounters host-mediated zinc starvation through redistribution or sequestration of zinc at the site of infection, but also zinc toxicity after phagocytosis by macrophages and neutrophils to limit bacterial growth (81).

RNA-based regulatory systems, such as riboswitches, are usually highly conserved between bacteria in comparison to protein-based inducible expression systems and, consequently, are better transferrable (82). Here, we have successfully demonstrated the applicability of an erythromycin-inducible riboswitch in *S. pyogenes*. We like to highlight that this system is not the only RNA-based regulatory system applicable to GAS, as another study demonstrated the use of a theophylline-sensitive synthetic riboswitch in *S. pyogenes,* showing high inducibility and very low basal expression in the absence of the inducer (83).

Attempts to establish the trehalose-inducible promoter (P*_tre_*) did not prove successful, probably due to carbon catabolite repression. We hypothesize that glucose utilization by GAS led to downregulation of this promoter during exponential growth. Previous research suggests that the trehalose operon is repressed by the catabolite control protein A (CcpA), despite a direct binding by CcpA could not be proven by ChIP-seq analysis (84). Potentially, induction could be observed in a growth medium devoid of glucose or fructose, which were shown to be the preferred carbon source for GAS (85).

For all luciferase reporter strains we observed a decrease of the luminescence signal towards stationary growth phase. It was proposed that the decline in luminescence observed in the transition and stationary growth phase could result from the associated decrease in intracellular ATP levels, subsequently leading to less availability of ATP for the conversion of the substrate D-luciferin to oxyluciferin and light (Figure 3A) (86,87). When stationary phase cultures are being studied, ATP-independent reporters, such as the luciferases from *Renilla reniformis* or *Gaussia princeps* could be applied (88).

### 4.4 mNeongreen and mKate2 as reporters for gene expression in GAS

Although a wide range of luciferases exists, these reporters are limited in their application as they, e.g., do not support studying protein localization. To provide alternative reporters for *S. pyogenes*, we aimed to assess the functionality of fluorescent reporters of newer generations with improved properties, such as increased brightness and higher stability at acidic pH.

Interestingly, although GFP and its derivatives have been widely used in many other Gram-negative and Gram-positive bacteria, the number of studies using fluorescence reporters in GAS is very limited (23–25). Here, we demonstrated that new generations of fluorescence proteins, such as mNeongreen and mKate2, can indeed be applied in GAS and, in combination with the new genetic toolset, could serve to understand protein localization, gene expression or protein-protein interactions.

Previous research already highlighted difficulties in measuring fluorescence signals in Group B Streptococci (64). Responsible for this autofluorescence are vitamin components in the medium, such as riboflavin, which is mainly responsible for green autofluorescence (68). One possibility to circumvent this problem is to wash and resuspend the cells in 1x PBS buffer to measure fluorescence. However, the required washing step is time-consuming and results in deviations from the original optical density of the culture. The RPMI4Spy medium was adapted from a previously published recipe in which growth of the GAS clinical isolate HKU16 was demonstrated (67). The medium showed a greatly decreased autofluorescence that enabled the monitoring of fluorescence signals from bacteria expressing mNeongreen or mKate2 over the course of growth, indicating no decrease in fluorescence signal when cells transitioned to the stationary phase. Not surprisingly, expressing mNeongreen at high levels seems to represent a fitness burden for *S. pyogenes* when cultured in chemically defined media, whereas no effect on growth was observed in nutrient-rich THY medium. Finally, RPMI4Spy opens up the use of methodologies that have previously been only rarely applied on GAS, such as time-lapse fluorescence microscopy (89).

When using fluorescent proteins as a readout for gene expression, their high stability can constitute a disadvantage as transient changes in gene expression, particularly downregulation, may not be accurately represented. Therefore, we tested the modulation of mNeongreen protein stability by adding different SsrA degradation tag variants. We found that mutating the last three amino acids of the native SsrA tag sequence altered mNeongreen stability to varying degrees. Although we barely detected a fluorescence signal for the ASV variant, the signal of the LDD variant differed only slightly from the untagged reporter. Although our data on the ASV variant mainly reflect results from previous studies in *Mycobacterium* and *E. coli* showing a similar degree of destabilization, the overall fluorescence intensity of this variant is strongly decreased in *S. pyogenes* (69,70,90,91). This indicates that many factors play a role in the design of such degradation tags, including the complete SsrA tag sequence, the microorganism used as chassis, and thus, the present degradation machinery (71). No signal was detected from mNeongreen fused to the native tag, which is also in agreement with previous research (70).

When we assessed the signal of the mKate2 reporter from different genomic locations, we noticed that integrations into *SPy_1078* using pSpy0K6 resulted in the highest signals. It has been shown that the proximity of a heterologous gene to the origin of replication influences its expression levels due to gene dosage effects during replication (79,92). In this case, all integration sites are located quite far from the origin of replication, and thus, expression is expected to be lower. The replication terminus of *S. pyogenes* SF370 is not well characterized, but a *dif*-like termination sequence identical to that in *E. coli* was identified, starting at position 929,320 (8). Interestingly, of the three integration sites evaluated, *SPy_1078* is the closest to the terminus and should, according to previous studies, demonstrate the lowest expression. As this was not the case, gene dosage cannot be the explanation here. A recent study was similarly unable to correlate gene dosage effects with expression levels of a fluorescent reporter expressed from different genomic regions in *E. coli* (93). The authors found that, although the same promoter was placed in front of the reporter, the RNA polymerase occupancy varied between the different genomic sites resulting in different fluorescence signal intensities (93). In line with this, recent findings showed that transcriptional regulation in bacteria is also mediated by chromosome re-modelling via nucleoid associated proteins (NAPs), making genes accessible or inaccessible to transcription depending on different growth conditions, such as stress (94). The changes in fluorescence signal observed for mKate2 between the exponential and early stationary phase could result from such chromatin re-modelling. Consequently, a variety of parameters can influence the expression of a heterologous gene from a specific genomic locus.

Finally, it remains to be investigated whether alternative reporters, e.g. fluorescent RNA aptamers such as Spinach or Mango, may also be applied in GAS (95–97). Such systems could be advantageous for the study of bacterial transcription, as the signal is directly proportional to RNA levels.

### 4.5 pERASE facilitates gene editing in GAS

Several studies showed the applicability of the *pheS\** counterselection marker for genome editing in various bacterial species (98–101). According to a recent review, the *pheS\** counterselection marker has already been successfully applied in the genetic manipulation of *S. pyogenes* by Caparon and Port (unpublished) (10). However, genetic engineering in GAS currently still relies on plasmids harboring thermosensitive origins, transposon-based systems or the Cre-*lox* strategy (10,32,102).

Unlike thermosensitive shuttle vectors, such as the recently published pBFK, pERASE offers several advantages, such as its minimal size, no need for temperature shifts, a counterselection marker to screen for plasmid loss and the possibility of red-white screening during cloning in *E. coli* (103). Particularly the use of a counterselection marker reduces the workflow for mutant generation to eight days as demonstrated for the clean deletion of *sagA*. We believe that this time period can be further shortened by reducing the number of rounds for plasmid curing in liquid THY or by adding PCPA already at this stage. The latter approach was previously shown for counterselection after transformation into *S. pneumoniae* and the protocol could therefore be further optimized (104). Moreover, we believe that pERASE can be applied not only for gene deletions but also for the introduction of point mutations, insertion of genes or small tags.

## 5 Author contributions

NL: Conceptualization, Methodology, Investigation, Validation, Formal analysis, Visualization, Writing – original draft. KS: Investigation, Validation, Writing – review & editing. CS: Conceptualization, Methodology, Investigation, Writing – review & editing. TFW: Conceptualization, Methodology, Formal analysis, Software, Writing – review & editing. KH: Investigation, Validation, Writing – review & editing. KF: Conceptualization, Formal analysis, Software, Visualization, Writing – review & editing. MM: Supervision, Writing – review & editing. AKWE: Conceptualization, Supervision. EC: Supervision, Funding acquisition, Writing – review & editing. All authors read and approved the manuscript. This manuscript has not been accepted or published elsewhere.

## 8 Acknowledgements

The authors thank the members of the Max Planck Unit for the Science of Pathogens, in particular Prof. Kürşad Turgay, Fabian Cornejo, Dmitriy Ignatov, Gita Naseri and Gisela Klauck, for their scientific contribution to the project and their critical reading of the manuscript. The authors acknowledge Prof. Michael Federle of the University of Illinois Chicago (USA) for kindly providing our laboratory with the p7INT plasmid.

## 9 Data Availability Statement

The nanopore sequencing data generated for this study have been deposited at the European Nucleotide Archive (ENA) under accession PRJEB72852. The source code for the identification of structural and single nucleotide variants based on the nanopore sequencing data has been made available on Github under the following link: https://github.com/MPUSP/snakemake-ont-bacterial-variants.

The external RNA-seq datasets analyzed for the identification of transcriptionally silent sites in *S. pyogenes* SF370 and M1T1 5448 are partially available in the European Nucleotide Archive (ENA) or in the gene expression omnibus (GEO) via the following SRP /DRP accession numbers: SRP066922, DRP008790, SRP389502, DRP007263, SRP267980, SRP266954, DRP004959, SRP073901, SRP119757 and SRP390529 or the NCBI BioProject identifiers: PRJNA304766, PRJDB13957, PRJDB8873, PRJNA640594, PRJNA638918, PRJDB8158, PRJNA319617, PRJNA412519, PRJNA297518. Furthermore, datasets published by our laboratory were included, accessible under GEO accession numbers GSE84641, GSE40198 or BioProject accession PRJNA193607; at NCBI under accession number SRP149896. The source code/ pipeline for the identification of transcriptionally silent sites has been made available on GitHub under the following link: https://github.com/MPUSP/lautenschlaeger_silent_sites.

