## Supplementary Material for "Expanding the genetic toolbox for the obligate human pathogen *Streptococcus pyogenes*"

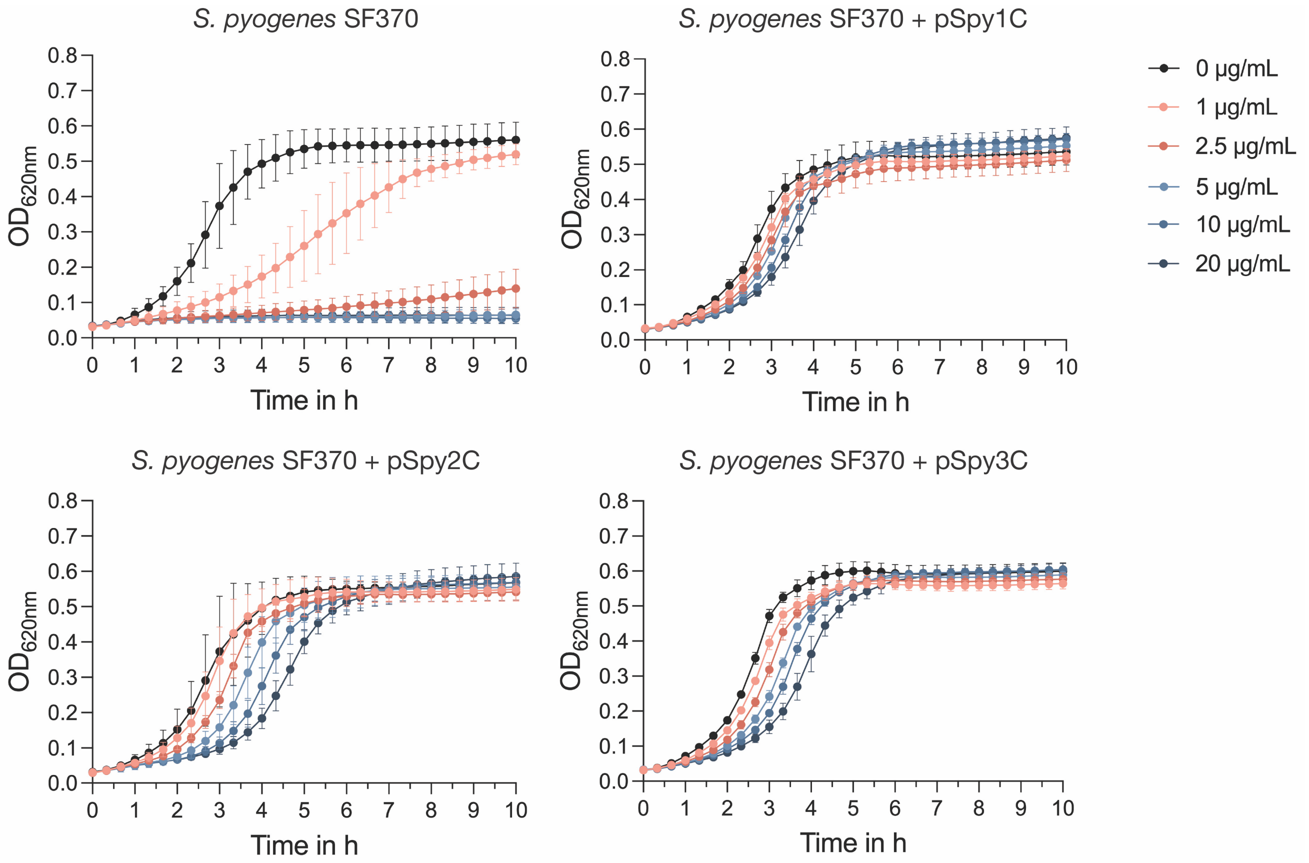


**Supplementary Figure 1.** Adjustments of chloramphenicol concentrations for the maintenance of replicative plasmids in *S. pyogenes* (EC2514) in THY medium. Growth in $OD_{620nm}$ is shown for the *S. pyogenes* SF370 wildtype strain (top left) and the wildtype strain transformed with the replicative plasmids pSpy1C (top right), pSpy2C (bottom left) and pSpy3C (bottom right). Chloramphenicol was added to the cultures at the beginning of growth (t=0) at different concentrations (1 µg/mL, 2.5 µg/mL, 5 µg/mL, 10 µg/mL and 20 µg/mL), indicated by the legend at the top right. Ethanol served as a control (0 µg/mL). Experiments were performed in biological triplicates with technical replicates each.

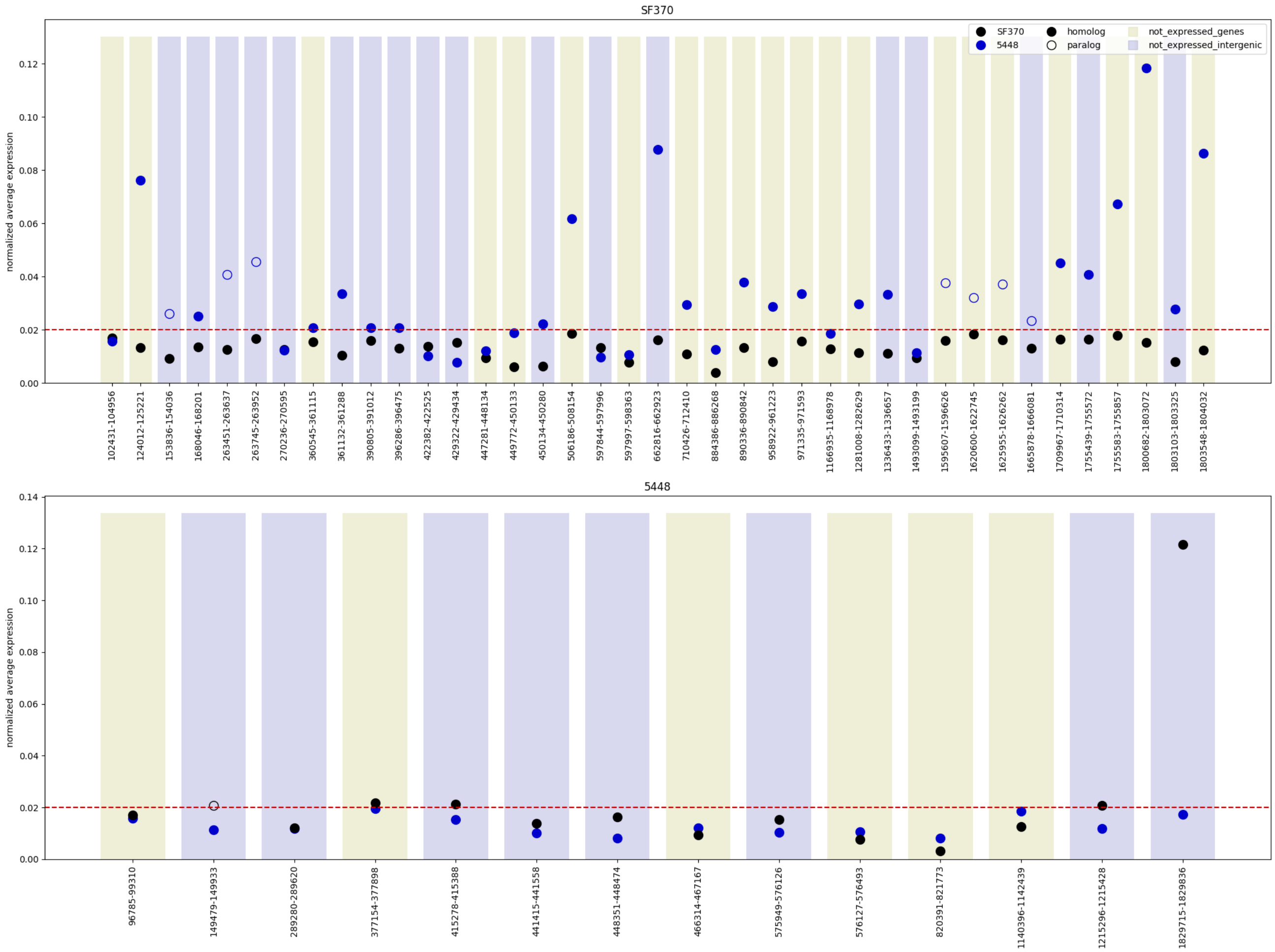


**Supplementary Figure 2:** Overview of the transcriptionally silent sites identified via bioinformatic analysis of publicly available RNA-seq datasets on *S. pyogenes* SF370 and M1T1 5448. Regions including genes of SF370 (black data points) or M1T1 5448 (blue data points) showing expression levels below the threshold (normalized average expression ≤0.02, highlighted by a red dashed line) are shown on a light green background, while intergenic regions are indicated on a lilac background. The top graph uses the SF370 genome as a reference and displays, which transcriptionally silent regions in SF370 (black data points, filled), may also be found in M1T1 5448 (blue data points) and whether this region is homologous (filled data point) or paralogous (data point without fill). The bottom graph displays all transcriptionally silent regions found for M1T1 5448 and whether they can also be found in SF370.

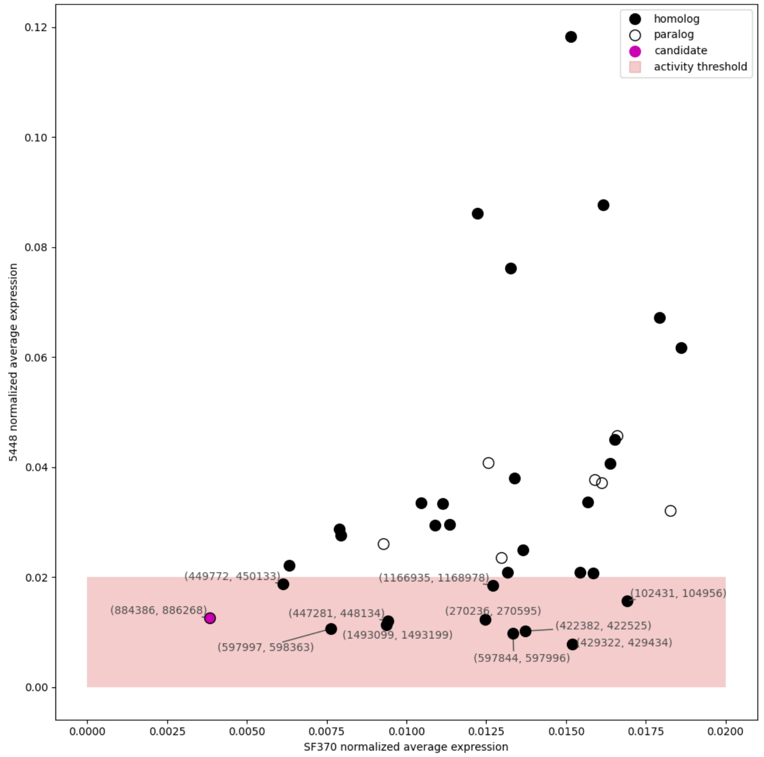


**Supplementary Figure 3:** Transcriptionally silent sites within the transcriptional activity threshold in *S. pyogenes* SF370 and M1T1 5448. Regions with a normalized average expression ≤0.02 in both strains are shown within the red area and the genomic coordinates are specified. The candidate region, which was used to design the integrative plasmid pSpy0K6, is highlighted in pink. Regions with homology between SF370 and M1T1 5448 are indicated by a filled data point, while paralogs are shown by data points without fill.

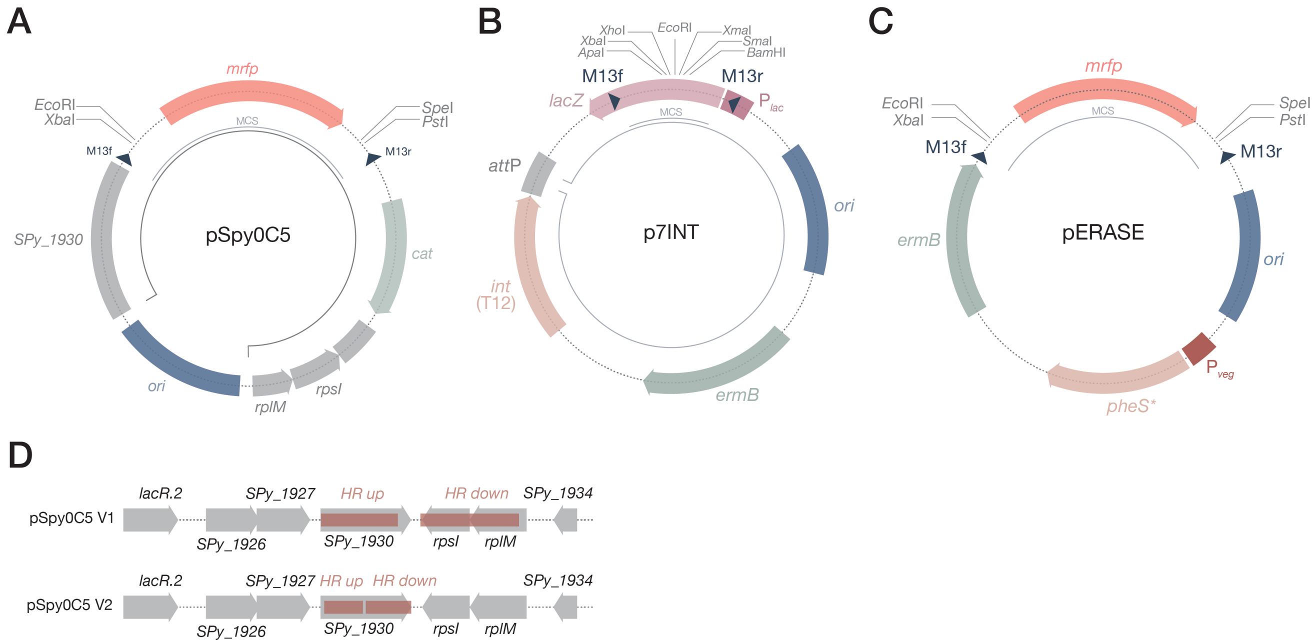


**Supplementary Figure 4.** Plasmid maps of the integrative plasmids pSpy0C5, p7INT and the genome editing plasmid pERASE. **(A)** Map of the pSpy0C5 version 1 plasmid. The pUC replicon for propagation in *E. coli* is marked in blue. The resistance cassette for chloramphenicol (*cat*) is highlighted in sage green. For pSpy0C5, DNA sequences with homology to the genome of SF370 are shown in grey. Restriction sites of the MCS flank the *mrfp* reporter gene under control of $P_{lac(Eco)}$, enabling red-white screening of clones containing the desired insert in *E. coli*. Small dark blue triangles indicate the binding sites for the M13f and M13r standard primers. The inner black circular line marks the part of the plasmid that will integrate into the streptococcal genome. **(B)** Plasmid map of the integrase-based plasmid p7INT. The MCS features a *lacZ* reporter gene, highlighted in pink, allowing for blue-white screening in *E. coli*. The integrase gene *int* and the respective *attP* site derived from bacteriophage T12 are highlighted in pale pink and grey, respectively. The pUC replicon for propagation in *E. coli* is marked in blue. The resistance cassette for erythromycin (*ermB*) is highlighted in sage green. The inner black circular line marks the part of the plasmid that will integrate into the streptococcal genome. **(C)** Map of the pERASE plasmid used to generate scarless gene deletions in *S. pyogenes*. pERASE contains the *mrfp* reporter for red-white screening in *E. coli* (salmon red) and the *pheS** counterselection marker (pale pink) allowing to screen for plasmid loss, and thus, deletion of the gene on 5 mM PCPA (4-Chlor-DL-Phenylalanine) agar plates. **(D)** Genomic context of the locus in the *S. pyogenes* SF370 genome targeted for integration by pSpy0C5 V1 (as in Supplementary Figure A) and V2. Sites selected to enable homologous recombination of pSpy0C5 are highlighted in red. All attempts to transform both versions of the plasmid were unsuccessful.

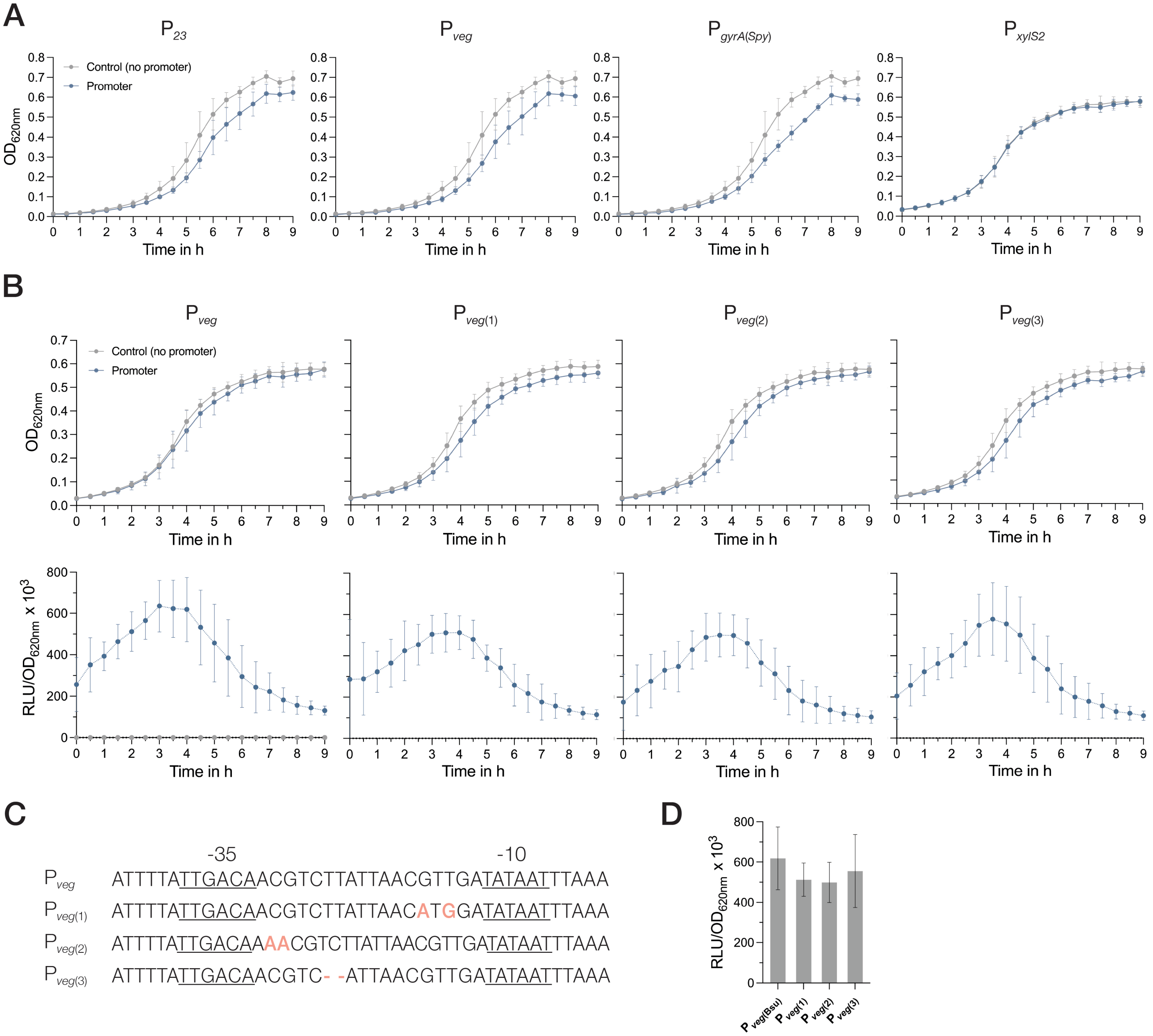


**Supplementary Figure 5.** *S. pyogenes* growth (solid lines) and luminescence (dotted lines) signal of the luciferase reporter strains harboring different constitutive promoters (blue) in comparison to the control without promoter (grey). **(A)** Growth (shown in $OD_{620nm}$) of *S. pyogenes* strains harboring either the control plasmid (pSpy1C-*ffluc*) or pSpy1C featuring the fusions of different constitutive promoters to the luciferase reporter *ffluc* (e.g. pSpy1C-$P_{23}$*_ffluc*). **(B)** Growth, shown in $OD_{620nm}$ (top panel), and promoter activities, shown in relative luminescence units normalized by optical density ($RLU/OD_{620nm}$) (bottom panel) of strains harboring luciferase reporter plasmids with different variants of the $P_{veg}$ promoter (1-3). **(C)** DNA sequences of the native $P_{veg}$ promoter and its derivatives (1-3) including the -35 region, the spacer and the -10 region. The respective changes to the spacer sequence to obtain $P_{veg(1)}$ (G-to-A and T-to-G mutation), $P_{veg(2)}$ (insertion of two adenines) and $P_{veg(3)}$ (deletion of two thymines) are highlighted in apricot. **(D)** Comparison of the mean promoter activities ($RLU/OD_{620nm}$) of the native $P_{veg}$ promoter and its derivatives $P_{veg(1)}$, $P_{veg(2)}$, and $P_{veg(3)}$ after 4 hours of growth in THY. Experiments were performed in biological triplicates and measurements in technical duplicates.

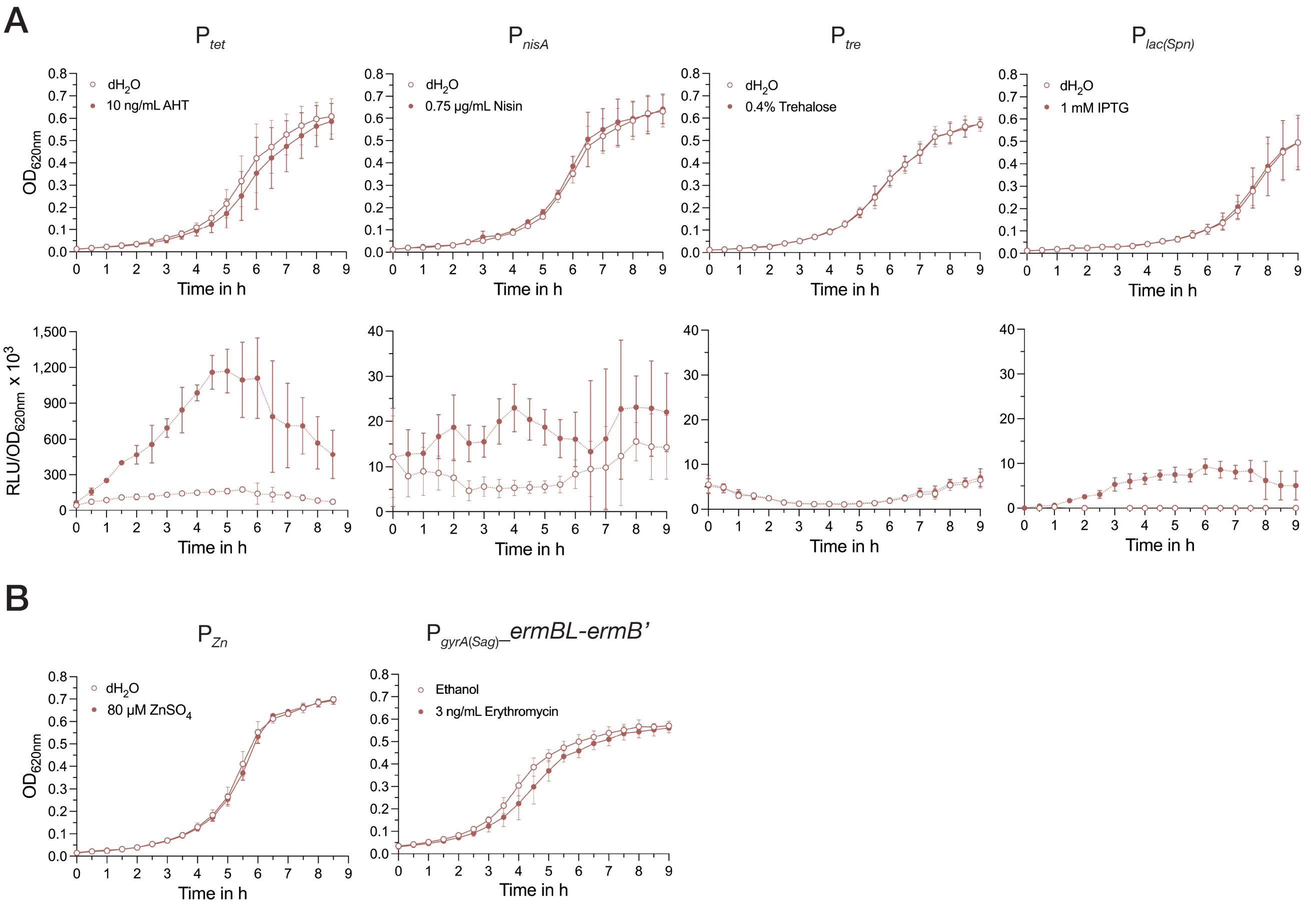


**Supplementary Figure 6.** Growth (solid lines) and luminescence signal (dotted lines) of the *S. pyogenes* reporter strains featuring different inducible promoters. **(A)** Growth of the reporter plasmids harboring the inducible promoter systems $P_{tet}$, $P_{nisA}$, $P_{lac(Spn)}$ and $P_{tre}$ is shown in $OD_{620nm}$ over time in the top row, while the luminescence signal is shown in relative luminescence units normalized by optical density ($RLU/OD_{620nm}$) in the bottom row. Cultures were induced with the respective inducer compounds or the control at the beginning of growth (t=0) ($P_{tet}$: 10 ng/mL anhydrotetracycline, $P_{nisA}$: 0.75 µg/mL nisin, $P_{tre}$: 0.4% trehalose, $P_{lac(Spn)}$: 1 mM IPTG, distilled water served as negative control). Data are shown for the uninduced condition (data points with white fill) and the induced condition (data points with red fill). **(B)** Growth of the strains harboring reporter plasmids with the inducible regulatory systems $P_{Zn}$ and $P_{gyrA(Sag)}$*_ermBL-ermB'* shown as $OD_{620nm}$ over time in the absence or presence of the respective inducer compounds ($P_{Zn}$: 80 µM $ZnSO_4$ and $P_{gyrA(Sag)}$*_ermBL-ermB'*: 3 ng/mL erythromycin). Data are shown for the uninduced condition (data points with white fill) and the induced condition (data points with red fill). Distilled water served as a negative control for $P_{Zn}$ induction and ethanol was used as a negative control for $P_{gyrA(Sag)}$*_ermBL-ermB'*. Experiments were performed in biological triplicates and measurements in technical duplicates.

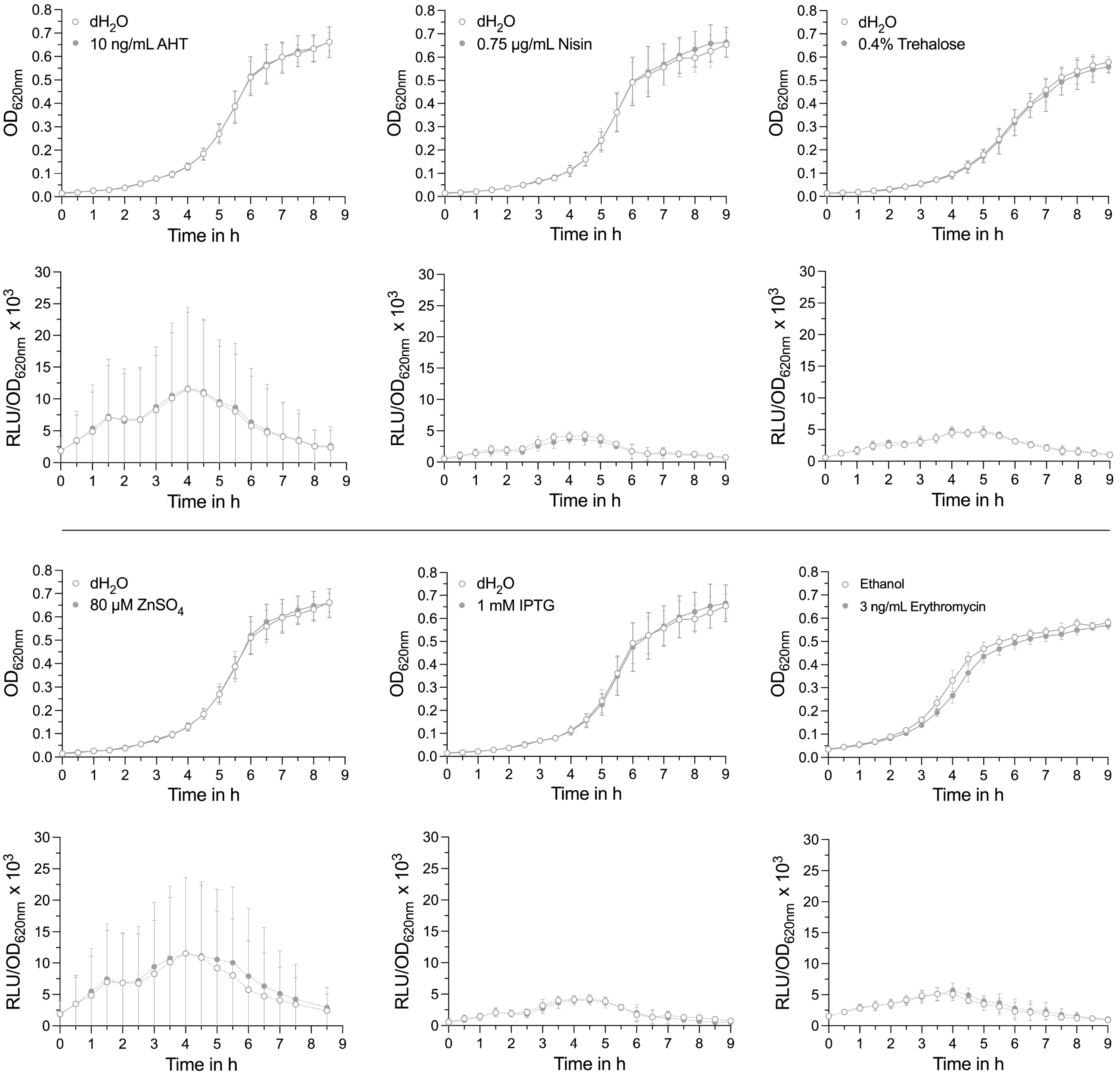


**Supplementary Figure 7.** Growth (solid lines) and luminescence signal (dotted lines) of the *S. pyogenes* control strains harboring pSpy1C-*ffluc* (no promoter) in the presence (data points with grey fill) and absence (data points with white fill) of different inducer compounds ($P_{tet}$: 10 ng/mL anhydrotetracycline, $P_{nisA}$: 0.75 µg/mL nisin, $P_{tre}$: 0.4% trehalose, $P_{Zn}$: 80 µM $ZnSO_4$, $P_{lac(Spn)}$: 1 mM IPTG, $P_{gyrA(Sag)}$*_ermBL-ermB'*: 3 ng/mL erythromycin). Distilled water ($dH_2O$) or ethanol were used as negative controls for induction. Growth is shown in $OD_{620nm}$ over time in the first and third row. The luminescence signal is shown in relative luminescence units normalized by optical density ($RLU/OD_{620nm}$) in the second and fourth row. Experiments were performed in biological triplicates and measurements in technical duplicates.

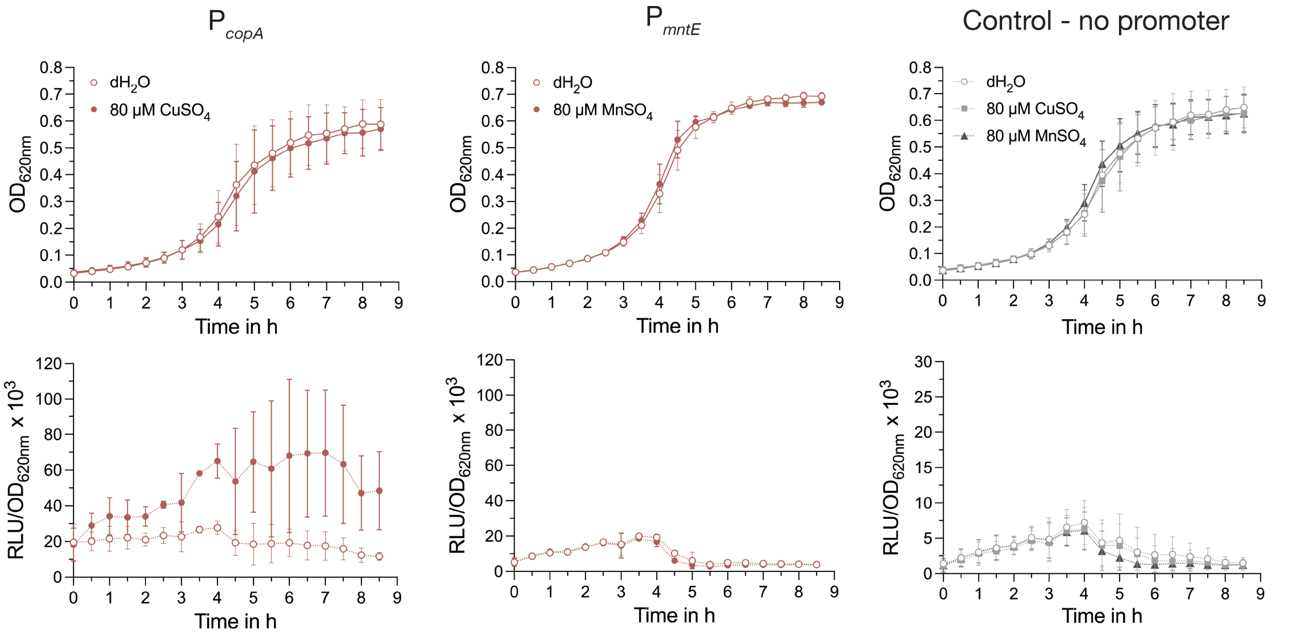


**Supplementary Figure 8.** Promoter activities of the metal-inducible promoters P*copA* and P*mntE* in the presence or absence of $CuSO_4$ and $MnSO_4$. Growth in $OD_{620nm}$ (top row and solid lines) and luminescence signal in relative luminescence units normalized by optical density ($RLU/OD_{620nm}$) (bottom row and dotted lines) of the *S. pyogenes* reporter strains harboring the metal-inducible promoters P*copA* or P*mntE* or the control (no promoter) in the presence (data points with red or grey fill) and absence (data points with white fill) of different inducer compounds (80 µM $MnSO_4$ and 80 µM $CuSO_4$). Distilled water ($dH_2O$) was used as a negative control for induction. Experiments were performed in biological triplicates and measurements in technical duplicates.

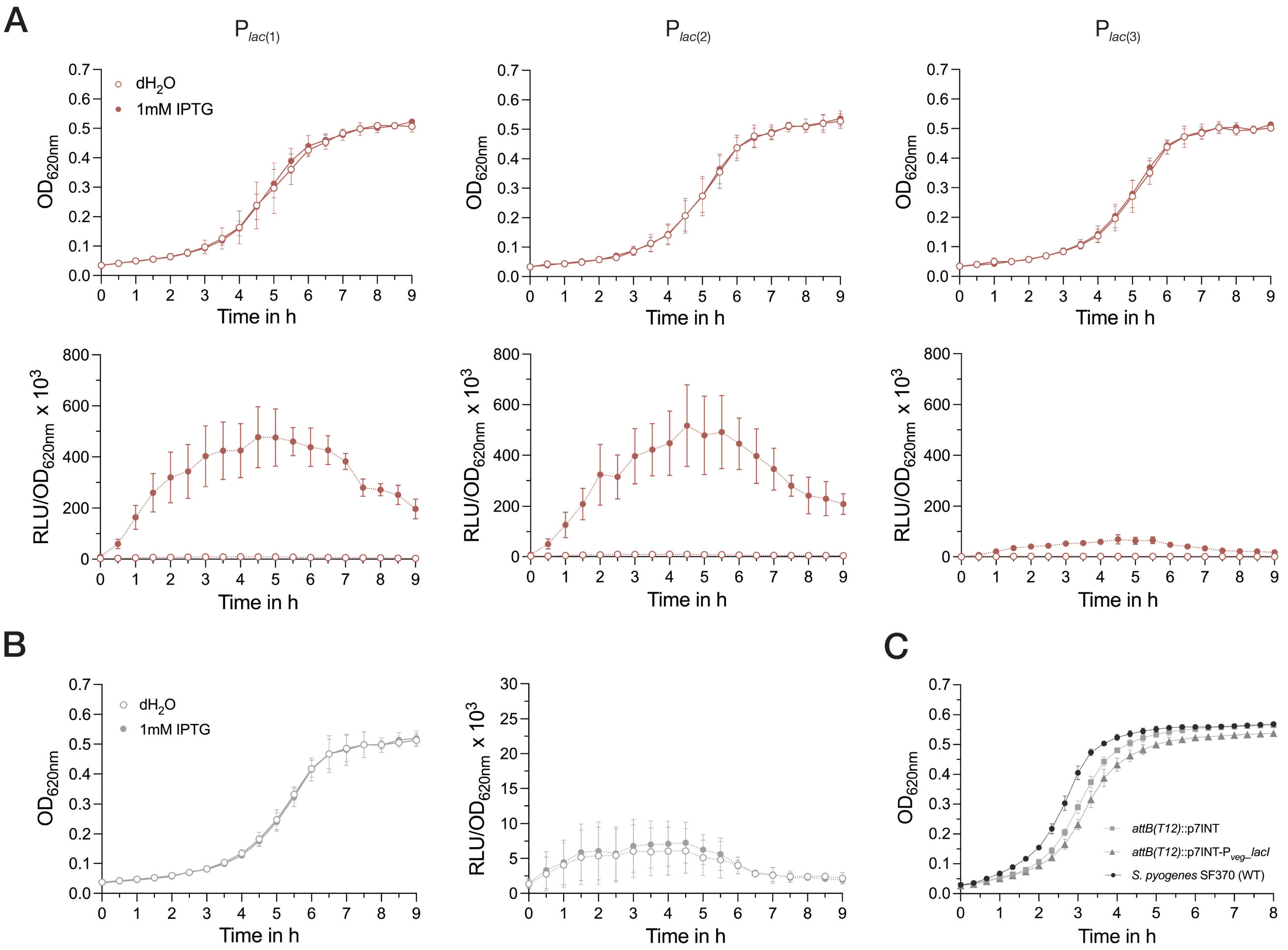


**Supplementary Figure 9.** Activity of IPTG-inducible $P_{lac(Spn)}$ promoters harboring different modifications within the translation initiation region in a LacI-expressing *S. pyogenes* strain. **(A)** *S. pyogenes* growth (solid lines) and luminescence signal (dotted lines) of the reporter strains harboring different metal-inducible promoters and the control (no promoter) in the presence (data points with red fill) and absence (data points with white fill) of 1 mM IPTG. Distilled water ($dH_2O$) was used as a negative control for induction. Growth is shown in $OD_{620nm}$ over time (top row), while the luminescence signal is shown in relative luminescence units normalized by optical density ($RLU/OD_{620nm}$) (bottom row). **(B)** Growth shown in $OD_{620nm}$ (left) and luminescence signal depicted in $RLU/OD_{620nm}$ (right) for the control strain harboring the reporter plasmid without a promoter in the presence (data points with grey fill) and absence (data points with white fill) of 1 mM IPTG. **(C)** Growth curves of the LacI expression strain (EC3732) compared to *S. pyogenes* wildtype (EC2514) and *S. pyogenes* containing the empty p7INT plasmid (EC3241). Experiments were performed in biological triplicates and measurements in technical duplicates.

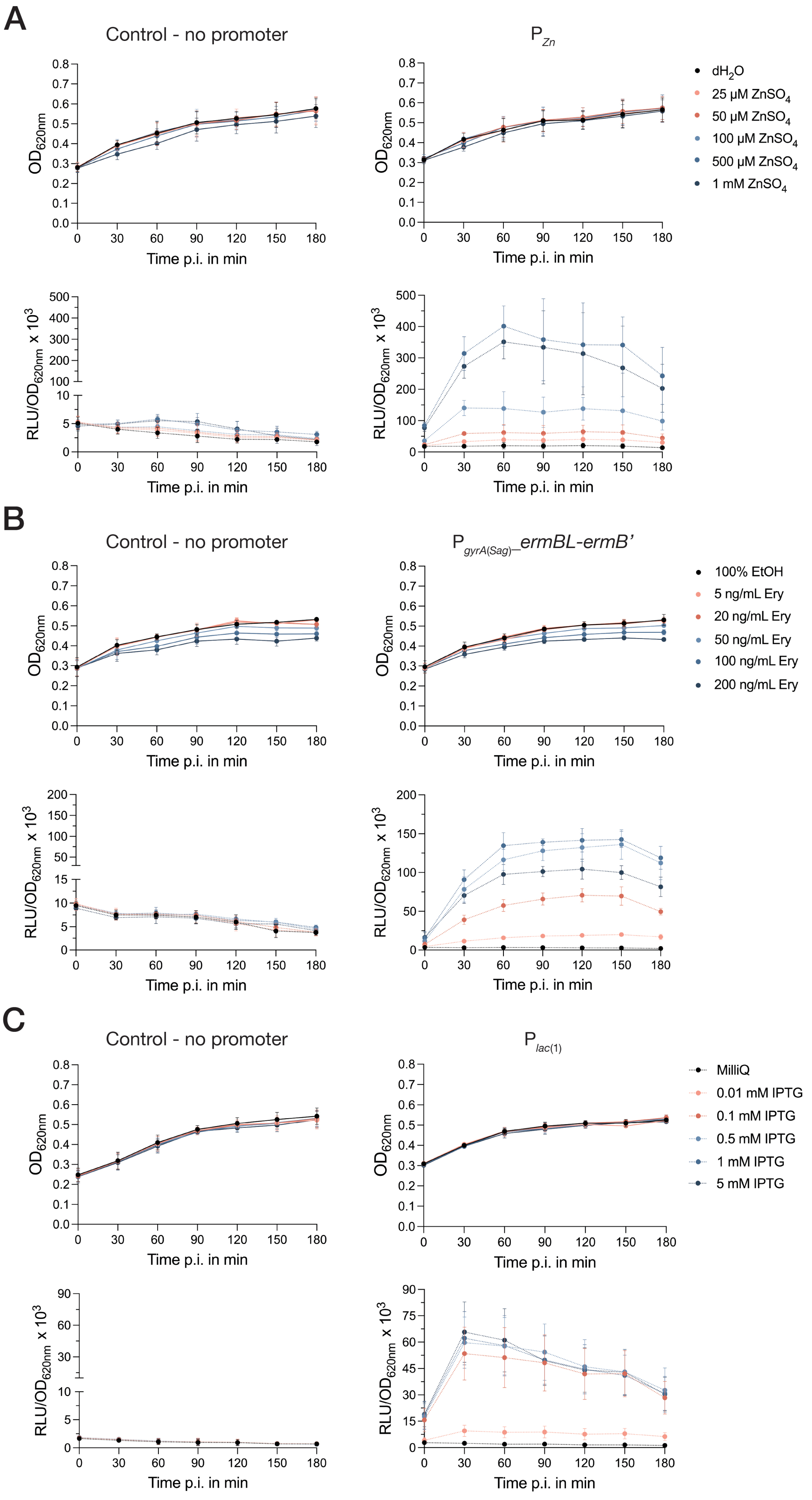
A
Control - no promoter
P Zn
OD620nm
Time p.i. in min
RLU/OD620nm x 10^3
dH2O
25 µM ZnSO4
50 µM ZnSO4
100 µM ZnSO4
500 µM ZnSO4
1 mM ZnSO4
B
Control - no promoter
P gyrA(Sag)–ermBL-ermB'
100% EtOH
5 ng/mL Ery
20 ng/mL Ery
50 ng/mL Ery
100 ng/mL Ery
200 ng/mL Ery
C
Control - no promoter
P lac(1)
MilliQ
0.01 mM IPTG
0.1 mM IPTG
0.5 mM IPTG
1 mM IPTG
5 mM IPTG

**Supplementary Figure 10.** Growth and luminescence signal of the three inducible systems $P_{Zn}$, $P_{gyrA(Sag)\_}$*ermBL-ermB'* and $P_{lac(1)}$ in response to increasing inducer concentrations. The legend to the right of each panel shows the applied inducer concentrations. Growth and luminescence signals are shown in $OD_{620nm}$ or relative luminescence units normalized by optical density ($RLU/OD_{620nm}$) on the y-axis, while the time post induction (p.i.) is indicated on the x-axis. **(A)** *S. pyogenes* growth (top row and solid lines) and luminescence signal (bottom row and dotted lines) of the strain harboring either the control plasmid without promoter (left) or the zinc-inducible promoter (right). **(B)** *S. pyogenes* growth (top row and solid lines) and luminescence signal (bottom row and dotted lines) of the strain harboring either the control plasmid without promoter (left) or the erythromycin-inducible riboswitch (right). **(C)** *S. pyogenes* growth (top row and solid lines) and luminescence signal (bottom row and dotted lines) of the strain harboring either the control plasmid without promoter (left) or the reporter plasmid with the IPTG-inducible $P_{lac(1)}$ promoter (right). Experiments were performed in biological triplicates and measurements in technical duplicates.

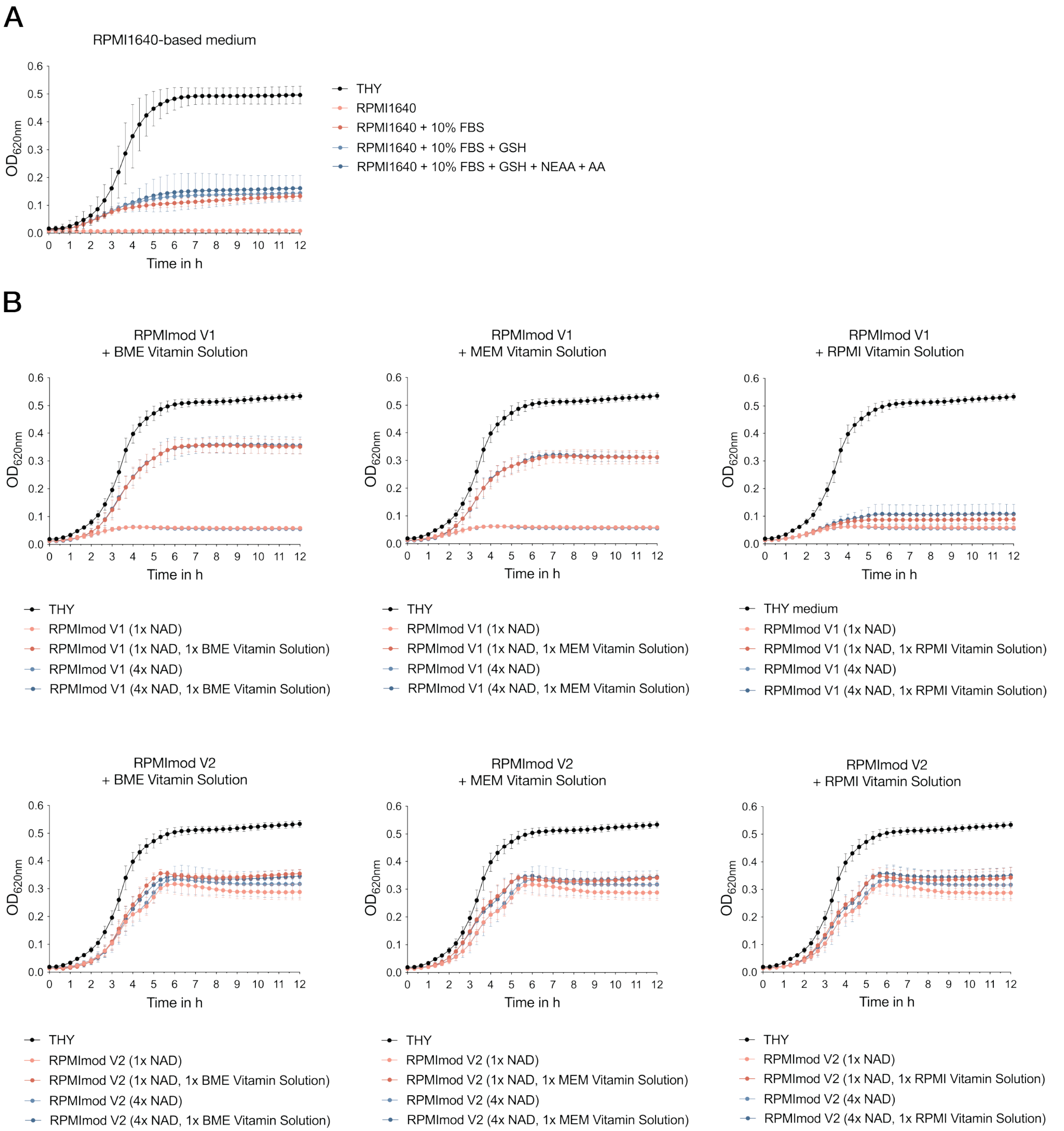


**Supplementary Figure 11.** Growth of *S. pyogenes* SF370 in different formulations of the RPMI4Spy chemically defined medium. The growth is shown in $OD_{620nm}$ on the y-axis, while the time in hours is indicated on the x-axis. **(A)** Initial test experiment using RPMI1640 cell culture medium supplemented with glutathione, fetal bovine serum, RPMI non-essential amino acid solution or RPMI amino acid solution or combinations thereof. Abbreviations: GSH = Glutathione, NEAA = non-essential amino acids, AA = amino acids, FBS = fetal bovine serum **(B)** Test experiments with different RPMI4Spy medium formulations supplemented with either the commercial RPMI amino acid solution (V1) or a self-prepared amino acid mix (V2). Both basic formulations were further supplemented with different vitamin solutions (BME Vitamin Solution, MEM Vitamin Solution or RPMI Vitamin Solution) and two different concentrations of niacinamide (NAD). Experiments were performed in biological triplicates. Abbreviations: RPMImod = modified RPMI1640.

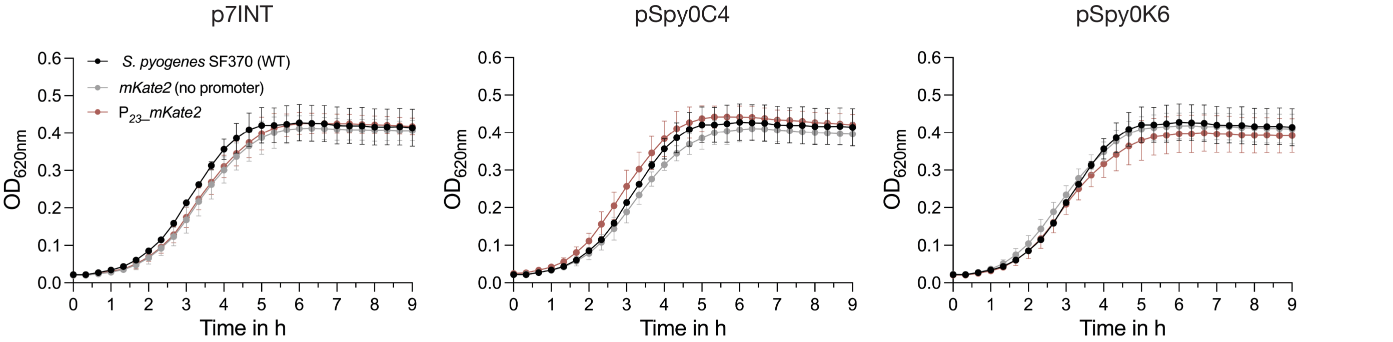


**Supplementary Figure 12.** Growth of strains harboring either the mKate2 reporter ($P_{23}$*_mKate2*, EC3391) or the negative control without a promoter (no promoter, EC3480) integrated at different genomic locations compared to the *S. pyogenes* wildtype (EC2514). The reporter was integrated at the tmRNA locus (*attB*(T12)) using p7INT (left), into the *sagB* gene using pSpy0C4 (center) and into a transcriptionally silent open reading frame (*SPy_1078*) using pSpy0K6 (right). The growth is shown in $OD_{620nm}$ on the y-axis, while the time in hours is indicated on the x-axis. Experiments were performed in biological triplicates and each measurement in technical duplicates.

**Supplementary Table 1.** Bacterial strains used and created in this study. Symbols: (~) indicates translational fusions; (*) indicates an introduced stop codon (TAA)

| Code | Strain Name | Features | Source | Alias Used |
|---|---|---|---|---|
| | | **Wildtype strains** | | |
| EC1 | DH5*α* | Cloning host: *E. coli* K12 *fhuA2* Δ(*argF-lacZ*)U169 *phoA glnV44* Φ80 Δ(*lacZ*)M15 *gyrA96 recA1 relA1 endA1 thi-1 hsdR17* | New England Biolabs | |
| RDN204 | OneShot™ TOP10 | F-*mcrA* Δ( *mrr-hsd*RMS-*mcr*BC) Φ80*Lac*ZΔM15 Δ *Lac*X74 *rec*A1 *ara*D139 Δ( *araleu*) 7697 *gal*U *gal*K *rps*L (StrR) *end*A1 *nup*G | Invitrogen | |
| EC2621 | BW25113 *lon*::*kanR* | *Δ(araD-araB)567 ΔlacZ4787*(::rrnB-3) *λ- rph-1 Δ(rhaD-rhaB)568 hsdR514 lon*::*kanR* | Hansen *et al.*, 2012 (1) | |
| EC2705 | *Bacillus subtilis* PY79 | wildtype, *trpC2* prototroph | Youngman *et al.*, 1984 (2) | |
| EC2514 | *S. pyogenes* SF370 | wildtype M1 strain | *Streptococcus pyogenes* Rosenbach ATCC700294 | wildtype |
| | | **Strains with empty integrative plasmids** | | |
| EC3217-3218 | EC2514 *amyA*::pSpy0C2 | *amyA*::pSpy0C2 | this study | n/a |
| EC3220-3221 | EC2514 *sagB*::pSpy0C4 | *sagB*::pSpy0C4 | this study | n/a |
| EC3729-3730 | EC2514 *Spy_1078*::pSpy0K6 | SP*y_1078*::pSpy0K6 | this study | n/a |
| EC3241-3242 | EC2514 *attB*(T12)::p7INT | *attB*(T12)::p7INT (integration does not impair functionality) | this study | n/a |
| EC3741-3742 | EC2514 *attB*(T12)::p7INT.1 | *attB*(T12)::p7INT.1 | this study | n/a |
| | | **Reporter strains** | | |
| EC3364-3365 | EC2514 *attB*(T12)::p7INT-$P_{23}$*_mNeongreen~ssrA* | *attB*(T12)::p7INT-$P_{23}$*_mNeongreen~ssrA,* $Erm^R$ | this study | native SsrA tag |
| EC3385-3386 | EC2514 *attB*(T12)::p7INT-$P_{23}$*_mNeongreen*~ssrA* | *attB*(T12)::p7INT-$P_{23}$*_mNeongreen*~ssrA,* $Erm^R$ | this study | mNeongreen reporter (untagged) |
| EC3388-3389 | EC2514 *attB*(T12)::p7INT- | *attB*(T12)::p7INT-$P_{23}$*_mNeongreen~ssrA* (LAA to LDD), $Erm^R$ | this study | LDD variant |

| Code | Strain Name | Features | Source | Alias Used |
|---|---|---|---|---|
| | $P_{23}$_*mNeongreen~ssrA* (LDD) | | | |
| EC3486-3487 | EC2514 *attB*(T12)::p7INT-$P_{23}$_*mNeongreen~ssrA* (ASV) | *attB*(T12)::p7INT-$P_{23}$_*mNeongreen~ssrA* (LAA to ASV), $Erm^R$ | this study | ASV variant |
| EC3489-3490 | EC2514 *attB*(T12)::p7INT-*mNeongreen*~STOP *~ssrA* | *attB*(T12)::p7INT-*mNeongreen*~ssrA,* $Erm^R$ | this study | mNeongreen control |
| EC3391-3392 | EC2514 *attB*(T12)::p7INT-$P_{23}$_*mKate2*~ssrA* | *attB*(T12)::p7INT-$P_{23}$_*mKate2*~ssrA,* $Erm^R$ | this study | mKate2 reporter |
| EC3480-3481 | EC2514 *attB*(T12)::p7INT-*mKate*~ssrA* | *attB*(T12)::p7INT-*mKate*~ssrA,* $Erm^R$ | this study | mKate2 control |
| EC3735-3736 | EC2514 *SPy_1078*::pSpy0K6-$P_{23}$_*mKate2*~ssrA* | *SPy_1078*::pSpy0K6-$P_{23}$_*mKate2*~ssrA*, $Kan^R$ | this study | n/a |
| EC3738-3739 | EC2514 *SPy_1078*::pSpy0K6-*mKate2*~ssrA* | *SPy_1078*::pSpy0K6-*mKate2*~ssrA*, $Kan^R$ | this study | n/a |
| EC3429-3430 | EC2514 *sagB*::pSpy0C4-$P_{23}$_*mKate*~ssrA* | *sagB*::pSpy0C4-$P_{23}$_*mKate*~ssrA*, $Cm^R$ | this study | n/a |
| EC3492-3493 | EC2514 *sagB*::pSpy0C4-*mKate*~ssrA* | *sagB*::pSpy0C4-*mKate*~ssrA*, $Cm^R$ | this study | n/a |
| | | **Other** | | |
| EC3732-3733 | EC2514 *attB*(T12)::p7INT-$P_{veg}$_*lacI* | *attB*(T12)::p7INT-$P_{veg}$_*lacI*, $Erm^R$ | this study | LacI expression strain |
| EC3756-3757 | EC2514 *ΔsagA* | *ΔsagA* | this study | *sagA* mutant |

**Supplementary Table 2.** Oligonucleotides used in this study. Overhangs used for cloning purposes are underlined.

| Oligo Code | Name | 5'-3'-Sequence |
|---|---|---|
| OLEC290 | *mga*_F | TTAACCTCTGTTTGATTCGC |
| OLEC291 | *mga*_R | GTCGTACTGACTTAACGAAA |
| OLEC4854 | *ropB*_F | AGCGACTATCATCCGAAACAT |
| OLEC4855 | *ropB*_R | GCCCTGGAGCTGTTGAGATA |
| OLEC4856 | *csrRS*_F | TCGCTAGAAGACTATTTGACCAT |
| OLEC4857 | *csrRS*_R | AAGACATCGCGATTGACAGT |
| OLEC9562 | *ffluc*_fwd | GATCGGTCTCAGCTGAATTCAAAAAAAAATCTAGAAGGAGGAATAAAAAAATGGAAGACG |
| OLEC9563 | *ffluc*_rev | GATCGGTCTCTTCACTGCAGTACTAGTATTACAATTTGGACTTTCCGCCC |
| OLEC9589 | $P_{23}$_goldenbrick_fwd | GATCGGTCTCAGCTGAATTCAAAAAAAAATCTAGACTCGAAAAGCCCTGACAACCC |
| OLEC9591 | $P_{gyrA}$_goldenbrick_fwd | GATCGGTCTCAGCTGAATTCAAAAAAAAATCTAGATTTTTGTGTTGTTTAGTTGC |
| OLEC9592 | $P_{gyrA}$_goldenbrick_rev | GATCGGTCTCTTCACTGCAGTACTAGTATTCTAGTCAATACTAGATTTC |
| OLEC9595 | $P_{tet}$_goldenbrick_fwd | GATCGGTCTCAGCTGAATTCAAAAAAAAATCTAGAGATAATGCCGACTGTACTTTTTAC |
| OLEC9596 | $P_{tet}$_goldenbrick_rev | GATCGGTCTCTTCACTGCAGTACTAGTACTTAACTAGACTCGAAGATC |
| OLEC9749 | $P_{veg}$_GBOH_fwd | GATCGGTCTCAGCTGAATTCAAAAAAAAATCTAGAGAGTTCTGAGAATTGGTATGC |
| OLEC9750 | $P_{veg}$_GBOH_rev | GATCGGTCTCTTCACTGCAGTACTAGTAACTACATTTATTGTACAACACGAGC |
| OLEC9751 | $P_{23}$_GBOH_rev | GATCGGTCTCTTCACTGCAGTACTAGTACCAACATCATTGTCATTCATATTTTTC |
| OLEC9773 | pIB166_GG_M13_fwd | GATCCGTCTCTATGTGTCATAGCTGTTTCCTGCAAAGAATGGTGATGACAATTG |
| OLEC9774 | pIB166_GG_M13_rev | GATCCGTCTCTGCTGACTGGCCGTCGTTTTACCATAAACACCAATAGCCTTAAC |
| OLEC9775 | MCS_GG_fwd | GATCCGTCTCCCAGCGAGACCAGCTGAATTCAAAAA |
| OLEC9776 | MCS_GG_rev | GATCCGTCTCGACATGAGACCTTCACTGCAGTACTAG |
| OLEC9777 | pIB184_GG_M13_fwd | GATCCGTCTCTATGTGTCATAGCTGTTTCCTGGACTAGCAAATACTAACAACAAGACAC |
| OLEC9778 | pIB184_GG_M13_rev | GATCCGTCTCTGCTGACTGGCCGTCGTTTTACGACGCTCTTCCGCTTCCTC |
| OLEC9779 | pIB185_GG_M13_fwd | GATCCGTCTCTATGTGTCATAGCTGTTTCCTGGATTCTAGACGCTCTTCCG |
| OLEC9780 | pIB185_GG_M13_rev | GATCCGTCTCTGCTGACTGGCCGTCGTTTTACCAACGTTAATAAGACGTTGTC |

| Oligo Code | Name | 5'-3'-Sequence |
|---|---|---|
| OLEC9781 | *mrfp*_GBOH_fwd | GATCGAGACCAGCTGAATTCAAAAAAAAATCTAGATGGCCGGCGATTCATTAATGCAGCTGGC |
| OLEC9782 | *mrfp*_GBOH_rev | GATCGAGACCTTCACTGCAGTACTAGTATTAACCGGTTATAAACGCAGAAAGGCCCAC |
| OLEC9799 | pIB184_*Esp*3Imut_fwd | CTTTTTGAACCCGTCGCCTTACGGCTTTATTAGATATGTAATC |
| OLEC9800 | pIB184_*Esp*3Imut_rev | CCGTAAGGCGACGGGTTCAAAAAGGTTTAAATAAAGGAGA |
| OLEC9801 | pIB166_*Esp*3Imut_fwd | AACGTCACAGAAACGATTTTCAGACGTTTTAATAAAAAATC |
| OLEC9802 | pIB166_*Esp*3Imut_rev | GTCTGAAAATCGTTTCTGTGACGTTTTAGCGTTTATTTCGTTTAG |
| OLEC9824 | kanR_GG_fwd | GATCCGTCTCTATGTGTCATAGCTGTTTCCTGCCTAATAATTTCTAGGTACTAAAAC |
| OLEC9825 | kanR_GG_rev | GATCCGTCTCGTTAGCCTTATAGCTTGTAAATTCTATC |
| OLEC9826 | pIB166_GG2_fwd | GATCCGTCTCTCTAATATGAGATAATGCCGACTGTAC |
| OLEC9827 | ermR_GG_fwd | GATCCGTCTCTATGTGTCATAGCTGTTTCCTGCGAAATGATACACCAATCAG |
| OLEC9828 | ermR_GG_rev | GATCCGTCTCGTTAGCTTGCATATGATCCGAGCTTC |
| OLEC9829 | pBAV1K-T5_fwd | GATCCGTCTCTATGTGTCATAGCTGTTTCCTGCTAGACCTAGTGTCATTTTATTTC |
| OLEC9830 | pBAV1K-T5_rev | GATCCGTCTCTGCTGACTGGCCGTCGTTTTACGTTATACGCCAACTTTGAAAAC |
| OLEC9831 | pBAV1K-T5_mut_fwd | ACGTCTGAGAAACGATTTTCAGACGTTTTAATAAAAAATC |
| OLEC9832 | pBAV1K-T5_mut_rev | ACGTCTGAAAATCGTTTCTCAGACGTTTTAGCGTTTATTTC |
| OLEC9833 | pBAV1K-T5_GG2_ fwd | GATCCGTCTCCCAGCGTTTTCAAAGTTGGCGTATAAC |
| OLEC9834 | pBAV1K-T5_GG2_rev | GATCCGTCTCGACATTACTGGATGAATTGTTTTAGTAC |
| OLEC9835 | cmR_GG2_fwd | GATCCGTCTCTATGTCAAAGAATGGTGATGACAATTG |
| OLEC9836 | cmR_GG2_rev | GATCCGTCTCTGCTGTTATAAAAGCCAGTCATTAGGC |
| OLEC9837 | ermR_GG2_fwd | GATCCGTCTCTATGTCGAAATGATACACCAATCAG |
| OLEC9838 | ermR_GG2_fwd | GATCCGTCTCTGCTGCTTGCATATGATCCGAGCTTC |
| OLEC9875 | pIB185_*Esp*3Imut_fwd | CTGGATTACATATCTAATAAAGCCGTAAGAAGACGGGTTC |
| OLEC9876 | pIB185_*Esp*3Imut_rev | CTTTATTTAAACCTTTTTGAACCCGTCTTCTTACGGCTTTA |
| OLEC9877 | pIB185new_fwd | GATCCGTCTCCCAGCGAAGCTCGGATCATATGCAAG |
| OLEC9878 | pIB185new_rev | GATCCGTCTCGACATCTGATTGGTGTATCATTTCG |
| OLEC10024 | *lux*_new*Bsa*IOH_fwd | GATCGGTCTCAGTGAAATTCAAAAAAAAATC |
| OLEC10026 | $P_{23}$_new*Bsa*IOH_fwd | GATCGGTCTCATCAGAATTCAAAAAAAAATC |
| OLEC10027 | $P_{23}$_new*Bsa*IOH_rev | GATCGGTCTCTTCACTGCAGTACTAG |
| OLEC10182 | *ffluc*_new*Bsa*IOH_rev | GATCGGTCTCTCACACTGCAGTACTAGTA |
| OLEC10311 | MCS+CmR_fwd | GATCGATCCGTCTCCCAGCGAACGAATATTGGATAAATATG |
| OLEC10312 | MCS+CmR_rev | GATCGATCCGTCTCGACATATCTGGAGCTGTAATATAAAAAC |
| OLEC10315 | pBR322ori_fwd | GATCGATCCGTCTCTCTAACTGTCAGACCAAGTTTACTCATAT |

| Oligo Code | Name | 5'-3'-Sequence |
|---|---|---|
| OLEC10316 | pBR322ori_rev | GATCGATCCGTCTCTAGGTCTTTCCAGTCGGGAAACCTGTC |
| OLEC10930 | *amyA*_up_fwd | GATCGATCCGTCTCGACCTGAATCAAGATTATTCTAGACGC |
| OLEC10931 | *amyA*_up_rev | GATCGATCCGTCTCTGCTGTTAGTTACTAAACACAATATATCC |
| OLEC10932 | *amyA*_down_fwd | GATCGATCCGTCTCTATGTATGGTAGAGAACCAAAGACAACAG |
| OLEC10933 | *amyA*_down_rev | GATCGATCCGTCTCGTTAGCTGTCGCACCATCAACGGTTG |
| OLEC11104 | pSpy1C+tt0_OH_fwd | CCGGCGGCAACCGAGCGTTCTGAATTATGGGGATGATGTTAAGGC |
| OLEC11105 | pSpy1C+tt0_OH_rev | CGCCGGGCGTTTTTTATGTTATCCAATATTCGTTCCTTA |
| OLEC11463 | pSpy0C3MCS_CmR_f | CAACTTTTTGGATTTGATTATTTGGAACGAATATTGGATAAATATGGG |
| OLEC11464 | pSpy0C3MCS_CmR_r | CCTTAATTCTAACATAGTTTTATTCCATCTGGAGCTGTAATATAAAAACCT |
| OLEC11465 | pSpy0C3*mazEF*_dw_f | TTTTTATATTACAGCTCCAGATGGAATAAAAACTATGTTAGAATTAAGG |
| OLEC11466 | pSpy0C3*mazEF*_dw_r | GAGTAAACTTGGTCTGACAGACTGGAATTTGTTTTAGGGGT |
| OLEC11467 | pSpy0C3pBR322ori_f | CCCTAAAACAAATTCCAGTCTGTCAGACCAAGTTTACTC |
| OLEC11468 | pSpy0C3pBR322ori_r | AATGTTGAAGCTATCGTTCCTTTCCAGTCGGGAAAC |
| OLEC11469 | pSpy0C3*mazEF*_up_f | GTTTCCCGACTGGAAAGGAACGATAGCTTCAACATTTAATCAG |
| OLEC11470 | pSpy0C3*mazEF*_up_r | CCCATATTTATCCAATATTCGTTCCAAATAATCAAATCCAAAAAAGTTGAC |
| OLEC11471 | pSpy0C4MCS_CmR_f | AGGTACTAGATAGTACCTGCGAACGATAAAAAACGCCCG |
| OLEC11472 | pSpy0C4MCS_CmR_r | GATATTTCATTGAGACTCCTTAGATCTGGAGCTGTAATATAAAAACC |
| OLEC11473 | pSpy0C4F_dw_f | GTTTTTATATTACAGCTCCAGATCTAAGGAGTCTCAATGAAATATC |
| OLEC11474 | pSpy0C4_dw_r | GCGTTTTTTATGTCTGACAGGGTTACTCGTCAAGGAG |
| OLEC11475 | pSpy0C4pBR322ori_f | AACTCCTTGACGAGTAACCCTGTCAGACATAAAAAACGCC |
| OLEC11476 | pSpy0C4pBR322ori_r | CAAGGAGGAAGTCCACTGCTTTCCAGTCGGGAAACCTG |
| OLEC11477 | pSpy0C4_up_f | GGTTTCCCGACTGGAAAGCAGTGGACTTCCTCCTTGG |
| OLEC11478 | pSpy0C4_up_r | CGGGCGTTTTTTATCGTTCGCAGGTACTATCTAGTACCTGC |
| OLEC11479 | pSpy0C4_dw_r | GAGTAAACTTGGTCTGACAGGGTTACTCGTCAAGGAG |
| OLEC11480 | pSpy0C4pBR322ori_f | AACTCCTTGACGAGTAACCCTGTCAGACCAAGTTTACTC |
| OLEC11535 | pSpy0C4_*sagB*_up_r | CGGGCGTTTTTTATCGTTCCGCGAGTCGTTTATTTTTAACC |
| OLEC12102 | *mKate2*_GibsOH_F | TCGCATGCTCCTCTAGACTCGAGCCAGTGAATTCATGGTTTCAGAACT |
| OLEC12103 | *mKate2*_GibsOH_R | CGCGTTGGGAGCTCTCCGGATCCCATGATTACGCCAAGCTTGCA |
| OLEC12104 | *mNeon*_GibsOH_F | TCGCATGCTCCTCTAGACTCGAGCAGTGAATTCATGGTTTCAAAAGGTG |

| Oligo Code | Name | 5'-3'-Sequence |
|---|---|---|
| OLEC12105 | p7INT_*mKate2*_OH_F | TGCAAGCTTGGCGTAATCATGGGATCCGGAGAGCTCCCAACGC |
| OLEC12106 | p7INT_*mKate2*_OH_R | AGTTCTGAAACCATGAATTCACTGGCTCGAGTCTAGAGGAGCATGCGA |
| OLEC12107 | p7INT_*mNeon*_OH_R | CACCTTTTGAAACCATGAATTCACTGCTCGAGTCTAGAGGAGCATGCGA |
| OLEC12108 | *mKate2*_F | CTTACCAGTGAATTCATGGTTTCAG |
| OLEC12109 | *mNeongreen*_F | CTTACAGTGAATTCATGGTTTCAAAAG |
| OLEC12110 | *mKate2_mNeon*_R | CTTACATGATTACGCCAAGCTTGC |
| OLEC12196 | pSpy0C4_*mKate2*_F | GTTTCAGAACTTATCAAAGAAAAC |
| OLEC12197 | pSpy0C4_R | GAATTCAGCTGGTCTCGCT |
| OLEC12198 | pSpy0C4_*mNg*_F | GTTTCAAAAGGTGAAGAAGAC |
| OLEC12228 | p7INT_*mNg*_F | GTTTCAAAAGGTGAAGAAGACAA |
| OLEC12229 | p7INT_*mNg_mK2*_R | CCTATAGTGAGTCGTATTACAAT |
| OLEC12230 | p7INT_*mK2*_F | GTTTCAGAACTTATCAAAGAAAA |
| OLEC12261 | $P_{23}$-5'UTR_0C4OH_F | ATATGGGGATGATGTTAAGGCTACGAATTGGAGCTCTCGAAAAGC |
| OLEC12262 | $P_{23}$-5'UTR_*mNg*_OH_R | TCTTCTTCACCTTTTGAAACCATTTTTATTCCTCCTTAGATCGCCG |
| OLEC12263 | $P_{23}$-5'UTR_*mK2*_OH_R | TCTTTGATAAGTTCTGAAACCATTTTTATTCCTCCTTAGATCGCCG |
| OLEC12264 | $P_{23}$-5'UTR_p7INT_OH_F | ACGACTCACTATAGGGCGAATTGCGAATTGGAGCTCTCGAAAAGC |
| OLEC12265 | *mNg*_$P_{23}$Gibs_F | ATGGTTTCAAAAGGTGAAGAAG |
| OLEC12266 | *mKate2*_$P_{23}$Gibs_F | ATGGTTTCAGAACTTATCAAAG |
| OLEC12267 | p7INT_Gibs_R | CAATTCGCCCTATAGTGAGTC |
| OLEC12268 | pSpy0C4_Gibs_R | TAGCCTTAACATCATCCCCA |
| OLEC13040 | Fmuta *ssrA* LAA to ASV | CAAACTCTTACGCTGCCTCTGTTTAAGATCGATCTGG |
| OLEC13041 | Rmuta *ssrA* LAA to ASV | CCAGATCGATCTTAAACAGAGGCAGCGTAAGAGTTTG |
| OLEC13042 | Fmuta *ssrA* LAA to LDD | CAAACTCTTACGCTTTAGATGATTAAGATCGATCTGG |
| OLEC13043 | Rmuta *ssrA* LAA to LDD | CCAGATCGATCTTAATCATCTAAAGCGTAAGAGTTTG |
| OLEC13044 | Fmuta STOP *mNeon* | CGAACTTTACAAATAAAATACAAACTC |
| OLEC13045 | Rmuta STOP *mNeon* | GAGTTTGTATTTTATTTGTAAAGTTCG |
| OLEC13046 | Fmuta STOP *mKate2* | GGTCACCGTGCATAAAATACAAACTC |
| OLEC13047 | Rmuta STOP *mKate2* | GAGTTTGTATTTTATGCACGGTGACC |
| OLEC13445 | $P_{nisA}$_F | AGTCTTATAACTATACTGACAAT |
| OLEC13446 | $P_{nisA}$_R | TTTGAGTGCCTCCTTATAATT |
| OLEC13447 | *nisRK*_F | ATTGTCAGTATAGTTATAAGACTTATAGTGTGTATCTCAATCC |
| OLEC13448 | *nisRK*_R | GAAGCTAGAGTAAGTAGTTC |

| Oligo Code | Name | 5'-3'-Sequence |
|---|---|---|
| OLEC13449 | 1C_*ffluc*_$P_{nisA}$OH_F | AATTATAAGGAGGCACTCAAAATGGAAGACGCCAAAAACAT |
| OLEC13450 | 1C_*ffluc*_*nisRK*OH_R | GAACTACTTACTCTAGCTTCGACTGGCCGTCGTTTTACC |
| OLEC13456 | $P_{Tre}$_F | AGGGTAAATTACCTTTCCAT |
| OLEC13457 | $P_{Tre}$_R | GTGACATCCCCTAGTCCTTT |
| OLEC13458 | 1C_*ffluc*_$P_{Tre}$OH_F | AGGACTAGGGGATGTCACATGGAAGACGCCAAAAACAT |
| OLEC13459 | 1C_*ffluc*_$P_{Tre}$OH_R | ATGGAAAGGTAATTTACCCTGACTGGCCGTCGTTTTACC |
| OLEC13460 | Pzn_F | ATCGGACACTTTCTAAATG |
| OLEC13461 | Pzn_R | AATTTTCTCCTTTTGTTATAC |
| OLEC13462 | 1C_*ffluc*_$P_{Zn}$OH_F | AGTATAACAAAAGGAGAAAATTATGGAAGACGCCAAAAAC AT |
| OLEC13463 | 1C_*ffluc*_$P_{Zn}$OH_R | AACATTTAGAAAGTGTCCGATGACTGGCCGTCGTTTTACC |
| OLEC13464 | *lacI*_F | TTATTGTCCACTTTCCAAACG |
| OLEC13465 | *lacI*_$P_{lacI}$OH_R | CTCACAATTACTAGTGAATTCGACTGTACGTCATCAGAAGTT |
| OLEC13466 | $P_{lacI}$_F | GAATTCACTAGTAATTGTGAG |
| OLEC13467 | $P_{lacI}$_R | AGATCCTTTCTCCTCTTTAGA |
| OLEC13468 | 1C_*ffluc*_$P_{lacI}$OH_F | TAAAGAGGAGAAAGGATCTATGGAAGACGCCAAAAACAT |
| OLEC13469 | 1C_*ffluc*_$P_{lacI}$OH_R | TTTGGAAAGTGGACAATAAGACTGGCCGTCGTTTTACC |
| OLEC13494 | *ffluc*_0C4GibsOH_F | GTAAAACGACGGCCAGTCGAATTCAAAAAAAAATCTAGAAG |
| OLEC13495 | *ffluc*_0C4GibsOH_R | CAGGAAACAGCTATGACCACTGCAGTACTAGTATTAC |
| OLEC13496 | pSpy0C4_Gibs_F | GTCATAGCTGTTTCCTGC |
| OLEC13497 | pSpy0C4_Gibs_R | GACTGGCCGTCGTTTTAC |
| OLEC14306 | MCS_F | CGAATTATTAACGCTTACGGCTCACCTTCGGGTGGGCCTTTC TGCGTTTATACCTAGGGATATAGTAAAACGACGGCCAG |
| OLEC14307 | MCS_R | GGAAGAGCGCCCAATACCAGGAAACAGCTATGAC |
| OLEC14308 | p7INT_F | GTATTGGGCGCTCTTCC |
| OLEC14309 | p7INT_R | GTAAGCGTTAATAATTCGAGC |
| OLEC14310 | Mut_*int*_F | GAACTTGTTGAAAAAACTGACAACATTGAAAAAATAG |
| OLEC14311 | Mut_*int*_R | CAACAAGTTCAAGGAATTGTTTAAGTTGATCAAGTG |
| OLEC14312 | $P_{veg}$_F | GCGGGTTTTTTATTTTAC |
| OLEC14313 | $P_{veg}$_R | GTTTCATTTTTTATTCCTCCTTTAACTACATTTATTGTACAAC |
| OLEC14314 | *lacI*_F | AAGGAGGAATAAAAAATGAAACCAGTCACCCTTTAC |
| OLEC14315 | *lacI*_R | CTATAGAATACTCAAGCTATGAAAAAAGGCCCACTTTTGTG GGCCTTTTTTGATCGATCGATCTTATTGTCCACTTTCCAAACG |
| OLEC14316 | p7INT(*lacI*)_F | CATAGCTTGAGTATTCTATAGTG |
| OLEC14317 | p7INT($P_{veg}$)_R | ATGTAAAATAAAAAACCCGCCCCTATAGTGAGTCGTATTAC |
| OLEC14322 | 0C5_Frag3_F | CTGTCAGACCAAGTTTACTC |
| OLEC14323 | 0C5_Frag3_R | CTTTCCAGTCGGGAAACC |
| OLEC14326 | 0C5_LFH_R | CTATATCCCTAGGTATAAACG |

| Oligo Code | Name | 5'-3'-Sequence |
|---|---|---|
| OLEC14327 | 0K6_MCS_F | GTTTGGATATAGTAAAACGACGGCCAGTC |
| OLEC14328 | 0K6_MCS_R | CAAGTTTTGGGATTGTTAAGCAGGAAACAGCTATGAC |
| OLEC14329 | 0K6_kanR_F | CTTAACAATCCCAAAACTTG |
| OLEC14330 | 0K6_kanR_R | AAGTCAATGGTTAGATTCACCCTATCTAGCGAACTTTTAG |
| OLEC14331 | 0K6_HRup_F | TGAAAAAGACTGAGGGTC |
| OLEC14332 | 0K6_HRup_R | GTCGTTTTACTATATCCAAACGCAGAAAGGCCCACCCGAAG GTGAGCCCATAATTCCATATCAATTTTTA |
| OLEC14333 | 0K6_ori_F | CTGTCAGACCAAGTTTACTC |
| OLEC14334 | 0K6_ori_R | GACCCTCAGTCTTTTTCACTTTCCAGTCGGGAAACC |
| OLEC14335 | 0K6_HRdown_F | GTGAATCTAACCATTGACTT |
| OLEC14336 | 0K6_HRdown_R | GTAAACTTGGTCTGACAGGTCAAGTTCATCTTGGCAT |
| OLEC14337 | 0K6_LFH_R | GTCGTTTTACTATATCCAAAC |
| OLEC14357 | 1C-tt0_*ffluc*_F | ATGGAAGACGCCAAAAAC |
| OLEC14358 | 1C-tt0_*ffluc*_R | GACTGGCCGTCGTTTTAC |
| OLEC14359 | $P_{Mn}$_F | GTAAAACGACGGCCAGTCCATCTCACCTACTTCCGTA |
| OLEC14360 | $P_{Mn}$_R | GTTTTTGGCGTCTTCCATAATAACTCCTTTGATTTCAAC |
| OLEC14363 | $P_{Cu}$_F | GTAAAACGACGGCCAGTCCATAAGGTCAGTAATAGACC |
| OLEC14364 | $P_{Cu}$_R | GTTTTTGGCGTCTTCCATAACGTCACCTTCATTCAC |
| OLEC14365 | pSpy2C_bb_F | CATATTCACCAATAGCCTTAAGTAAAACGACGGCCAGTC |
| OLEC14366 | pSpy2C_bb_R | TTATAAAAGCCAGTCATTAGGC |
| OLEC14367 | pSpy2C_pAMb1_F | CTAATGACTGGCTTTTATAAATGTCTGGTCCCTAGCGCTTAG AA |
| OLEC14368 | pSpy2C_pAMb1_R | TTAAGGCTATTGGTGAATATGGTACCGAGCTCGAATTCAG |
| OLEC14369 | pSpy2E_bb_R | AAGCGACTCATAGAATTAT |
| OLEC14370 | pSpy2E_pAMb1_F | ATAATTCTATGAGTCGCTTTGGTCCCTAGCGCTTAGAA |
| OLEC14377 | $P_{ermB}$_F | GCCTATCATTTTCAATGAAAG |
| OLEC14380 | 1C-tt0-*ffluc*2_R | CTTTCATTGAAAATGATAGGCGACTGGCCGTCGTTTTAC |
| OLEC14398 | Ori_1_F | CTGTCAGACCAAGTTTACTCATATATAC |
| OLEC14399 | Ori_1_R | GTTAAAATACAATTACTATAACCAAAAATGAGCACCTGTTC AATAGTGAACAACATTTAGAAAGTGTCCGATGAGAGGCGGT TTGCGTATTG |
| OLEC14402 | Ery_3_F | ATCCCCCACAAAAAGAAAAAC |
| OLEC14403 | Ery_3_R | CTGGCCGTCGTTTTACGTTTTAGATTTACCTAATAATTTATCT AC |
| OLEC14404 | MRFP1_4_F | GTAAAACGACGGCCAGTCAGC |
| OLEC14405 | MRFP1_4_R | GAGTAAACTTGGTCTGACAGCAGGAAACAGCTATGACACAT GAG |
| OLEC14522 | ErmB_3_F_new | AAACGAAATGATACACCAATCAG |
| OLEC14523 | Ori_1_R_new | aaagagaggcggtttgcgtattg |

| Oligo Code | Name | 5'-3'-Sequence |
|---|---|---|
| OLEC14524 | $P_{veg}$_frag_2_F | caatacgcaaaccgcctctc |
| OLEC14525 | Pterm_frag_2_R | ATTGGTGTATCATTTCG |
| OLEC14502 | OC5_F1_R_new | CCCATTTAAGAGCGCATAGATCTGGAGCTGTAATATAAAAAC |
| OLEC14503 | OC5_F2_F_new | CTATGCGCTCTTAAATGGG |
| OLEC14504 | OC5_F2_R_new | GTAAACTTGGTCTGACAGGTAGCATGACTGTATGTATCAAG |
| OLEC14505 | OC5_F4_F_new | GGTTTCCCGACTGGAAAGATGAGTTCGGGCTGTTTAGTAG |
| OLEC14506 | OC5_F4_R_new | CTATATCCCTAGGTATAAACGCAGAAAGGCCCACCCGAAGGTGAGCCGTGCGTAGTGCTTGTAAGG |
| OLEC14548 | $P_{lac}$_mut_F | GTAAAACGACGGCCAGTCTTATAGAATTCACTAGTAATTG |
| OLEC14549 | $P_{lac}$_mut1_R | GTTTTTGGCGTCTTCCATATGTTCCTCCTTAGATCTTTTGAATTCG |
| OLEC14550 | $P_{lac}$_mut2_R | GTTTTTGGCGTCTTCCATATGTTTCCTCCTTAGATCTTTTGAATTCG |
| OLEC14551 | $P_{lac}$_mut3_R | GTTTTTGGCGTCTTCCATATGTTCCTCTTTAGATCTTTTGAATTCG |
| OLEC14582 | *ermB*_UTR_new_R | GTTTTGAGAATATTTTATATTTTTG |
| OLEC14583 | 1C-*ffluc*_*ermB*OH_F | CAAAAATATAAAATATTCTCAAAACGAAGACGCCAAAAACATAAAG |
| OLEC14584 | $P_{xylS2}$_F | GAATTAGATATTTAAAAGTATCAT |
| OLEC14585 | $P_{xylS2}$_R | TGATTTAAGTGAACAAGTTTATC |
| OLEC14586 | 1C-tt0-*ffluc*_$P_{xylS2}$_F | AAACTTGTTCACTTAAATCAAGAATTAAATTAAAAAGGGAGGCC |
| OLEC14587 | 1C-tt0-*ffluc*_$P_{xylS2}$_R | ATGATACTTTTAAATATCTAATTCCTGACTGGCCGTCGTTTTA |
| OLEC14589 | $P_{veg}$_mut1_F | CTTATTAACATGGATATAATTTAAATTTTATTTGAC |
| OLEC14590 | $P_{veg}$_mut1_R | GTTAATAAGAAAATGTCAATAAAATTATTTTGAC |
| OLEC14591 | $P_{veg}$_mut2_F | GACAAAACGTCTTATTAACGTTGATATAATTTAAA |
| OLEC14592 | $P_{veg}$_mut2_R | GACGTTTTGTCAATAAAATTATTTTGACAAAATTC |
| OLEC14593 | $P_{veg}$_mut3_F | CAACGTCATTAACGTTGATATAATTTAAATTTTATTTG |
| OLEC14594 | $P_{veg}$_mut3_R | GTTAATGACGTTGTCAATAAAATTATTTTGAC |
| OLEC14717 | $P_{23}$_*mKate2*_F | CGAGACCAGCTGAATTCCTCGAAAAGCCCTGACAACCC |
| OLEC14718 | *mKate2*_R | GACACATGAGACCTTCAGATCCGAGAAAAAAGGCCCAC |
| OLEC14719 | 0K6_Gibs_F | TGAAGGTCTCATGTGTCATAG |
| OLEC14720 | 0K6_Gibs_R | GAATTCAGCTGGTCTCGCTG |
| OLEC14721 | *mKate2*_F | CGAGACCAGCTGAATTCATGGTTTCAGAACTTATCAAAGAAAAC |
| OLEC14726 | RBS_F | AAGGAGGAATAAAAAATGGACCTTCAAGCTCAATTG |
| OLEC14727 | RBS_R | TTTTTATTCCTCCTTTAACTACATTTATTGTACAAC |
| OLEC14875 | *sagA*_up_F | GATCCTTCCTCCTTGGTTTTAAG |
| OLEC14876 | *sagA*_up_R | GATCCAACTATCTAGTTCTTATCAC |

| Oligo Code | Name | 5'-3'-Sequence |
|---|---|---|
| OLEC14877 | *sagA*_do_F | GATCCTATTTAGCATCTCTATGTG |
| OLEC14878 | *sagA*_do_R | GATCCAATTTTCAGCATTAATAGC |
| OLEC14879 | *sagA*_up_OH_F | CTGAATTCAAAAAAAATCTAGATGCTTCCTCCTTGGTTTTAAG |
| OLEC14880 | *sagA*_up_OH_R | CAACTATCTAGTTCTTATCAC |
| OLEC14881 | *sagA*_do_OH_F | GTGATAAGAACTAGATAGTTGCTATTTAGCATCTCTATGTG |
| OLEC14882 | *sagA*_do_OH_R | CTATGACACATGAGACCTTCCAATTTTCAGCATTAATAGC |
| OLEC14883 | pERASE_F | GAAGGTCTCATGTGTCATAG |
| OLEC14884 | pERASE_R | CATCTAGATTTTTTTTGAATTCAG |

**Supplementary Table 3.** Plasmids used in this study. Information on all plasmids used either for cloning purposes or for *in vivo* experiments including the plasmid names ('designation'), their encoded resistance genes ('marker') and a short description of their most important characteristics ('features'). Symbols: (~) indicates translational fusions; TA = Toxin-Antitoxin; (*) indicates stop codon (TAA), except for *pheS**, where (*) highlights the introduction of two mutations.

| Code | Designation | Marker | Reference | Features |
|---|---|---|---|---|
| **Plasmids used for cloning purposes** | | | | |
| pEC536 | pEU8517 | Kan$^R$, Cm$^R$ | Bugrysheva & Scott 2010 (3) | $P_{tet}$(O)3 cassette, *cat86*, Ω-Km2 element, pUC ori |
| pRDN18 | pMSP3535 | Erm$^R$ | Bryan *et al.* 2000 (4) | *ermB*, *nisRK* operon, P*nisA* promoter, pAMβ1 replicon, pUC ori |
| pEC2173 | pLZ12Km2-P*23*R:TA:*ffluc* | Kan$^R$ | Loh & Proft 2013 (5) | *aph(3')-IIIa*, ω-ε-ζ (TA cassette of pBT286), pSH71 replicon, *ffluc* under control of $P_{23}$ |
| pEC2192 | pBAV1K-T5-*gfp* | Kan$^R$ | Bryksin *et al.* 2010 (6) | *aph(3')-IIIa*, $P_{T5}$_*gfp*, modified pWV01 replicon |
| pEC2466 | pIB166 | Cm$^R$ | Biswas *et al.* 2008 (7) | *cat*, pSH71 replicon, $P_{23}$ promoter |
| pEC2467 | pIB167 | Cm$^R$ | Biswas *et al.* 2008 (7) | *cat*, pSH71 replicon, $P_{spac}$ promoter, coding regions for N-terminal His-tag and C-terminal Strep-tag |
| pEC2468 | pIB184-Km | Kan$^R$ | Biswas *et al.* 2008 (7) | *aph(3')-IIIa*, pAMβ1 replicon, $P_{23}$ promoter, pUC ori |
| pEC2469 | pIB185 | Erm$^R$ | Biswas *et al.* 2008 (7) | *ermB*, pAMβ1 replicon, $P_{veg}$ promoter, pUC ori |
| pEC2471 | pBS3C*lux* | Cm$^R$ | Radeck *et al.* 2013 (8) | *bla*, *cat*, *luxABCDE* operon, pUC ori, MCS with $P_{lac(Eco)}$_*mrfp*, integrates into *sacA* in *B. subtilis* |
| pEC3003 | pJWV102-$P_{lac}$_*dcas9sp* | Amp$^R$, Tet$^R$ | Liu *et al.* 2017 (9) | *dcas9* under control of IPTG inducible promoter ($P_{lac}$), *bla*, integrates into *bgaA* in *S. pneumoniae*, *tetM* |
| pEC3004 | pPEPY-PF6_*lacI* | Kan$^R$, Gm$^R$ | Liu *et al.* 2017 (9) | *lacI-aacC1* under control of $P_{F6}$ promoter, *aph(3')-IIIa*, integrates into *prsA* in *S. pneumoniae* |
| pEC3045 | pLZ12Km2-TA:$P_{gyrA(Sag)}$-*ermBL-ermB'*:A2I:*ffluc*-tt3 | Kan$^R$ | Wulff *et al.* 2023 (10) | *aph(3')-IIIa*, ω-ε-ζ (TA cassette of pBT286), *ffluc* under control of erythromycin-inducible riboswitch $P_{gyrA(Sag)}$-*ermBL-ermB'*, pSH71 replicon |
| **Replicative backbones** | | | | |
| pEC2483 | pSpy1C | Cm$^R$ | This study | pSH71 replicon, *cat*, $P_{lac(Eco)}$_*mrfp* in MCS |
| pEC2485 | pSpy1K | Kan$^R$ | This study | pSH71 replicon, *aph(3')-IIIa*, $P_{lac(Eco)}$_*mrfp* in MCS |
| pEC2484 | pSpy1E | Erm$^R$ | This study | pSH71 replicon, *ermB*, $P_{lac(Eco)}$_*mrfp* in MCS |
| pEC3112 | pSpy2C | Cm$^R$ | This study | pAMβ1 replicon, *cat*, $P_{lac(Eco)}$_*mrfp* in MCS, pUC ori |

| Code | Designation | Marker | Reference | Features |
|---|---|---|---|---|
| pEC3113 | pSpy2E | $Erm^R$ | This study | pAMβ1 replicon, *ermB*, $P_{lac(Eco)}$_*mrfp* in MCS, pUC ori |
| pEC2498 | pSpy3C | $Cm^R$ | This study | modified pWV01 replicon, *cat*, $P_{lac(Eco)}$_*mrfp* in MCS |
| pEC2488 | pSpy3K | $Kan^R$ | This study | modified pWV01 replicon, *aph(3')-IIIa*, $P_{lac(Eco)}$_*mrfp* in MCS |
| pEC2506 | pSpy3E | $Erm^R$ | This study | modified pWV01 replicon, *ermB*, $P_{lac(Eco)}$_*mrfp* in MCS |
| | | **Integrative backbones** | | |
| pEC2735 | pSpy0C2 | $Cm^R$ | This study | pUC ori, *cat*, flanking regions for allelic replacement of *amyA* locus, MCS with $P_{lac(Eco)}$_*mrfp* |
| pEC2794 | pSpy0C3 | $Cm^R$ | This study | pUC ori, *cat*, flanking regions for allelic replacement of *SPy_0938-0939*, MCS with $P_{lac(Eco)}$_*mrfp* |
| pEC2796 | pSpy0C4 | $Cm^R$ | This study | pUC ori, *cat*, flanking regions for allelic replacement of *sagB*, MCS with $P_{lac(Eco)}$_*mrfp* |
| pEC3077 | pSpy0C5 V1 | $Cm^R$ | This study | pUC ori, *cat*, flanking regions for allelic replacement of *SPy_1930*, MCS with $P_{lac(Eco)}$_*mrfp* |
| pEC3138 | pSpy0C5 V1 | $Cm^R$ | This study | pUC ori, *cat*, adapted flanking regions for allelic replacement of *SPy_1930*, MCS with $P_{lac(Eco)}$_*mrfp* |
| pEC3078 | pSpy0K6 | $Kan^R$ | This study | pUC ori, *aph(3')-IIIa*, flanking regions for allelic replacement of *SPy_1078*, MCS with $P_{lac(Eco)}$_*mrfp* |
| pEC2825 | p7INT | $Erm^R$ | McShan et al. 1998 (11) | pUC ori, *ermB*, *int* and *att*P from T12 bacteriophage, MCS with $P_{lacZ}$_*lacZα* |
| pEC3103 | p7INT.1 | $Erm^R$ | This study | pUC ori, *ermB*, *int* and *att*P from T12 bacteriophage, MCS with $P_{lac(Eco)}$_*mrfp* |
| | | **Gene deletion plasmids** | | |
| pEC3135 | pERASE | $Erm^R$ | This study | pUC ori, MCS with $P_{lac(Eco)}$_*mrfp, pheS** under control of $P_{veg}$, *ermB* |
| pEC3136 | pERASE-*ΔsagA* | $Erm^R$ | This study | pUC ori, *pheS** under control of P*veg*, *ermB*, *sagA* flanking regions |
| | | **Reporter plasmids** | | |
| pEC2858 | pSpy1C-$P_{23}$_*ffluc* | $Cm^R$ | This study | *ffluc* reporter under control of $P_{23}$, *cat*, pSH71 replicon |
| pEC2860 | pSpy1C-$P_{veg}$_*ffluc* | $Cm^R$ | This study | *ffluc* reporter under control of $P_{veg}$, *cat*, pSH71 replicon |
| pEC2861 | pSpy1C-$P_{gyrA}$_*ffluc* | $Cm^R$ | This study | *ffluc* reporter under control of $P_{gyrA}$, *cat*, pSH71 replicon |
| pEC3105 | pSpy1C-$P_{xylS2}$_*ffluc* | $Cm^R$ | This study | *ffluc* reporter under control of $P_{xylS2}$, *cat*, pSH71 replicon |

| Code | Designation | Marker | Reference | Features |
|---|---|---|---|---|
| pEC3106 | pSpy1C-P$_{veg(1)}$_*ffluc* | Cm$^R$ | This study | *ffluc* reporter under control of P$_{veg(1)}$, *cat*, pSH71 replicon |
| pEC3107 | pSpy1C-P$_{veg(2)}$_ *ffluc* | Cm$^R$ | This study | *ffluc* reporter under control of P$_{veg(2)}$, *cat*, pSH71 replicon |
| pEC3108 | pSpy1C-P$_{veg(3)}$_ *ffluc* | Cm$^R$ | This study | *ffluc* reporter under control of P$_{veg(3)}$, *cat*, pSH71 replicon |
| pEC2862 | pSpy1C-*tetR*-P$_{tet}$_*ffluc* | Cm$^R$ | This study | *ffluc* reporter under control of tetracycline-inducible cassette *tetR*-P$_{Tet}$, *cat*, pSH71 replicon |
| pEC2863 | pSpy1C-P$_{gyrA(Spy)}$_*xylR*-P$_{xylS2}$_*ffluc* | Cm$^R$ | This study | *ffluc* reporter under control of xylose-inducible cassette P$_{gyrA(Spy)}$_*xylR*-P$_{xylS2}$, *cat*, pSH71 replicon |
| pEC3005 | pSpy1C-P$_{nisRK}$_*nisRK*-P$_{nisA}$_*ffluc* | Cm$^R$ | This study | *ffluc* reporter under control of nisin-inducible cassette P$_{nisRK}$_*nisRK*-P$_{nisA}$, *cat*, pSH71 replicon |
| pEC3006 | pSpy1C-P$_{Zn}$_*ffluc* | Cm$^R$ | This study | *ffluc* reporter under control of zinc-inducible promoter P$_{czcD}$, *cat*, pSH71 replicon |
| pEC3007 | pSpy1C-P$_{tre}$_*ffluc* | Cm$^R$ | This study | *ffluc* reporter under control of zinc-inducible promoter P$_{SPy_2097}$, *cat*, pSH71 replicon |
| pEC3020 | pSpy1C-PF6_*lacI*-P$_{lac(Spn)}$_*ffluc* | Cm$^R$ | This study | *ffluc* reporter under control of IPTG-inducible promoter P$_{lac(Spn)}$, *lacI* under control of P$_{F6}$, *cat*, pSH71 replicon |
| pEC3081 | pSpy1C-P$_{Cu}$_*ffluc* | Cm$^R$ | This study | *ffluc* reporter under control of copper-inducible promoter P$_{copA}$, *cat*, pSH71 replicon |
| pEC3080 | pSpy1C-P$_{Mn}$_*ffluc* | Cm$^R$ | This study | *ffluc* reporter under control of manganese-inducible promoter P$_{mntE}$, *cat*, pSH71 replicon |
| pEC3109 | pSpy1C-P$_{lac(1)}$_*ffluc* | Cm$^R$ | This study | *ffluc* reporter under control of IPTG-inducible promoter P$_{lac(1)}$, *cat*, pSH71 replicon |
| pEC3110 | pSpy1C-P$_{lac(2)}$_*ffluc* | Cm$^R$ | This study | *ffluc* reporter under control of IPTG-inducible promoter P$_{lac(2)}$, *cat*, pSH71 replicon |
| pEC3111 | pSpy1C-P$_{lac(3)}$_*ffluc* | Cm$^R$ | This study | *ffluc* reporter under control of IPTG-inducible promoter P$_{lac(t3)}$, *cat*, pSH71 replicon |
| pEC3104 | pSpy1C- P$_{gyrA(Sag)}$-*ermBL_ermB'~ffluc* | Erm$^R$ | This study | *ffluc* reporter under control of erythromycin-inducible riboswitch P$_{gyrA(Sag)}$-*ermBL_ermB'*, *cat*, pSH71 replicon |
| pEC2826 | pUC19-*mKate2~ssrA* | Amp$^R$ | This study | *bla, mKate2* fused to native *ssrA* tag (codon optimized for *S. pyogenes*), pUC ori |
| pEC2827 | pUC19-*mNeongreen~ssrA* | Amp$^R$ | This study | *bla, mKate2* fused to native *ssrA* tag (codon optimized for *S. pyogenes*), pUC ori |

| Code | Designation | Marker | Reference | Features |
|---|---|---|---|---|
| pEC2828 | p7INT-*mKate2~ssrA* | $Erm^R$ | This study | pUC ori, *ermB*, *int* and *att*P from T12 bacteriophage, *mKate2* fused to native *ssrA* tag |
| pEC3137 | p7INT-*mKate2~*ssrA* | $Erm^R$ | This study | pUC ori, *ermB*, *int* and *att*P from T12 bacteriophage, *mKate2* fused to native *ssrA* tag with a preceding stop codon |
| pEC2829 | p7INT-*mNeongreen~ssrA* | $Erm^R$ | This study | pUC ori, *ermB*, *int* and *att*P from T12 bacteriophage, *mNeongreen* fused to native *ssrA* tag |
| pEC2968 | p7INT-*mNeongreen~*ssrA* | $Erm^R$ | This study | pUC ori, *ermB*, *int* and *att*P from T12 bacteriophage, *mNeongreen* fused to native *ssrA* tag with a preceding stop codon |
| pEC2931 | p7INT-$P_{23}$_*mNeongreen ~ssrA* | $Erm^R$ | This study | pUC ori, *ermB*, *int* and *att*P from T12 bacteriophage, *mNeongreen* fused to native *ssrA* tag under control of $P_{23}$ |
| pEC2934 | p7INT-$P_{23}$_*mKate2~*ssrA* | $Erm^R$ | This study | pUC ori, *ermB*, *int* and *att*P from T12 bacteriophage, *mKate2* fused to native *ssrA* tag with a preceding stop codon under control of $P_{23}$ |
| pEC2936 | p7INT-$P_{23}$_*mNeongreen~* ssrA* | $Erm^R$ | This study | pUC ori, *ermB*, *int* and *att*P from T12 bacteriophage, *mNeongreen* fused to native *ssrA* tag with a preceding stop codon under control of $P_{23}$ |
| pEC2935 | p7INT-$P_{23}$_*mNeongreen~ ssrA*(LDDmut) | $Erm^R$ | This study | pUC ori, *ermB*, *int* and *att*P from T12 bacteriophage, *mNeongreen* fused to native *ssrA* tag with introduced LDD mutation under control of $P_{23}$ |
| pEC2938 | p7INT-$P_{23}$_*mNeongreen~ ssrA*(ASVmut) | $Erm^R$ | This study | pUC ori, *ermB*, *int* and *att*P from T12 bacteriophage, *mNeongreen* fused to native *ssrA* tag with introduced ASV mutation under control of $P_{23}$ |
| pEC3115 | pSpy0K6-$P_{23}$_*mKate2~*ssrA* | $Kan^R$ | This study | pUC ori, *aph(3')-IIIa*, flanking regions for allelic replacement of *SPy_1078*, *mKate2* fused to native *ssrA* tag with a preceding stop codon under control of $P_{23}$ |
| pEC3116 | pSpy0K6-*mKate2~*ssrA* | KanR | This study | pUC ori, *aph(3')-IIIa*, flanking regions for allelic replacement of *SPy_1078*, *mKate2* fused to native *ssrA* tag with a preceding stop codon |
| pEC2939 | pSpy0C4-$P_{23}$_*mKate2~*ssrA* | $Cm^R$ | This study | pUC ori, *cat*, flanking regions for allelic replacement of *sagB*, *mKate2* fused to native *ssrA* tag with a preceding stop codon under control of $P_{23}$ |
| pEC3114 | pSpy0C4-*mKate2~*ssrA* | $Cm^R$ | This study | pUC ori, *cat*, flanking regions for allelic replacement of *sagB*, *mKate2* fused to native *ssrA* tag with a preceding stop codon |

**Supplementary Table 4.** Information on the DNA sequence of the reporters, promoters and other genetic parts used in this study. Modifications to the original sequences are highlighted in red. The start codons (ATG) as well as the sequence part of the erythromycin-inducible riboswitch are underlined. The -10 and -35 motifs of the promoter sequences were predicted using BPROM and are highlighted in italics (12). The truncated *ermB'* coding sequence that is translationally fused to the downstream reporter is marked in yellow. For the constitutive promoters, we added the optimized 5'UTR of pLZ12Km2-P*23*R:TA:*ffluc* that is colored in grey. Putative binding sites of regulatory proteins are highlighted in bold. The *ssrA* tag sequence attached to the fluorescent reporter genes is marked in blue. $P_{gyrA(Sag)}$ – $P_{gyrA}$ promoter from *Streptococcus agalactiae*.

| Designation | 5'-3' DNA sequence |
|---|---|
| | **Reporter genes** |
| *ffluc* | ATGGAAGACGCCAAAAACATAAAGAAAGGCCCGGCGCCATTCTATCCTCTAGAGGATGGAACCGCTGGAGAGCAACTGCATA<br>AGGCTATGAAGAGATACGCCCTGGTTCCTGGAACAATTGCTTTTACAGATGCACATATCGAGGTGAACATCACGTACGCGGAA<br>TACTTCGAAATGTCCGTTCGGTTGGCAGAAGCTATGAAACGATATGGGCTGAATACAAATCACAGAATCGTCGTATGCAGTGA<br>AAACTCTCTTCAATTCTTTATGCCGGTGTTGGGCGCGTTATTTATCGGAGTTGCAGTTGCGCCCGCGAACGACATTTATAATGA<br>ACGTGAATTGCTCAACAGTATGAACATTTCGCAGCCTACCGTAGTGTTTGTTTCCAAAAAGGGGTTGCAAAAAATTTTGAACG<br>TGCAAAAAAAATTACCAATAATCCAGAAAATTATTATCATGGATTCTAAAACGGATTACCAGGGATTTCAGTCGATGTACACG<br>TTCGTCACATCTCATCTACCTCCCGGTTTTAATGAATACGATTTTGTACCAGAGTCCTTTGATCGTGACAAAACAATTGCACTG<br>ATAATGAATTCCTCTGGATCTACTGGGTTACCTAAGGGTGTGGCCCTTCCGCATAGAACTGCCTGCGTCAGATTCTCGCATGCC<br>AGAGATCCTATTTTTGGCAATCAAATCATTCCGGATACTGCGATTTTAAGTGTTGTTCCATTCCATCACGGTTTTGGAATGTTTA<br>CTACACTCGGATATTTGATATGTGGATTTCGAGTCGTCTTAATGTATAGATTTGAAGAAGAGCTGTTTTTACGATCCCTTCAGG<br>ATTACAAAATTCAAAGTGCGTTGCTAGTACCAACCCTATTTTCATTCTTCGCCAAAAGCACTCTGATTGACAAATACGATTTAT<br>CTAATTTACACGAAATTGCTTCTGGGGGCGCACCTCTTTCGAAAGAAGTCGGGGAAGCGGTTGCAAAACGCTTCCATCTTCCA<br>GGGATACGACAAGGATATGGGCTCACTGAGACTACATCAGCTATTCTGATTACACCCGAGGGGGATGATAAACCGGGCGCGG<br>TCGGTAAAGTTGTTCCATTTTTTGAAGCGAAGGTTGTGGATCTGGATACCGGGAAAACGCTGGGCGTTAATCAGAGAGGCGAA<br>TTATGTGTCAGAGGACCTATGATTATGTCCGGTTATGTAAACAATCCGGAAGCGACCAACGCCTTGATTGACAAGGATGGATG<br>GCTACATTCTGGAGACATAGCTTACTGGGACGAAGACGAACACTTCTTCATAGTTGACCGCTTGAAGTCTTTAATTAAATACA<br>AAGGATATCAGGTGGCCCCCGCTGAATTGGAATCGATATTGTTACAACACCCCAACATCTTCGACGCGGGCGTGGCAGGTCTT<br>CCCGACGATGACGCCGGTGAACTTCCCGCCGCCGTTGTTGTTTTGGAGCACGGAAAGACGATGACGGAAAAAGAGATCGTGG<br>ATTACGTCGCCAGTCAAGTAACAACCGCGAAAAAGTTGCGCGGAGGAGTTGTGTTTGTGGACGAAGTACCGAAAGGTCTTACC<br>GGAAAACTCGACGCAAGAAAAATCAGAGAGATCCTCATAAAGGCCAAGAAGGGCGGAAAGTCCAAATTGTAA |
| *mKate2~ssrA*<br>(codon-optimized for *S. pyogenes*) | ATGGTTTCAGAACTTATCAAAGAAAACATGCACATGAAACTTTACATGGAAGGTACTGTTAACAACCACCACTTCAAATGTAC<br>TTCAGAAGGTGAAGGTAAACCATACGAAGGTACTCAAACTATGCGTATCAAAGCTGTTGAAGGTGGTCCACTTCCATTCGCTT<br>TCGACATCCTTGCTACTTCATTCATGTACGGTTCAAAAACTTTCATCAACCACACTCAAGGTATCCCAGACTTCTTCAAACAAT<br>CATTCCCAGAAGGTTTCACTTGGGAACGTGTTACTACTTACGAAGACGGTGGTGTTCTTACTGCTACTCAAGACACTTCACTTC<br>AAGACGGTTGTCTTATCTACAACGTTAAAATCCGTGGTGTTAACTTCCCATCAAACGGTCCAGTTATGCAAAAAAAAACTCTTG<br>GTTGGGAAGCTTCAACTGAAACTCTTTACCCAGCTGACGGTGGTCTTGAAGGTCGTGCTGACATGGCTCTTAAACTTGTTGGTG<br>GTGGTCACCTTATCTGTAACCTTAAAACTACTTACCGTTCAAAAAAACCAGCTAAAAACCTTAAAATGCCAGGTGTTTACTACG<br>TTGACCGTCGTCTTGAACGTATCAAAGAAGCTGACAAAGAAACTTACGTTGAACAACACGAAGTTGCTGTTGCTCGTTACTGT<br>GACCTTCCATCAAAACTTGGTCACCGTGCAAAAAATACAAACTCTTACGCTTTAGCTGCCTAA |

| Designation | 5'-3' DNA sequence |
|---|---|
| *mNeongreen~ssrA* (codon-optimized for *S. pyogenes*) | ATGGTTTCAAAAGGTGAAGAAGACAACATGGCTTCACTTCCAGCTACTCACGAACTTCACATCTTCGGTTCAATCAACGGTGTT<br>GACTTCGACATGGTTGGTCAAGGTACTGGTAACCCAAACGACGGTTACGAAGAACTTAACCTTAAATCAACTAAAGGTGACCT<br>TCAATTCTCACCATGGATCCTTGTTCCACACATCGGTTACGGTTTCCACCAATACCTTCCATACCCAGACGGTATGTCACCATTC<br>CAAGCTGCTATGGTTGACGGTTCAGGTTACCAAGTTCACCGTACTATGCAATTCGAAGACGGTGCTTCACTTACTGTTAACTAC<br>CGTTACACTTACGAAGGTTCACACATCAAAGGTGAAGCTCAAGTTAAAGGTACTGGTTTCCCAGCTGACGGTCCAGTTATGAC<br>TAACTCACTTACTGCTGCTGACTGGTGTCGTTCAAAAAAAACTTACCCAAACGACAAAACTATCATCTCAACTTTCAAATGGTC<br>ATACACTACTGGTAACGGTAAACGTTACCGTTCAACTGCTCGTACTACTTACACTTTCGCTAAACCAATGGCTGCTAACTACCT<br>TAAAAACCAACCAATGTACGTTTTCCGTAAAACTGAACTTAAACACTCAAAAACTGAACTTAACTTCAAAGAATGGCAAAAAG<br>CTTTCACTGACGTTATGGGTATGGACGAACTTTACAAAGCAAAAAATACAAACTCTTACGCTTTAGCTGCCTAA |
| | **Inducible promoters** |
| $P_{czcD}$ ($P_{Zn}$) | ATCGGACACTTTCTAAATGTT**TGTTCACTA*TTGAACA***GGTGCTCATTTTT*GGTTATAGT*AATTGTATTTTAACTTTTAGATAATAG<br>AATAAAGCATTCTACTATACATATTACCAGTATAACAAAAGGAGAAAATT |
| $P_{MntE}$ ($P_{Mn}$) | CATCTCACCTACTTCCGTATTGTCTTTATCTTAATGGTAGGCCTAGGAGGCTTCCTTAAGTTAGAACTAATTTGGGTTTTGGCTG<br>ATATTGTCAATGGTTTAATGGCACTTCCTAACCTGATTGCCCTCCTAGCTCTCTCGCCAGTTGTTATTTTAGAAACCAAGCATTA<br>CTTTATTAAGTAAACATGACGTTAGTACTACCCAGTTATACCTAGTCTTCTTCTGCCAAGTTCAAGACGCTAGGCTT*TTTTCT*TA<br>AGCCCCTTTTTA*TGGTATAAT*ATAAAGTTGAAATCAAAGGAGTTATT |
| $P_{copA}$ ($P_{Cu}$) | CATAAGGTCAGTAATAGACCTAAGGTGACAAGTGTTTTTCGAATAGGATGTTTCAAGGGTATCTCCTTAATGTAAATGTTATTT<br>TTATTGCAATAATATTATAGTAAAAACGATTCATTTGTCAATATAAGGATAGAAAAATGAGAAAAGTTATTTA*TTTAC*AAATGT<br>AGTTAAATT*CGTTATAAT*TAATTTACAAACGTAAATATATTGTGAATGAAGGTGACGTT |
| $P_{SPy\_2097}$ ($P_{tre}$) | AGGGTAAATTACCTTTCCATTTGTCTATT*TTTTTA*CAGACAACACC*TTTTATTTT*ACCAAAATAATGGCTTTTTTGGCACAA**ACAA**<br>**CTTGTAGACAAATTTG**CAATCGATTACAATAAAGGTGTCAGTTAGATTCTATCTAAGAATTTGAATATTTCATTTTGACGTTG<br>CTTGTTAAAAGCAACTAGAACAAAGGACTAGGGGATGTCAC |
| $P_{lac(Spn)}$ | GAATTCACTAGTAATTGTGAGGCGGATAACAATTCTCACTCATTCTACAGTTTATTC*TTGACA*TTGCACTGTCCCCCTGG*TATAAT*<br>AACT**AATTGTGAGCGCTCACAATT**AAGATCCCCGGGGCGGCCGCGAATTCAAAAGATCTAAAGAGGAGAAAGGATCT |
| $P_{lac(1)}$ | GAATTCACTAGTAATTGTGAGGCGGATAACAATTCTCACTCATTCTACAGTTTATTC*TTGACA*TTGCACTGTCCCCCTGG*TATAAT*<br>AACTAATTGTGAGCGCTCACAATTAAGATCCCCGGGGCGGCCGCGAATTCAAAAGATCTAAGGAGGAACAT |
| $P_{lac(2)}$ | GAATTCACTAGTAATTGTGAGGCGGATAACAATTCTCACTCATTCTACAGTTTATTC*TTGACA*TTGCACTGTCCCCCTGG*TATAAT*<br>AACTAATTGTGAGCGCTCACAATTAAGATCCCCGGGGCGGCCGCGAATTCAAAAGATCTAAGGAGGAAACAT |
| $P_{lac(3)}$ | GAATTCACTAGTAATTGTGAGGCGGATAACAATTCTCACTCATTCTACAGTTTATTC*TTGACA*TTGCACTGTCCCCCTGG*TATAAT*<br>AACTAATTGTGAGCGCTCACAATTAAGATCCCCGGGGCGGCCGCGAATTCAAAAGATCTAAAGAGGAACAT |
| $P_{gyrA(Sag)}$_*ermBL*_ *ermB'* | GCCTATCATTTTCAATGAAAGAAGTCACTAATAAAATGTGAAAAAATTTGCAAACAAATTGAAATATCATAGAAAACTATTGA<br>AAACGCTGAGTGTC*TTGATA*TTTTTGATACCG*ATTTATCAT*CGCAATTTTACCTAAATTATGGTACAATGTAAGAGGAAGTTAAA<br>TTAGATGCTAAAAATTTGTAATTAAGAAGGAGGGATTCGTCATGTTGGTATTCCAAATGCGTAATGTAGATAAAACATCTACT<br>GTTTTGAAACAGACTAAAAACAGTGATTACGCAGATAAATAAATACGTTAGATTAATTCCTACCAGTGACTAATCTTATGACT<br>TTTTAAACAGATAACTAAAATTACAAACAAATCGTTTAACTTCTGTATTTATTTATAGATGTAATCACTTCAGGAGTGATTACA<br>TGAACAAAAATATAAAATATTCTCAAAAC |
| | **Constitutive promoters** |
| $P_{gyrA(Spy)}$_5'UTR | TTTTTGTGTTGTTTAGTTGCAGTCATTTCTAAACATCTCCTTAATTTTATTAGAGCAAAAGCTCATACGGTCTTATTTTAACATA*T*<br>*TGATA*ATTGTTTGTCATTGTATTCAG*AAAAAT*AATGAAAATAGTTTACTGATAACGCTTCCAATAAGAAAAAATAAGGAATTTAT |

| Designation | 5'-3' DNA sequence |
|---|---|
| | GGTATAATGAAATCTAGTATTGACTAGAATACTAGTACTGCAGTGAAATTCAAAAAAATGGCCGGCAGAATTAAATTAAAAA<br>GGGAGGCCAAATATAATGAAAAATATGAATGACAATGATGTTCCGCGGTGGCGGCGATCTAAGGAGGAATAAAAA |
| $P_{23}$_5'UTR | CTCGAAAAGCCCTGACAACCCTTGTTCCTAAAAAGGAATAAGCGTTCGGTCAGTAAATAATAGAAATAAAAAATCAGACCTA<br>AGACTG*ATGACA*AAAAGAGAAAATTTTGA*TAAAAT*AGTCTTAGAATTAAATTAAAAAGGGAGGCCAAATATAATGAAAAATAT<br>GAATGACAATGATGTTCCGCGGTGGCGGCGATCTAACCGGTTAATACTAGAATTAAATTAAAAAGGGAGGCCAAATATAATG<br>AAAAATATGAATGACAATGATGTTCCGCGGTGGCGGCGATCTAAGGAGGAATAAAAA |
| $P_{xylS2}$_5'UTR | AGCGAGACCAGCTGAATTCAATTAGATATTTAAAAGTATCATATCTAATATTATAACTAAATTTTCTAAAAAAAACA*TTGAAA*T<br>AAACATTTATTTTGTA*TATGAT*GAGATAAAGTTAGTTTATTGGATAAACAAACTAACTCAATTAAGATAGTTGATGGATAAACT<br>TGTTCACTTAAATCAGAATTAAATTAAAAAGGGAGGCCAAATATAATGAAAAATATGAATGACAATGATGTTCCGCGGTGGC<br>GGCGATCTAAGGAGGAATAAAAA |
| $P_{veg}$_5'UTR | GGGAGTTCTGAGAATTGGTATGCCTTATAAGTCCAATTAACAGTTGAAAACCTGCATAGGAGAGCTATGCGGGTTTTTTATTTT<br>ACATAATGATACATAATTTACCGAAACTTGCGGAACATAATTGAGGAATCATAGAATTTTGTCAAAATAATTTTA*TTGACA*ACG<br>TCTTATTAACGTTGA*TATAAT*TTAAATTTTATTTGACAAAAATGGGCTCGTGTTGTACAATAAATGTAGTTACTAGTACTGCAGT<br>GAAATTCAAAAAAATGGCCGGCAGAATTAAATTAAAAAGGGAGGCCAAATATAATGAAAAATATGAATGACAATGATGTTCC<br>GCGGTGGCGGCGATCTAAGGAGGAATAAAAA |
| $P_{veg(1)}$_5'UTR | GGGAGTTCTGAGAATTGGTATGCCTTATAAGTCCAATTAACAGTTGAAAACCTGCATAGGAGAGCTATGCGGGTTTTTTATTTT<br>ACATAATGATACATAATTTACCGAAACTTGCGGAACATAATTGAGGAATCATAGAATTTTGTCAAAATAATTTTA*TTGACA*ACG<br>TCTTATTAACATGGA*TATAAT*TTAAATTTTATTTGACAAAAATGGGCTCGTGTTGTACAATAAATGTAGTTACTAGTACTGCAGT<br>GAAATTCAAAAAAATGGCCGGCAGAATTAAATTAAAAAGGGAGGCCAAATATAATGAAAAATATGAATGACAATGATGTTCC<br>GCGGTGGCGGCGATCTAAGGAGGAATAAAAA |
| $P_{veg(2)}$_5'UTR | GGGAGTTCTGAGAATTGGTATGCCTTATAAGTCCAATTAACAGTTGAAAACCTGCATAGGAGAGCTATGCGGGTTTTTTATTTT<br>ACATAATGATACATAATTTACCGAAACTTGCGGAACATAATTGAGGAATCATAGAATTTTGTCAAAATAATTTTA*TTGACA*AAA<br>CGTCTTATTAACGTTGA*TATAAT*TTAAATTTTATTTGACAAAAATGGGCTCGTGTTGTACAATAAATGTAGTTACTAGTACTGCA<br>GTGAAATTCAAAAAAATGGCCGGCAGAATTAAATTAAAAAGGGAGGCCAAATATAATGAAAAATATGAATGACAATGATGTT<br>CCGCGGTGGCGGCGATCTAAGGAGGAATAAAAA |
| $P_{veg(3)}$_5'UTR | GGGAGTTCTGAGAATTGGTATGCCTTATAAGTCCAATTAACAGTTGAAAACCTGCATAGGAGAGCTATGCGGGTTTTTTATTTT<br>ACATAATGATACATAATTTACCGAAACTTGCGGAACATAATTGAGGAATCATAGAATTTTGTCAAAATAATTTT*ATTGACA*ACG<br>TC**ATTAACGTTGA*TATAAT*TTAAATTTTATTTGACAAAAATGGGCTCGTGTTGTACAATAAATGTAGTTACTAGTACTGCAGT<br>GAAATTCAAAAAAATGGCCGGCAGAATTAAATTAAAAAGGGAGGCCAAATATAATGAAAAATATGAATGACAATGATGTTCC<br>GCGGTGGCGGCGATCTAAGGAGGAATAAAAA |
| **Other** | |
| $P_{f6}$_*lacI* | GACTGTACGTCATCAGAAGTTTCAGCGACCCTAGATCCTGCTCGAAGATTTCAGC*TTGACA*TTGCACTGTCCCCCTGG*TATAAT*A<br>ACTATACATGCAAGGATCTAAATAAATAAGGAGGAAAATTAATGAAACCAGTCACCCTTTACGATGTCGCAGAATACGCAGG<br>AGTCTCTTACCAAACAGTCAGTCGTGTAGTCAATCAGGCTAGTCATGTATCAGCAAAAACTCGTGAAAAGGTTGAAGCTGCAA<br>TGGCTGAACTTAATTACATCCCAAATCGTGTTGCCCAACAGTTGGCTGGTAAACAAAGTTTGCTTATCGGAGTTGCTACCTCTT<br>CATTGGCACTTCATGCCCCTTCTCAAATTGTTGCCGCTATCAAATCTCGTGCAGATCAATTGGGTGCAAGTGTTGTCGTATCTAT<br>GGTTGAACGTAGTGGAGTCGAAGCCTGTAAGGCAGCCGTCCACAATTTATTGGCTCAACGTGTATCTGGTTTAATTATCAATTA<br>CCCATTGGATGACCAAGATGCCATTGCTGTCGAAGCTGCATGTACCAATGTACCAGCATTATTTTTGGATGTTTCAGATCAAAC<br>ACCTATCAATTCAATCATCTTTAGTCATGAAGATGGTACTCGTTTGGGAGTCGAACATTTGGTAGCTCTTGGTCACCAACAGAT<br>TGCTCTTTTAGCAGGACCACTTAGTTCTGTTTCAGCCCGTCTTCGTTTAGCTGGTTGGCACAAATACTTAACTCGTAATCAAATT |

| Designation | 5'-3' DNA sequence |
|---|---|
| | CAGCCTATCGCTGAACGTGAGGGTGATTGGTCTGCAATGTCAGGATTTCAACAGACAATGCAAATGTTGAATGAAGGTATTGT<br>ACCAACTGCAATGTTGGTTGCCAATGACCAAATGGCCCTTGGAGCTATGCGTGCAATTACCGAATCAGGTTTGCGTGTTGGAG<br>CTGATATTAGTGTTGTCGGTTACGATGATACAGAAGATTCAAGTTGTTACATTCCACCTTTGACAACTATCAAGCAAGATTTTC<br>GTTTGCTTGGACAGACAAGTGTTGATCGTTTATTGCAACTTTCTCAGGGTCAGGCTGTCAAGGGAAATCAGCTTTTACCTGTAT<br>CTCTTGTTAAGCGTAAGACCACATTAGCACCAAATACTCAAACCGCCTCACCTCGTGCCTTGGCAGATAGTCTTATGCAGCTTG<br>CACGTCAGGTCTCTCGTTTGGAAAGTGGACAATAA |
| $P_{veg}$_*lacI* | GCGGGTTTTTATTTTACATAATGATACATAATTTACCGAAACTTGCGGAACATAATTGAGGAATCATAGAATTTTGTCAAAAT<br>AATTTTA*TTGACA*ACGTCTTATTAACGTTGA*TATAAT*TTAAATTTTATTTGACAAAAATGGGCTCGTGTTGTACAATAAATGTAGT<br>TAAAGGAGGAATAAAAA<u>ATG</u>AAACCAGTCACCCTTTACGATGTCGCAGAATACGCAGGAGTCTCTTACCAAACAGTCAGTCG<br>TGTAGTCAATCAGGCTAGTCATGTATCAGCAAAAACTCGTGAAAAGGTTGAAGCTGCAATGGCTGAACTTAATTACATCCCAA<br>ATCGTGTTGCCCAACAGTTGGCTGGTAAACAAAGTTTGCTTATCGGAGTTGCTACCTCTTCATTGGCACTTCATGCCCCTTCTCA<br>AATTGTTGCCGCTATCAAATCTCGTGCAGATCAATTGGGTGCAAGTGTTGTCGTATCTATGGTTGAACGTAGTGGAGTCGAAGC<br>CTGTAAGGCAGCCGTCCACAATTTATTGGCTCAACGTGTATCTGGTTTAATTATCAATTACCCATTGGATGACCAAGATGCCAT<br>TGCTGTCGAAGCTGCATGTACCAATGTACCAGCATTATTTTTGGATGTTTCAGATCAAACACCTATCAATTCAATCATCTTTAG<br>TCATGAAGATGGTACTCGTTTGGGAGTCGAACATTTGGTAGCTCTTGGTCACCAACAGATTGCTCTTTTAGCAGGACCACTTAG<br>TTCTGTTTCAGCCCGTCTTCGTTTAGCTGGTTGGCACAAATACTTAACTCGTAATCAAATTCAGCCTATCGCTGAACGTGAGGG<br>TGATTGGTCTGCAATGTCAGGATTTCAACAGACAATGCAAATGTTGAATGAAGGTATTGTACCAACTGCAATGTTGGTTGCCA<br>ATGACCAAATGGCCCTTGGAGCTATGCGTGCAATTACCGAATCAGGTTTGCGTGTTGGAGCTGATATTAGTGTTGTCGGTTACG<br>ATGATACAGAAGATTCAAGTTGTTACATTCCACCTTTGACAACTATCAAGCAAGATTTTCGTTTGCTTGGACAGACAAGTGTTG<br>ATCGTTTATTGCAACTTTCTCAGGGTCAGGCTGTCAAGGGAAATCAGCTTTTACCTGTATCTCTTGTTAAGCGTAAGACCACAT<br>TAGCACCAAATACTCAAACCGCCTCACCTCGTGCCTTGGCAGATAGTCTTATGCAGCTTGCACGTCAGGTCTCTCGTTTGGAAA<br>GTGGACAATAA |
| *pheS** | <u>ATG</u>GACCTTCAAGCTCAATTGGAAGAACTTAAAACTAAAACCTTGGAAACTCTTCAAAGTCTTACTGGTAACCATACAAAAGA<br>ATTGCAAGATTTGCGTGTTGCAGTTCTTGGTAAGAAGGGTTCACTTACTGATTTGCTTAAAGGTCTTAAGGATTTGTCAAACGA<br>TTTGCGTCCTGTGGTAGGCAAACAAGTAAACGAAGTTCGTGACTTATTGACAAAAGCTTTCGAAGAGCAAGCTAAGATCGTTG<br>AAGCCGCTAAAATTCAAGCTCAATTGGATGCTGAATCAATCGACGTTACTCTTCCTGGACGTCAAATGACTCTGGGGCATCGT<br>CACGTTCTTACTCAAACATCAGAAGAAATCGAAGACATCTTCCTTGGTATGGGTTTCCAAATTGTAGATGGTTTCGAAGTTGAA<br>AAAGATTATTATAACTTCGAACGTATGAACCTTCCAAAAGACCACCCTGCTCGTGACATGCAAGACACTTTCTACATTACTGA<br>AGAGATCTTGTTACGTACTCACACTTCACCAGTACAAGCTCGTACTCTTGATCAACACGACTTCTCTAAGGGCCCTTTAAAAAT<br>GGTTTCACCAGGCCGTGTATTCCGCCGCGACACTGACGACGCTACTCACTCACACCAATTCCACCAAATCGAAGGTCTTGTAGT<br>TGGCAAAAATATTTCAATGGGTGACTTGAAAGGCACTTTGGAAATGATCATTAAAAAAATGTTCGGTGACGAACGTAGCATCC<br>GTCTTCGCCCTTCTTATTTCCCTTTCTCAGAACCATCAGTAGAAGTTGACGTATCATGTTTCAAATGTGGAGGTAAAGGCTGTA<br>ACGTTTGTAAAAAAACTGGTTGGATCGAAATCTTGGGTGCTGGTATGGTTCACCCAAGTGTTCTTGAAATGTCAGGTGTTGATG<br>CAAAAGAGTACTCAGGTTTCGGTTTCGGCTTGGGACAAGAGCGTATCGCTATGTTGCGCTACGGAATTAACGATATTCGTGGC<br>TTCTACCAAGGCGACCAACGTTTCTCAGAACAATTTAACTAA |

**Supplementary Table 5.** Details of the construction of reporter plasmids featuring different constitutive promoters upstream of the firefly luciferase reporter *ffluc*. The first column ('Plasmid') indicates the name of the plasmid that was constructed. The second column ('Assembly method') defines which assembly strategy was used. The amplicons or fragments required for plasmid assembly were obtained by PCR and the amplicons, respective oligonucleotides and the templates used are specified in the following columns. Abbreviations: TA = Toxin-Antitoxin

| Plasmid | Assembly Method | Amplicon | Oligo Code | Template |
|---|---|---|---|---|
| pSpy1C-$P_{23}$_*ffluc* | Golden Gate Assembly | $P_{23}$ promoter | OLEC10026 | pIB166 |
| | | | OLEC10027 | |
| | | *ffluc* reporter (+ Cloning overhangs) | OLEC10182 | pLZ12Km2-P*23*R:TA:*ffluc* |
| | | | OLEC10024 | |
| pSpy1C-$P_{veg}$_*ffluc* | Golden Gate Assembly | $P_{veg}$ promoter | OLEC9749 | *Bacillus subtilis* genomic DNA |
| | | | OLEC9750 | |
| | | $P_{veg}$ promoter (+ *Eco*31I recognition sites) | OLEC10026 | $P_{veg}$ amplicon |
| | | | OLEC10027 | |
| | | *ffluc* reporter | OLEC9562 | pLZ12Km2-P*23*R:TA:*ffluc* |
| | | | OLEC9563 | |
| | | *ffluc* reporter (+ Cloning overhangs) | OLEC10182 | *ffluc* amplicon |
| | | | OLEC10024 | |
| pSpy1C-$P_{gyrA}$_*ffluc* | Golden Gate Assembly | $P_{gyrA}$ promoter | OLEC9591 | *Streptococcus pyogenes* SF370 genomic DNA |
| | | | OLEC9592 | |
| | | $P_{gyrA}$ promoter (+ *Eco*31I recognition sites) | OLEC10026 | $P_{gyrA}$ amplicon |
| | | | OLEC10027 | |
| | | *ffluc* reporter | OLEC9562 | pLZ12Km2-P*23*R:TA:*ffluc* |
| | | | OLEC9563 | |
| | | *ffluc* reporter (+ Cloning overhangs) | OLEC10182 | *ffluc* amplicon |
| | | | OLEC10024 | |
| pSpy1C-$P_{xylS2}$_*ffluc* | Gibson Assembly | $P_{xylS2}$ promoter | OLEC14584 | gene synthesis fragment (GenScript services) |
| | | | OLEC14585 | |
| | | pSpy1C-*ffluc* | OLEC14586 | pSpy1C-*ffluc* |
| | | | OLEC14587 | |
| pSpy1C-$P_{veg(1)}$_*ffluc* | Gibson Assembly | PCR mutagenesis | OLEC14589 | pSpy1C-$P_{veg}$_*ffluc* |
| | | | OLEC14590 | |
| pSpy1C-$P_{veg(2)}$_*ffluc* | Gibson Assembly | PCR mutagenesis | OLEC14591 | pSpy1C-$P_{veg}$_*ffluc* |
| | | | OLEC14592 | |
| pSpy1C-$P_{veg(3)}$_*ffluc* | Gibson Assembly | PCR mutagenesis | OLEC14593 | pSpy1C-$P_{veg}$_*ffluc* |
| | | | OLEC14594 | |

**Supplementary Table 6.** Details of the construction of reporter plasmids featuring different inducible promoters upstream of the firefly luciferase reporter *ffluc*. The first column ('Plasmid') indicates the name of the plasmid that was constructed. The second column ('Assembly method') defines which assembly strategy was used to ligate the fragments together to create the plasmid. The amplicons or fragments required for plasmid assembly were obtained by PCR and the amplicons, respective oligonucleotides and the templates used are specified in following columns. Symbols: (~) indicates translational fusions; TA = Toxin-Antitoxin

| Plasmid | Assembly Method | Amplicon | Oligo Name | Template |
|---|---|---|---|---|
| pSpy1C-*nisRK*-$P_{nisA}$_*ffluc* | Gibson Assembly | $P_{nisRK}$_*nisRK* | OLEC13447 | pMSP3535 |
| | | | OLEC13448 | |
| | | $P_{nisA}$ promoter | OLEC13445 | pMSP3535 |
| | | | OLEC13446 | |
| | | pSpy1C-*ffluc* | OLEC13449 | pSpy1C-$P_{23}$_*ffluc* |
| | | | OLEC13450 | |
| pSpy1C-*tetR*-$P_{tet}$_*ffluc* | Golden Gate Assembly | *tetR*-$P_{tet}$ | OLEC9595 | pEU8517 |
| | | | OLEC9596 | |
| | | *tetR*-$P_{tet}$ (+*Eco*31I recognition sites) | OLEC10026 | *tetR*-$P_{tet}$ amplicon |
| | | | OLEC10027 | |
| | | *ffluc* reporter | OLEC10024 | pLZ12Km2-P*23*R:TA:*ffluc* |
| | | | OLEC10182 | |
| pSpy1C-$P_{tre}$_*ffluc* | Gibson Assembly | $P_{SPy\_2097}$ ($P_{tre}$) | OLEC13456 | *Streptococcus pyogenes* SF370 genomic DNA |
| | | | OLEC13457 | |
| | | pSpy1C-*ffluc* | OLEC13458 | pSpy1C-$P_{23}$_ffluc |
| | | | OLEC13459 | |
| pSpy1C-$P_{Zn}$_*ffluc* | Gibson Assembly | $P_{czcD}$ ($P_{Zn}$) | OLEC13460 | *Streptococcus pyogenes* SF370 genomic DNA |
| | | | OLEC13461 | |
| | | pSpy1C-*ffluc* | OLEC13462 | pSpy1C-$P_{23}$_*ffluc* |
| | | | OLEC13463 | |
| pSpy1C-$P_{Cu}$_*ffluc* | Gibson Assembly | $P_{copA}$ ($P_{Cu}$) | OLEC14363 | *Streptococcus pyogenes* SF370 genomic DNA |
| | | | OLEC14364 | |
| | | pSpy1C-*ffluc* | OLEC14357 | pSpy1C-$P_{Zn}$_*ffluc* |
| | | | OLEC14358 | |
| pSpy1C-$P_{Mn}$_*ffluc* | Gibson Assembly | $P_{mntE}$ ($P_{Mn}$) | OLEC14359 | *Streptococcus pyogenes* SF370 genomic DNA |
| | | | OLEC14360 | |
| | | pSpy1C-*ffluc* | OLEC14357 | pSpy1C-$P_{Zn}$_*ffluc* |
| | | | OLEC14358 | |
| pSpy1C-$P_{F6}$_*lacI*-$P_{lac(Spn)}$_*ffluc* | Gibson Assembly | $P_{F6}$_*lacI* | OLEC13464 | pPEPY-$P_{F6}$_*lacI* |
| | | | OLEC13465 | |
| | | $P_{lac(Spn)}$ promoter | OLEC13466 | pJW102-PL-dCas9 |

| Plasmid | Assembly Method | Amplicon | Oligo Name | Template |
|---|---|---|---|---|
| | | | OLEC13467 | |
| | | pSpy1C-*ffluc* | OLEC13468 | pSpy1C-$P_{23}$_*ffluc* |
| | | | OLEC13469 | |
| p7INT-$P_{veg}$_*lacI* | Gibson Assembly | $P_{veg}$ promoter | OLEC14312 | pSpy1C-$P_{veg}$_*ffluc* |
| | | | OLEC14313 | |
| | | *lacI* repressor | OLEC14314 | pSpy1C-$P_{F6}$_*lacI*-$P_{lac}$_*ffluc* |
| | | | OLEC14315 | |
| | | p7INT backbone | OLEC14316 | p7INT |
| | | | OLEC14317 | |
| pSpy1C-$P_{lac(1)}$_*ffluc* | Gibson Assembly | $P_{lac(1)}$ | OLEC14548 | pSpy1C-$P_{F6}$_*lacI*-P*lac*_*ffluc* |
| | | | OLEC14549 | |
| | | pSpy1C-*ffluc* | OLEC13457 | pSpy1C-$P_{Zn}$_*ffluc* |
| | | | OLEC13458 | |
| pSpy1C-$P_{lac(2)}$_*ffluc* | Gibson Assembly | $P_{lac(2)}$ | OLEC14548 | pSpy1C-$P_{F6}$_*lacI*-P*lac*_*ffluc* |
| | | | OLEC14550 | |
| | | pSpy1C-*ffluc* | OLEC13457 | pSpy1C-$P_{Zn}$_*ffluc* |
| | | | OLEC13458 | |
| pSpy1C-$P_{lac(3)}$_*ffluc* | Gibson Assembly | $P_{lac(3)}$ | OLEC14548 | pSpy1C-$P_{F6}$_*lacI*-P*lac*_*ffluc* |
| | | | OLEC14551 | |
| | | pSpy1C-*ffluc* | OLEC13457 | pSpy1C-$P_{Zn}$_*ffluc* |
| | | | OLEC13458 | |
| pSpy1C-$P_{gyrA}$_*ermBL-ermB'~ffluc* | Gibson Assembly | $P_{gyrA}$_*ermBL-ermB'* | OLEC14377 | pLZ12-$P_{gyrA}$-*ermBL-ermB'*:A2I:*ffluc* |
| | | | OLEC14582 | |
| | | pSpy1C-*ffluc* | OLEC14583 | pSpy1C-$P_{Zn}$_*ffluc* |
| | | | OLEC14380 | |

**Supplementary Table 7.** Details of the construction of reporter plasmids featuring the fluorescent reporter genes *mKate2* and *mNeongreen*. The first column (‘Plasmid’) indicates the name of the plasmid that was constructed. The second column (‘Assembly method’) defines which assembly strategy was used to ligate the fragments together to create the plasmid. The amplicons or fragments required for plasmid assembly were obtained by PCR and the amplicons, respective oligonucleotides and the templates used are specified in following columns. Symbols: (~) indicates translational fusions; (*) indicates stop codon (TAA)

| Plasmid | Assembly Method | Amplicon | Oligo Name | Template |
|---|---|---|---|---|
| p7INT-*mNeongreen-ssrA* | Gibson Assembly | *mNeongreen-ssrA* | OLEC12109 | pUC19-*mNeongreen-ssrA* |
| | | | OLEC12110 | |
| | | *mNeongreen-ssrA* (+ Cloning overhangs) | OLEC12103 | mNeongreen-ssrA amplicon |
| | | | OLEC12104 | |
| | | p7INT backbone | OLEC12105 | p7INT |
| | | | OLEC12107 | |
| p7INT-*mKate2~ssrA* | Gibson Assembly | *mKate2~ssrA* | OLEC12108 | pUC19-*mKate2~ssrA* |
| | | | OLEC12110 | |
| | | *mKate2~ssrA* (+ Cloning overhangs) | OLEC12102 | *mKate2~ssrA* amplicon |
| | | | OLEC12103 | |
| | | p7INT backbone | OLEC12105 | p7INT |
| | | | OLEC12106 | |
| p7INT-$P_{23}$_*mNeongreen~ssrA* | Gibson Assembly | $P_{23}$ promoter | OLEC12264 | pSpy1C-$P_{23}$_*ffluc* |
| | | | OLEC12262 | |
| | | p7INT-*mNeongreen~ssrA* | OLEC12265 | p7INT-*mNeongreen~ssrA* |
| | | | OLEC12267 | |
| p7INT-$P_{23}$_*mKate2~ssrA* | Gibson Assembly | $P_{23}$ promoter | OLEC12263 | pSpy1C-$P_{23}$_*ffluc* |
| | | | OLEC12264 | |
| | | p7INT-*mKate2~ssrA* | OLEC12266 | p7INT-*mKate2~ssrA* |
| | | | OLEC12267 | |
| pSpy0C4-$P_{23}$_*mNeongreen~ssrA* | Gibson Assembly | $P_{23}$ promoter | OLEC12261 | pSpy1C-$P_{23}$_*ffluc* |
| | | | OLEC12262 | |
| | | pSpy0C4-*mNeongreen~ssrA* | OLEC12265 | pSpy0C4-*mNeongreen~ssrA* |
| | | | OLEC12268 | |
| pSpy0C4-$P_{23}$_*mKate2~ssrA* | Gibson Assembly | $P_{23}$ promoter | OLEC12261 | pSpy1C-$P_{23}$_*ffluc* |
| | | | OLEC12263 | |
| | | pSpy0C4-*mKate2~ssrA* | OLEC12266 | pSpy0C4-*mKate2~ssrA* |
| | | | OLEC12268 | |
| pSpy0K6-*mKate2~*ssrA* | Gibson Assembly | *mKate2~*ssrA* | OLEC14721 | p7INT-*mKate2~*ssrA* |
| | | | OLEC14718 | |
| | | pSpy0K6 backbone | OLEC14719 | pSpy0K6 |

| Plasmid | Assembly Method | Amplicon | Oligo Name | Template |
|---|---|---|---|---|
| | | | OLEC14720 | |
| pSpy0K6-P23_*mKate2~*ssrA* | Gibson Assembly | P23_*mKate2~*ssrA* | OLEC14717 | p7INT-P23_*mKate2~*ssrA* |
| | | | OLEC14718 | |
| | | pSpy0K6 backbone | OLEC14719 | pSpy0K6 |
| | | | OLEC14720 | |

**Supplementary Table 8.** Transcriptionally silent sites in the genome of *S. pyogenes* SF370. The start indicates the start coordinate of the transcriptionally silent region in the genome of the target strain, while the end respectively marks the last coordinate of this region. The different region types are differentiated as follows: (1) not_expressed_genes: regions annotated as genes including the promoter region upstream, but not expressed above threshold and not within annotated prophage region; (2) not_expressed_intergenic: regions not annotated as genes, not expressed above threshold and not within annotated prophage region. For each region, the average normalized read coverage is listed.

| Start | End | Length | Region Type | Average Norm. Coverage |
|---|---|---|---|---|
| 102432 | 104957 | 2526 | not_expressed_genes | 0.016931121523403325 |
| 1800683 | 1803073 | 2391 | not_expressed_genes | 0.015151034098748165 |
| 958923 | 961224 | 2302 | not_expressed_genes | 0.007905311763261175 |
| 1620601 | 1622746 | 2146 | not_expressed_genes | 0.01827871995969429 |
| 1166936 | 1168979 | 2044 | not_expressed_genes | 0.012705785045752685 |
| 710427 | 712411 | 1985 | not_expressed_genes | 0.010902720300538214 |
| 506187 | 508155 | 1969 | not_expressed_genes | 0.018604086811677006 |
| 884387 | 886269 | 1883 | not_expressed_genes | 0.0038432372393893974 |
| 1281009 | 1282630 | 1622 | not_expressed_genes | 0.0113646590107065 |
| 124013 | 125222 | 1210 | not_expressed_genes | 0.01327014526837671 |
| 1595608 | 1596627 | 1020 | not_expressed_genes | 0.015906317658427017 |
| 447282 | 448135 | 854 | not_expressed_genes | 0.009422351236340117 |
| 360546 | 361116 | 571 | not_expressed_genes | 0.015460012804892526 |
| 890337 | 890843 | 507 | not_expressed_genes | 0.013401895442700288 |
| 1803549 | 1804033 | 485 | not_expressed_genes | 0.012248901120239897 |
| 597998 | 598364 | 367 | not_expressed_genes | 0.0076555940892690045 |
| 449773 | 450134 | 362 | not_expressed_genes | 0.006159199007649874 |
| 1709968 | 1710315 | 348 | not_expressed_genes | 0.016581919802625255 |
| 1625956 | 1626263 | 308 | not_expressed_genes | 0.016164889027133717 |
| 1755584 | 1755858 | 275 | not_expressed_genes | 0.01799136215746988 |
| 971336 | 971594 | 259 | not_expressed_genes | 0.01574077903543419 |
| 270237 | 270596 | 360 | not_expressed_intergenic | 0.012492034248109662 |
| 1336434 | 1336658 | 225 | not_expressed_intergenic | 0.011190354406972709 |
| 1803104 | 1803326 | 223 | not_expressed_intergenic | 0.007972241466287083 |
| 390806 | 391013 | 208 | not_expressed_intergenic | 0.015926178855598355 |
| 263746 | 263953 | 208 | not_expressed_intergenic | 0.016675393034894626 |
| 1665879 | 1666082 | 204 | not_expressed_intergenic | 0.013026521027912227 |
| 153837 | 154037 | 201 | not_expressed_intergenic | 0.009320644543472298 |
| 396287 | 396476 | 190 | not_expressed_intergenic | 0.013234815957621913 |
| 263452 | 263638 | 187 | not_expressed_intergenic | 0.01262922746677333 |
| 361133 | 361289 | 157 | not_expressed_intergenic | 0.010517298835780842 |
| 168047 | 168202 | 156 | not_expressed_intergenic | 0.01373345329790649 |
| 597845 | 597997 | 153 | not_expressed_intergenic | 0.01342972821309218 |

| Start | End | Length | Region Type | Average Norm. Coverage |
|---|---|---|---|---|
| 450135 | 450281 | 147 | not_expressed_intergenic | 0.0063777400406086595 |
| 422383 | 422526 | 144 | not_expressed_intergenic | 0.013825369110658483 |
| 1755440 | 1755573 | 134 | not_expressed_intergenic | 0.016500749179115624 |
| 429323 | 429435 | 113 | not_expressed_intergenic | 0.015345124650857998 |
| 662817 | 662924 | 108 | not_expressed_intergenic | 0.016308723798256586 |
| 1493100 | 1493200 | 101 | not_expressed_intergenic | 0.009454247295459012 |

**Supplementary Table 9.** Transcriptionally silent sites in the genome of *S. pyogenes* M1T1 5448. The start indicates the start coordinate of the non-transcribed region in the genome of the target strain, while the end respectively marks the last coordinate of this region. The different region types are differentiated as follows: (1) not_expressed_genes: regions annotated as genes including the promoter region upstream, but not expressed above threshold and not within annotated prophage region; (2) not_expressed_intergenic: regions not annotated as genes, not expressed above threshold and not within annotated prophage region. For each region, the average normalized read coverage is listed.

| Start | End | Length | Region type | Average Norm. Coverage |
|---|---|---|---|---|
| 96786 | 99311 | 2526 | not_expressed_genes | 0.015717676329681903 |
| 1140397 | 1142440 | 2044 | not_expressed_genes | 0.018527613542344997 |
| 820392 | 821774 | 1383 | not_expressed_genes | 0.008018699105305255 |
| 466315 | 467168 | 854 | not_expressed_genes | 0.012071768732113186 |
| 377155 | 377899 | 745 | not_expressed_genes | 0.019622565568269714 |
| 576128 | 576494 | 367 | not_expressed_genes | 0.01062323039617106 |
| 149480 | 149934 | 455 | not_expressed_intergenic | 0.011283169309065798 |
| 289281 | 289621 | 341 | not_expressed_intergenic | 0.011835205232508523 |
| 575950 | 576127 | 178 | not_expressed_intergenic | 0.010403985602063976 |
| 441416 | 441559 | 144 | not_expressed_intergenic | 0.01024269522951294 |
| 1215297 | 1215429 | 133 | not_expressed_intergenic | 0.011895730748578531 |
| 448352 | 448475 | 124 | not_expressed_intergenic | 0.008256789618203633 |
| 1829716 | 1829837 | 122 | not_expressed_intergenic | 0.01747821937253233 |
| 415279 | 415389 | 111 | not_expressed_intergenic | 0.015494377002127614 |

**Supplementary Table 10.** Identification of transcriptionally silent sites showing homology between *S. pyogenes* SF370 and M1T1 5448. The start indicates the start coordinate of the transcriptionally silent region in the genome of the target strain, while the end respectively marks the last coordinate of this region. While the target strain shows the strain in which the transcriptionally silent region was identified, the reference strain is the strain to be compared against. For each region, the average read coverage is listed. Regions that are homologous between reference and target strain are indicated by 'true', while non-homologous sequences are indicated by 'false'. Conserved expression of the regions is marked by 'true', while differences in expression are indicated by 'false'.

| Target Strain | Region Start | Region End | Target_Cov | Ref Strain | Ref Start | Ref End | Alignment Score Norm. | Homolog | Ref Cov | Expression Conserved |
|---|---|---|---|---|---|---|---|---|---|---|
| **SF370** | 102431 | 104956 | 0.01692441878329113 | 5448 | 96786 | 99311 | 100.0 | TRUE | 0.01571145397167332 | TRUE |
| **SF370** | 1800682 | 1803072 | 0.015144697405273156 | 5448 | 1777827 | 1780217 | 100.0 | TRUE | 0.11822973164217583 | FALSE |
| **SF370** | 958922 | 961223 | 0.007901877657369228 | 5448 | 894428 | 896729 | 100.0 | TRUE | 0.02875329957213702 | FALSE |
| **SF370** | 1620600 | 1622745 | 0.01827020238282584 | 5448 | 243742 | 245887 | 64.4 | FALSE | 0.03202424476452032 | FALSE |
| **SF370** | 1166935 | 1168978 | 0.012699568908254764 | 5448 | 1140397 | 1142440 | 100.0 | TRUE | 0.018518549152157948 | TRUE |
| **SF370** | 710426 | 712410 | 0.010897227746230639 | 5448 | 688557 | 690541 | 99.9 | TRUE | 0.029349018400147385 | FALSE |
| **SF370** | 506186 | 508154 | 0.018594638316597436 | 5448 | 525219 | 527187 | 99.9 | TRUE | 0.06169221625124542 | FALSE |
| **SF370** | 884386 | 886268 | 0.0038411962212059722 | 5448 | 819892 | 821774 | 99.9 | TRUE | 0.012549518468624307 | TRUE |
| **SF370** | 1281008 | 1282629 | 0.011357652439183253 | 5448 | 1254343 | 1255964 | 100.0 | TRUE | 0.02955715063034292 | FALSE |
| **SF370** | 124012 | 125221 | 0.013259178206171441 | 5448 | 118367 | 119576 | 100.0 | TRUE | 0.07616098714840958 | FALSE |
| **SF370** | 1595607 | 1596626 | 0.015890723229350128 | 5448 | 269867 | 270886 | 66.7 | FALSE | 0.037698408991480144 | FALSE |
| **SF370** | 447281 | 448134 | 0.0094113180381711 | 5448 | 466315 | 467168 | 100.0 | TRUE | 0.01205763317153694 | TRUE |
| **SF370** | 360545 | 361115 | 0.015432937475987287 | 5448 | 379579 | 380149 | 100.0 | TRUE | 0.020822473022778263 | FALSE |
| **SF370** | 890336 | 890842 | 0.013375461723878395 | 5448 | 825842 | 826348 | 100.0 | TRUE | 0.03791765663422581 | FALSE |
| **SF370** | 1803548 | 1804032 | 0.0122236456540126 | 5448 | 1780693 | 1781177 | 100.0 | TRUE | 0.08617077108755264 | FALSE |
| **SF370** | 597997 | 598363 | 0.00763473415987045 | 5448 | 576128 | 576494 | 100.0 | TRUE | 0.010594284264301384 | TRUE |
| **SF370** | 449772 | 450133 | 0.006142184645750289 | 5448 | 468806 | 469167 | 100.0 | TRUE | 0.018761988756782807 | TRUE |
| **SF370** | 1709967 | 1710314 | 0.016534270607790124 | 5448 | 1700641 | 1700988 | 100.0 | TRUE | 0.04497696285272824 | FALSE |
| **SF370** | 1625955 | 1626262 | 0.01611240562120146 | 5448 | 240225 | 240532 | 63.0 | FALSE | 0.037063036076846925 | FALSE |
| **SF370** | 1755583 | 1755857 | 0.01792593902235181 | 5448 | 1746257 | 1746531 | 100.0 | TRUE | 0.06719254670818217 | FALSE |
| **SF370** | 971335 | 971593 | 0.015680003826803162 | 5448 | 906841 | 907099 | 100.0 | TRUE | 0.03363872223879218 | FALSE |
| **SF370** | 270236 | 270595 | 0.012457334152976025 | 5448 | 289262 | 289621 | 100.0 | TRUE | 0.012332801130031166 | TRUE |

| Target Strain | Region Start | Region End | Target_Cov | Ref Strain | Ref Start | Ref End | Alignment Score Norm. | Homolog | Ref Cov | Expression Conserved |
|---|---|---|---|---|---|---|---|---|---|---|
| **SF370** | 1336433 | 1336657 | 0.011140619498497276 | 5448 | 1308505 | 1308729 | 100.0 | TRUE | 0.03328478649100947 | FALSE |
| **SF370** | 1803103 | 1803325 | 0.007936491504554855 | 5448 | 1780248 | 1780470 | 100.0 | TRUE | 0.027655154604246012 | FALSE |
| **SF370** | 390805 | 391012 | 0.01584961068802336 | 5448 | 409839 | 410046 | 100.0 | TRUE | 0.020757886124931547 | FALSE |
| **SF370** | 263745 | 263952 | 0.016595222876073016 | 5448 | 1597680 | 1597887 | 60.6 | FALSE | 0.04564855355173125 | FALSE |
| **SF370** | 1665878 | 1666081 | 0.012962665532677364 | 5448 | 200406 | 200609 | 66.7 | FALSE | 0.023479065415544193 | FALSE |
| **SF370** | 153836 | 154036 | 0.009274273177584376 | 5448 | 148232 | 148403 | 79.6 | FALSE | 0.026017710789881875 | FALSE |
| **SF370** | 396286 | 396475 | 0.013165159031529166 | 5448 | 415320 | 415509 | 100.0 | TRUE | 0.02083540296653639 | FALSE |
| **SF370** | 263451 | 263637 | 0.012561691491015183 | 5448 | 1597995 | 1598181 | 62.0 | FALSE | 0.04077800987449826 | FALSE |
| **SF370** | 361132 | 361288 | 0.010450309671221729 | 5448 | 380166 | 380322 | 100.0 | TRUE | 0.033433110818197664 | FALSE |
| **SF370** | 168046 | 168201 | 0.013645418340868628 | 5448 | 162457 | 162612 | 99.4 | TRUE | 0.024988144409190127 | FALSE |
| **SF370** | 597844 | 597996 | 0.013341952211699419 | 5448 | 575975 | 576127 | 100.0 | TRUE | 0.009700551619978964 | TRUE |
| **SF370** | 450134 | 450280 | 0.006334354053937852 | 5448 | 469168 | 469314 | 100.0 | TRUE | 0.02209334145262602 | FALSE |
| **SF370** | 422382 | 422525 | 0.013729359602945577 | 5448 | 441416 | 441559 | 100.0 | TRUE | 0.010171565401530212 | TRUE |
| **SF370** | 1755439 | 1755572 | 0.016377609259868493 | 5448 | 1746113 | 1746246 | 100.0 | TRUE | 0.04063210108949035 | FALSE |
| **SF370** | 429322 | 429434 | 0.01520932708757607 | 5448 | 448356 | 448468 | 100.0 | TRUE | 0.00783594089547085 | TRUE |
| **SF370** | 662816 | 662923 | 0.016157717096420877 | 5448 | 640947 | 641054 | 100.0 | TRUE | 0.08769146438524542 | FALSE |
| **SF370** | 1493099 | 1493199 | 0.009360640886593082 | 5448 | 1505973 | 1506073 | 100.0 | TRUE | 0.011315965669465576 | TRUE |
| **5448** | 96785 | 99310 | 0.01571145397167332 | SF370 | 102432 | 104957 | 100.0 | TRUE | 0.01692441878329113 | TRUE |
| **5448** | 1140396 | 1142439 | 0.018518549152157948 | SF370 | 1166936 | 1168979 | 100.0 | TRUE | 0.012699568908254764 | TRUE |
| **5448** | 820391 | 821773 | 0.008012901058229835 | SF370 | 884887 | 886269 | 99.9 | TRUE | 0.003157348477179127 | TRUE |
| **5448** | 466314 | 467167 | 0.01205763317153694 | SF370 | 447282 | 448135 | 100.0 | TRUE | 0.0094113180381711 | TRUE |
| **5448** | 377154 | 377898 | 0.019596226554084117 | SF370 | 358122 | 358866 | 100.0 | TRUE | 0.021847510452131447 | FALSE |
| **5448** | 576127 | 576493 | 0.010594284264301384 | SF370 | 597998 | 598364 | 100.0 | TRUE | 0.00763473415987045 | TRUE |
| **5448** | 149479 | 149933 | 0.01125837113476016 | SF370 | 155240 | 155571 | 71.2 | FALSE | 0.020826826742172695 | FALSE |
| **5448** | 289280 | 289620 | 0.011800497885785624 | SF370 | 270256 | 270596 | 100.0 | TRUE | 0.012104571593766784 | TRUE |
| **5448** | 575949 | 576126 | 0.01034553624474901 | SF370 | 597820 | 597997 | 100.0 | TRUE | 0.01525247166468627 | TRUE |
| **5448** | 441415 | 441558 | 0.010171565401530212 | SF370 | 422383 | 422526 | 100.0 | TRUE | 0.013729359602945577 | TRUE |
| **5448** | 1215296 | 1215428 | 0.011806289164002752 | SF370 | 1241948 | 1242080 | 100.0 | TRUE | 0.020715668882553055 | FALSE |

| Target Strain | Region Start | Region End | Target_Cov | Ref Strain | Ref Start | Ref End | Alignment Score Norm. | Homolog | Ref Cov | Expression Conserved |
|---|---|---|---|---|---|---|---|---|---|---|
| **5448** | 448351 | 448474 | 0.008190202605153605 | SF370 | 429319 | 429442 | 100.0 | TRUE | 0.016255511663884777 | TRUE |
| **5448** | 1829715 | 1829836 | 0.017334955279314855 | SF370 | 109 | 230 | 100.0 | TRUE | 0.12158330937543402 | FALSE |
| **5448** | 415278 | 415388 | 0.015354788020126466 | SF370 | 396246 | 396356 | 100.0 | TRUE | 0.02114353605087216 | FALSE |

**Supplementary Table 11.** Initial compositions for the development of an RPMI1640-based chemically defined medium. Specification of components and their concentration supplemented to the RPMI4Spy interim media RPMImod version 1 (V1) and RPMImod version 2 (V2). The recipe in the third column of RPMImod V1 resembles the final RPMI4Spy recipe (with 1x Niacinamide and 1x BME vitamin solution).

| Component | RPMImod V1 | | | RPMImod V2 | | |
|---|---|---|---|---|---|---|
| 2 M Glucose | 7 g/L | 7 g/L | 7 g/L | 7 g/L | 7 g/L | 7 g/L |
| 100x Nucleobase Mix (2 g/L A, U and G) | 1x | 1x | 1x | 1x | 1x | 1x |
| 10 mM HEPES Buffer, pH 7.4 | 0.02 mM | 0.02 mM | 0.02 mM | 0.02 mM | 0.02 mM | 0.02 mM |
| 10 mg/mL Niacinamide (1000x) | 1x / 4x | 1x / 4x | 1x / 4x | 1x / 4x | 1x / 4x | 1x / 4x |
| 50x RPMI1640 Amino Acid Solution (AA) | 1x | 1x | 1x | n.a. | n.a. | n.a. |
| 40 mM Amino Acid mix (lab recipe) | n.a. | n.a. | n.a. | 100 µM | 100 µM | 100 µM |
| DL-Methionine | n.a. | n.a. | n.a. | 100 µM | 100 µM | 100 µM |
| 100x RPMI Vitamin Solution | 1x / without | n.a. | n.a. | 1x / without | n.a. | n.a. |
| 100x MEM Vitamin Solution | n.a. | 1x / without | n.a. | n.a. | 1x / without | n.a. |
| 100x BME Vitamin Solution | n.a. | n.a. | 1x / without | n.a. | n.a. | 1x / without |
| RPMI1640 with glutamine, without phenol red | 1x | 1x | 1x | 1x | 1x | 1x |

**Supplementary Table 12.** Nucleotide sequence conservation of the *amyA* locus between different serotypes of *S. pyogenes*. For this analysis, we selected the genomic region of *S. pyogenes* SF370 including the regions of homologous recombination as well as the sequence that is deleted by allelic replacement (coordinates used: start 1,078,523, end: 1,082,749) and performed a multiple sequence alignment. Sequence identity shown in % was analyzed using the integrated MUSLE alignment tool using standard parameters within the geneious software.

| | M28 GAS6180 | M4 GAS10750 | M1 SF370 | M1T1 5448 |
|---|---|---|---|---|
| M28 GAS6180 | | 99.882 | 99.882 | 99.882 |
| M4 GAS10750 | 99.882 | | 99.953 | 99.953 |
| M1 SF370 | 99.882 | 99.953 | | 100.000 |
| M1T1 5448 | 99.882 | 99.953 | 100.000 | |

**Supplementary Table 13.** Nucleotide sequence conservation of the *sagB* locus between different serotypes of *S. pyogenes*. For this analysis, we selected the genomic region of *S. pyogenes* SF370 including the regions of homologous recombination as well as the sequence that is deleted by allelic replacement (coordinates used: start 598,115, end: 599,750) and performed a multiple sequence alignment. Sequence identity shown in % was analyzed using the integrated MUSLE alignment tool using standard parameters within the geneious software.

| | M49 NZ131 | M12 GAS9429 | M28 GAS6180 | M1 SF370 | M1T1 5448 | M4 GAS10750 | M44 STAB901 | M3 GAS315 | M53 Alab49 | M18 GAS8232 | M5 Manfredo | M6 GAS10394 |
|---|---|---|---|---|---|---|---|---|---|---|---|---|
| M49 NZ131 | | 99.450 | 99.511 | 99.511 | 99.511 | 99.511 | 99.328 | 84.674 | 83.136 | 81.591 | 81.924 | 81.980 |
| M12 GAS9429 | 99.450 | | 99.572 | 99.572 | 99.572 | 99.572 | 99.389 | 84.792 | 83.249 | 81.813 | 82.147 | 82.202 |
| M28 GAS6180 | 99.511 | 99.572 | | 100.000 | 100.000 | 99.756 | 99.572 | 84.733 | 83.192 | 81.646 | 81.980 | 82.036 |
| M1 SF370 | 99.511 | 99.572 | 100.000 | | 100.000 | 99.756 | 99.572 | 84.733 | 83.192 | 81.646 | 81.980 | 82.036 |
| M1T1 5448 | 99.511 | 99.572 | 100.000 | 100.000 | | 99.756 | 99.572 | 84.733 | 83.192 | 81.646 | 81.980 | 82.036 |
| M4 GAS10750 | 99.511 | 99.572 | 99.756 | 99.756 | 99.756 | | 99.694 | 84.850 | 83.305 | 81.758 | 82.091 | 82.147 |
| M44 STAB901 | 99.328 | 99.389 | 99.572 | 99.572 | 99.572 | 99.694 | | 84.792 | 83.136 | 81.803 | 81.924 | 81.980 |
| M3 GAS315 | 84.674 | 84.792 | 84.733 | 84.733 | 84.733 | 84.850 | 84.792 | | 93.078 | 92.677 | 92.620 | 92.620 |
| M53 Alab49 | 83.136 | 83.249 | 83.192 | 83.192 | 83.192 | 83.305 | 83.136 | 93.078 | | 98.800 | 99.086 | 99.086 |
| M18 GAS8232 | 81.591 | 81.813 | 81.646 | 81.646 | 81.646 | 81.758 | 81.803 | 92.677 | 98.800 | | 99.268 | 99.155 |
| M5 Manfredo | 81.924 | 82.147 | 81.980 | 81.980 | 81.980 | 82.091 | 81.924 | 92.620 | 99.086 | 99.268 | | 99.887 |
| M6 GAS10394 | 81.980 | 82.202 | 82.036 | 82.036 | 82.036 | 82.147 | 81.980 | 92.620 | 99.086 | 99.155 | 99.887 | |

**Supplementary Table 14.** Nucleotide sequence conservation of the *Spy_1078* locus between different serotypes of *S. pyogenes*. For this analysis, we selected the genomic region of *S. pyogenes* SF370 including the regions of homologous recombination as well as the sequence that is deleted by allelic replacement (coordinates used: start 884,129, end: 886,197) and performed a multiple sequence alignment. Sequence identity shown in % was analyzed using the integrated MUSLE alignment tool, using standard parameters within the geneious software.

| | M44_STAB901 | M12_MGAS9429 | M1 SF370 | M1T1 5448 |
|---|---|---|---|---|
| M44 STAB901 | | 99.228 | 99.275 | 99.227 |
| M12 MGAS9429 | 99.228 | | 99.372 | 99.324 |
| M1 SF370 | 99.275 | 99.372 | | 99.952 |
| M1T1 5448 | 99.227 | 99.324 | 99.952 | |
